# Simulating clinical trials for model-informed precision dosing: Using warfarin treatment as a use case

**DOI:** 10.1101/2023.07.31.551404

**Authors:** David Augustin, Ben Lambert, Martin Robinson, Ken Wang, David Gavaghan

## Abstract

Treatment response variability across patients is a common phenomenon in clinical practice. For many drugs this inter-individual variability does not require much (if any) individualisation of dosing strategies. However, for some drugs, including chemotherapies and some monoclonal antibody treatments, individualisation of dosages are needed to avoid harmful adverse events. Model-informed precision dosing (MIPD) is an emerging approach to guide the individualisation of dosing regimens of otherwise difficult-to-administer drugs. Several MIPD approaches have been suggested to predict dosing strategies, including regression, reinforcement learning (RL) and pharmacokinetic and pharmacodynamic (PKPD) modelling. A unified framework to study the strengths and limitations of these approaches is missing. We develop a framework to simulate clinical MIPD trials, providing a cost and time efficient way to test different MIPD approaches. Central for our framework is a clinical trial model that emulates the complexities in clinical practice that challenge successful treatment individualisation. We demonstrate this framework using warfarin treatment as a use case and investigate three popular MIPD methods: 1. neural network regression; 2. deep RL; and 3. PKPD modelling. We find that the PKPD model individualises warfarin dosing regimens with the highest success rate and the highest efficiency: 75.1% of the individuals display INRs inside the therapeutic range at the end of the simulated trial; and the median time in the therapeutic range (TTR) is 74 %. In comparison, the regression model and the deep RL model have success rates of 47.9% and 65.8 %, and median TTRs of 45 % and 68 %. We also find that the MIPD models can attain different degrees of individualisation: the Regression model individualises dosing regimens up to variability explained by covariates; the Deep RL model and the PKPD model individualise dosing regimens accounting also for additional variation using monitoring data. However, the Deep RL model focusses on control of the treatment response, while the PKPD model uses the data also to further the individualisation of predictions.

## 1 INTRODUCTION

Model-informed precision dosing (MIPD) is an emerging technique used to individualise dosing regimens of otherwise difficult-to-administer drugs (Sheiner, 1969; Keizer et al., 2018). Typical examples that would benefit from MIPD are drugs with narrow therapeutic windows and large treatment response variability, such as warfarin, docetaxel and infliximab (Wadelius and Pirmohamed, 2007; Ma et al., 2021; Gill et al., 2016). Other examples include antibiotics, like vancomycin, where MIPD has been suggested and partially implemented to guide dosing strategies for critically ill patients, balancing the treatment of severe infections and the risks for harmful adverse events (Broeker et al., 2019; Wicha et al., 2021; Matsumoto et al., 2022). MIPD may also be used to efficiently adapt dosing regimens to continuously changing conditions, for example to stabilise the blood glucose levels of diabetes patients with insulin treatment (Wang et al., 2019; Zhu et al., 2020).

The most prominent MIPD methods are regression, reinforcement learning (RL) and pharmacokinetic and pharmacodynamic (PKPD) modelling (Johnson et al., 2011, 2017; Ribba et al., 2020; Keizer et al., 2018). These methods may be further categorised into different subvariants. RL variants include, for example, model-free algorithms such as Q learning (Zadeh et al., 2023) and model-based algorithms such as Monte Carlo tree search (Maier et al., 2021). MIPD variants of PKPD modelling include maximum a posteriori-guided dosing and Bayesian data assimilation-guided dosing (Maier et al., 2020). Although differences across MIPD methods exist, all have in common that they need to process data in two steps in order to make individualised predictions. First models are fitted to population data. This step establishes a relationship between patient characteristics and dosing strategies. In a second step, data specific to the to-be-treated patient is used to predict individualised dosing regimens. For example, regression models have been fitted using records of genetic information and dosages across patients (Gage et al., 2008; IWPC, 2009; Gong et al., 2011), enabling the prediction of individualised dosages based on genetic information.

The data used for model fitting and dosing regimen individualisation differ substantially across MIPD methods and include clinical factors, genetic factors and monitoring data. The type and volume of data are key to both the accuracy of predictions and the ease of implementation in clinical practice (Darwich et al., 2017; Ribba et al., 2022). The more data are collected, the better dosing regimens can be individualised. However, practical constraints limit how much and what type of data may be available for MIPD. As a result, the trade-off between accuracy and practicality needs to be considered when applying MIPD approaches to new medicines. The systematic study of this trade-off for a specific application is, however, complicated by the astronomical costs of clinical trials, rendering repeated clinical trials for different MIPD methods infeasible. In this study, we propose a framework for the simulation of clinical MIPD trials, facilitating a resource-efficient way to test and develop MIPD approaches.

Using simulations to understand MIPD approaches is not a new concept and several MIPD simulation studies exist in the literature. Moore et al. (2004) and Ribba et al. (2022) use simulated treatment responses to investigate RL approaches as a strategy to individualise dosages of anaesthetics in intensive care units. For the simulation, they use a semi-mechanistic PKPD model to emulate the time course of treatment responses and a nonlinear mixed effects (NLME) model structure to capture inter-individual variability (IIV). A similar approach is adopted by Maier et al. (2020, 2021) to study the individualisation of paclitaxel-based chemotherapy using different MIPD approaches, including PKPD modelling, RL and a hybrid PKPD-RL approach. An NLME model simulation approach is also used by Zadeh et al. (2023) to study deep RL as a candidate for MIPD of warfarin. Abrantes et al.(2019) extend this NLME simulation approach, adding inter-occasion variability (IOV) of treatment responses as an extra dimension to the MIPD simulation. They model IOV following Karlsson and Sheiner (1993) and randomly vary the PKPD model parameters of virtual patients over time. An analogous approach is used by Keutzer and Simonsson (2020) to understand MIPD-based treatment individualisation of rifampicin.

We propose an extended framework for the simulation of clinical MIPD trials. Our framework complements the emulation of IIV and IOV by other previously established elements of clinical trial simulation (Holford et al., 2000, 2010). In particular, our framework includes treatment response emergence from complex interactions of pharmacological and physiological processes as a central feature of the trial simulation. In addition, deviations of dose administrations from nominal dosing regimens, as well as delayed monitoring measurements are incorporated in the simulation to more faithfully represent practical limitations of monitoring-based MIPD approaches.

The article is divided into three sections: methods; results & discussion; and conclusion. In the methods, we first introduce the general framework for MIPD trial simulation and subsequently demonstrate its implementation in terms of a clinical trial model for the warfarin use case. We then introduce the three MIPD models, which we will investigate with the help of the clinical trial model. The considered models are: 1. a neural network regression model; 2. a deep reinforcement learning model; and 3. a PKPD model. In the results & discussion, we use the clinical trial model to simulate MIPD trials for each of the three models and analyse the strengths and limitations of the MIPD models. In our analysis, we pay careful attention to attributing generic strengths and limitations to the MIPD methodology and specific strengths and limitations to our implementations of the models. In the conclusion, we summarise our findings and propose future directions for MIPD research.

## 2 METHODS

We first introduce the proposed framework for clinical MIPD trial simulation and discuss the specific implementation for warfarin. We then outline the investigated MIPD methods. The data, models and scripts used in this article are hosted on GitHub (https://github.com/DavAug/mipd-warfarin). A user-friendly API for MIPD approaches using PKPD modelling has been implemented in the open source Python package `chi` (Augustin, 2021). MIPD approaches using neural networks were implemented in `PyTorch` (Paszke et al., 2019).

### 2.1 General framework for MIPD trial simulation

The objective of MIPD is to achieve desired treatment outcomes across individuals despite large treatment response variability by optimising individual dosing regimens. Challenges for this individualisation are nonlinear and delayed treatment responses (Mager, 2006; Dirks and Meibohm, 2010; Véronneau-Veilleux et al., 2020), making it difficult to adjust and extrapolate dosages based on feedback from monitoring. Variability of the treatment response as a result of time-varying pharmacokinetics, concomitant food-intake, comedication and disease progression further complicate the dosing regimen adjustment (Keutzer and Simonsson, 2020). A realistic assessment of MIPD approaches in a simulated trial therefore needs to account not only for IIV, but also for treatment response delays, nonlinearities and IOV. However, PKPD-related aspects are not the only factors influencing the success of MIPD methods. There are also practical limitations for MIPD, including imperfect measurements and deviations from nominal dosing and monitoring schedules (Holford et al., 2000, 2010). While measurement noise is commonly included in simulations, the variability in the execution of the trial often remains neglected. Deviations from the nominal schedule can impact the success of MIPD methods, especially when they are not fed back into the model.

To faithfully emulate the performance of MIPD in clinical practice, our simulation framework accounts for these PKPD-related and practical challenges using a clinical trial (CT) model composed of five components: 1. a mechanistic model; 2. a population model; 3. an inter-occasion model; 4. an execution model and 5. a measurement model (see left panel in Fig 1). We describe each of these components in Section 2.1.2. The right panel of the figure illustrates the workflow of using the CT model for simulating MIPD trials.

**Figure 1.**
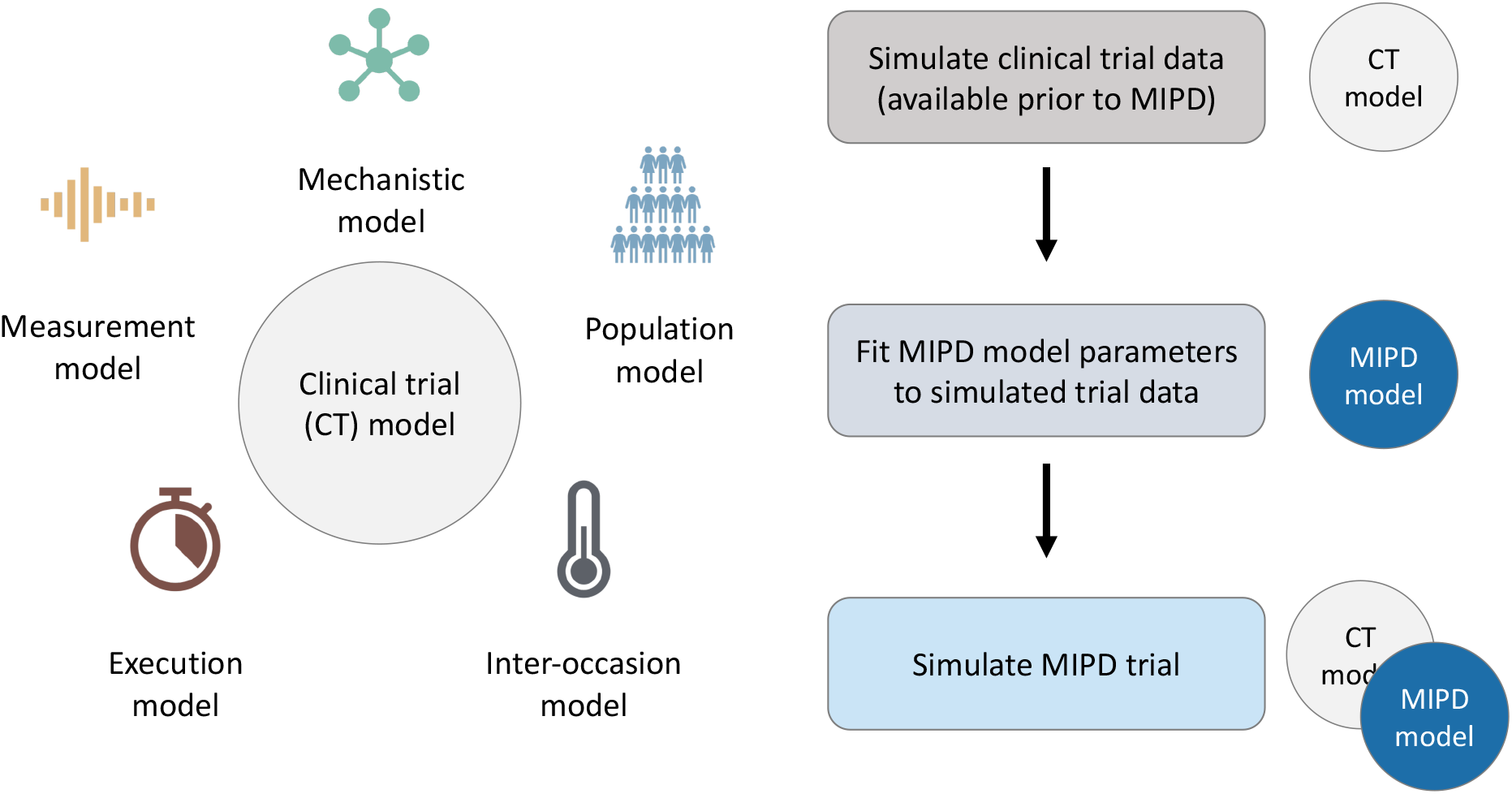
Framework for MIPD trial simulation. The left panel shows the components of the clinical trial (CT) model: 1. a mechanistic model; 2. a population model; 3. an inter-occasion model; 4. an execution model; and 5. a measurement model. The right panel shows the workflow leading up to the MIPD trial simulation. First, the CT model is used to simulate typical clinical trial data, available prior to the MIPD trial. Second, the simulated trial data is used to fit the parameters of the MIPD model. This emulates the starting point of MIPD trials in practice. Finally, the CT model and the fitted MIPD model are used to simulate the MIPD trial.

#### 2.1.1 Workflow of MIPD trial simulation

Our method of MIPD trial simulation involves three steps (see right panel in Fig 1): 1. simulation of pre-MIPD clinical trial data; 2. fitting of the MIPD model to this simulated data; 3. simulation of the MIPD trial. The first two steps emulate the MIPD model development that takes place in practice prior to MIPD trials. In our method, we first simulate typical clinical data from e.g. phase I trials, phase II trials and/or phase III trials using the CT model. We then fit the MIPD model parameters to the simulated data. The details of the fitting process are specific to the MIPD model and are presented in Section 2.3. It is important that the simulated data are used for the fitting, even if real clinical data are available, in order to facilitate a clear attribution of limitations observed in a simulated MIPD trial to the MIPD model. If instead, the MIPD model was fitted to real clinical data, the approximation error of the CT model with respect to the data-generating process of the real clinical trial may also contribute to limitations observed in the simulated trials, making it harder to draw conclusions about the MIPD model. The real data should, instead, be used to calibrate the CT model prior to the data simulation in order to minimise the approximation error as much as possible. The final step of the workflow is the simulation of the MIPD trial using the fitted MIPD model and the CT model.

#### 2.1.2 Components of the CT model

We now describe the components of the CT model. By design, the components of the simulation are non-overlapping and are as modular as possible. This simplifies the model structure and enables an iterative development of CT models, making it possible to replace or further develop individual components without requiring changes in other model components.

##### 1. The mechanistic model

This component models the dynamics of treatment responses as a function of time, *t*, and the dosing regimen, *r*,

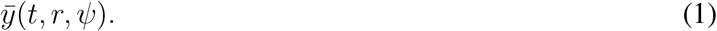

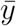 denotes quantities of interest that may be monitored in clinical practice, and *ψ* denotes the parameters of the model. The main purpose of the mechanistic model is to faithfully reflect nonlinearities and delays of the treatment response, emergent from complex cascades of pharmacological and physiological processes. Emulating this complexity provides a tool to test the ability of different MIPD approaches to approximate the treatment response and predict individualised dosing regimens. Popular choices to simulate treatment response dynamics include PKPD models and quantitative systems pharmacology (QSP) models (Ribba et al., 2020; Maier et al., 2021; Azer et al., 2021).

An example mechanistic model of warfarin treatment developed by Wajima et al. (2009) is illustrated in Fig 2. Warfarin is an oral anticoagulant widely used for the prevention and treatment of venous thrombosis, pulmonary embolism and thromboembolic complications associated with atrial fibrillation and/or cardiac valve replacement (FDA, 2010). The left panel shows the 53 blood components described by the model, including warfarin, vitamin K and different coagulation factors, such as thrombin and fibrin. Edges between the components represent interactions, such as transitions, inhibitions or activations. The monitored quantity of the treatment response,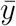, is the prothrombin time. The prothrombin time measures the time it takes for plasma to clot after exposure to a thromboplastin reagent and is routinely measured in clinical practice. In Wajima et al’s model, this prothrombin time test is simulated by measuring the time elapsed between adding the reagent (300 nM tissue factor) and reaching a fibrin area-under-the-curve (AUC) of 1500 nMs (see right panel in Fig 2). The prothrombin time is commonly reported in terms of the international normalised ratio (INR), which measures the prothrombin time of a patient’s blood sample in units of the prothrombin time of a reference sample. We will use this model in our warfarin clinical trial simulation (see Section 2.2 for details).

**Figure 2.**
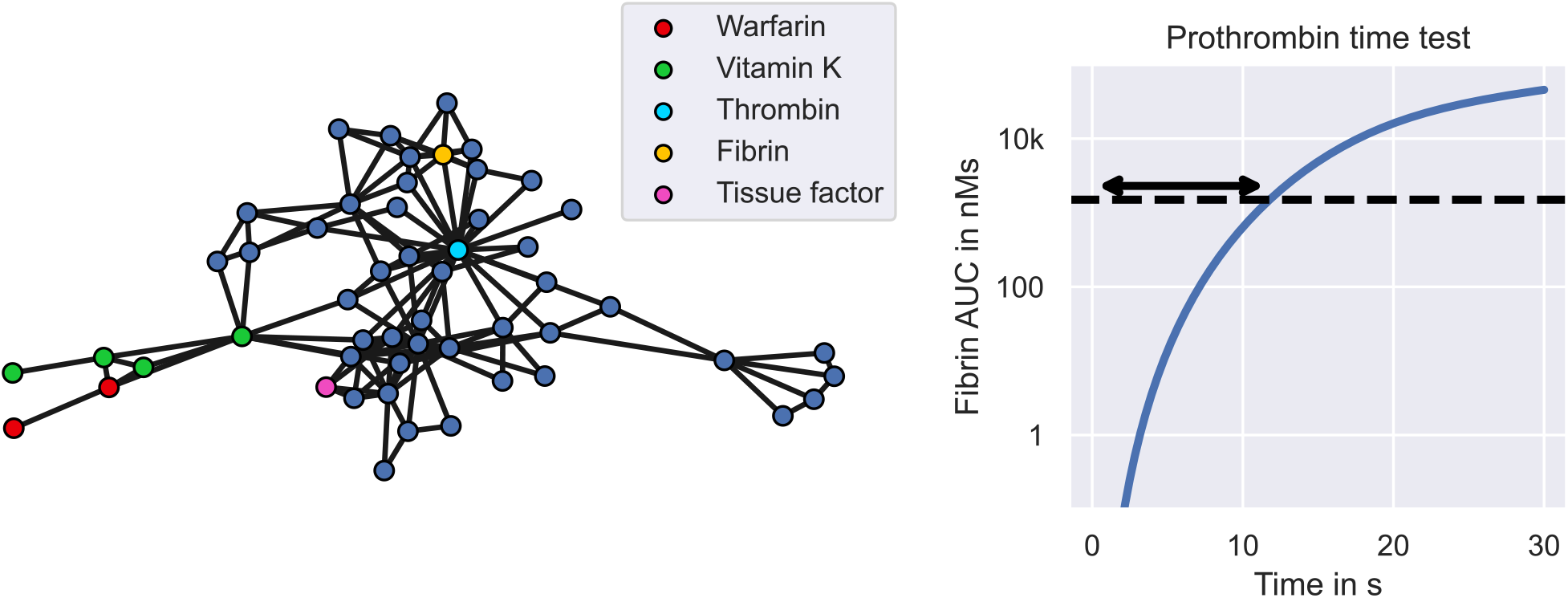
Warfarin clinical trial simulation - the mechanistic model. The figure shows Wajima et al’s model of the warfarin treatment response mechanism. The left panel shows the model. Nodes represent states of the model, including warfarin (red), vitamin **K** (green) and different coagulation factors (blue). Central coagulation factors include thrombin (light blue) and fibrin (yellow). Multiple nodes for warfarin refer to the drug amount in different compartments, while multiple nodes for vitamin **K** also refer to different forms of vitamin **K**, such as vitamin **K** epoxide and vitamin **K** hydroquinone. Interactions between states, such as transitions, inhibitions or activations, are represented by edges. The right panel shows the model simulation of the prothrombin time test. The arrow indicates the prothrombin time which marks the time elapsed between exposure to tissue factor and reaching a fibrin AUC of 1500 nMs.

##### 2. The population model

This component models the variability in the treatment response across individuals using a mixed effects model extension of the mechanistic model. A mixed effects model defines a population distribution of model parameters

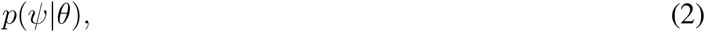

capturing the differences between individuals, i.e. the IIV (Lavielle, 2014; Augustin et al., 2023). *θ* denotes the parameters of the population distribution. Each sample, *ψ*, from the population distribution represents an individual with treatment response 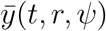. Thus, differences between individuals arise in this model structure from differences in the mechanistic model parameters. For some applications, these differences can be partially explained by covariates, *χ*, which may divide the population into subpopulations, *p*(*ψ*|*θ, χ*). The full population distribution across covariates is then given by the average of the supopulations weighted by the relative frequency of the covariates, *p*(*ψ*|*θ*) = 𝔼_*χ*_ [*p*(*ψ*|*θ, χ*)].

Covariates of the variability can range from clinical factors to genetic factors. For example for warfarin treatment, the VKORC1 genotype explains 27 % of the observed response variability (Wadelius et al., 2009). Other covariates of warfarin treatment include mutations in the CYP2C9 gene and the age of the patient. The link between covariates and pharmacological or physiological processes makes it possible to define mixed effects models that reflect the mechanistic relationship between covariates and the treatment response variability (Hamberg et al., 2007, 2010; Hartmann et al., 2016, 2020). For example, warfarin’s mode of action is the inhibition of the vitamin K epoxide reductase complex (VKORC), and mutations in VKORC’s subunit 1 (VKORC1) affect the inhibitory activity, which can be implemented with a reduced EC50 parameter in Wajima et al’s mechanistic model.

The left panel of Fig 3 illustrates the emergence of inter-individual treatment response variability in our warfarin clinical trial model. The figure shows simulated INR treatment responses of 6 individuals to daily administrations of warfarin. 3 individuals have the GG genotype (VKORC1), the *1/*1 genotype (CYP2C9) and are 71 years old (see blue lines). The remaining 3 individuals have the GA genotype (VKORC1), the *1/*2 genotype (CYP2C9) and are 46 years old (see red lines). The dashed line indicates the treatment response of an average 71 year old individual with the GG and *1/*1 genotypes. We can see that individuals with *χ* = (GG, *1/*1, 71) tend to respond less strongly to warfarin treatment than individuals with *χ* = (GA, *1/*2, 46). However, there remains substantial IIV that is not explained by covariates.

**Figure 3.**
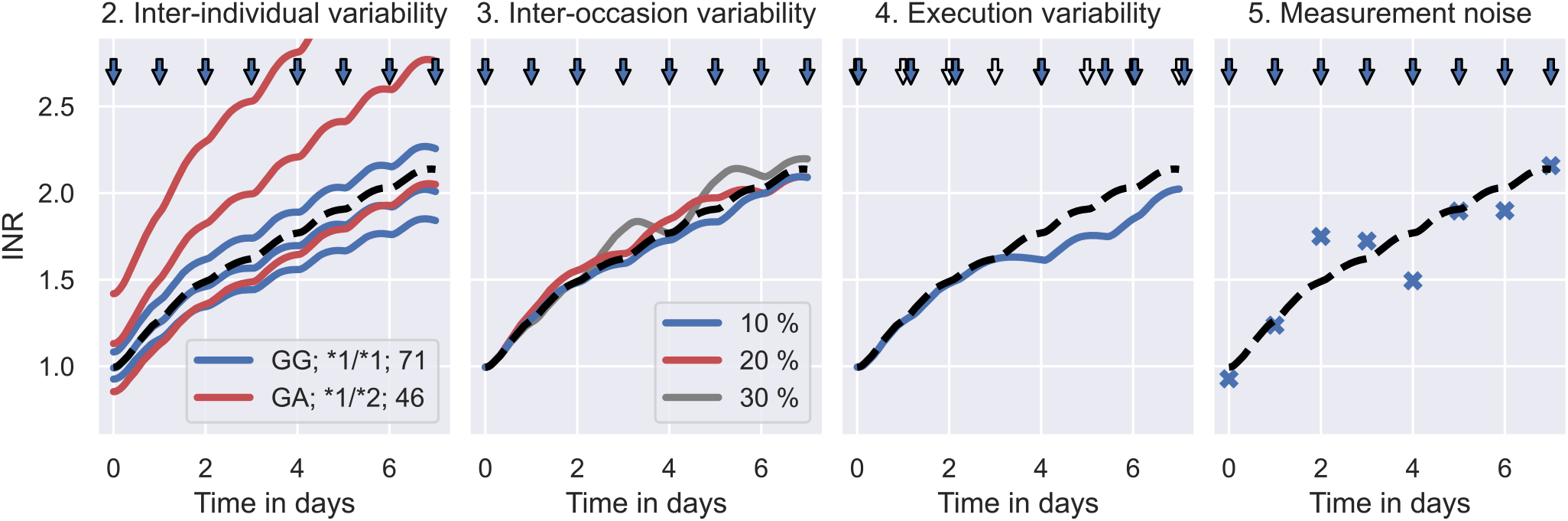
Warfarin clinical trial simulation - sources of treatment response variability. Panel 1: Illustrates the effect of the population model on the clinical trial simulation. The solid lines indicate the treatment responses of 6 simulated individuals to daily warfarin administrations: 3 of which have the covariates *χ* = (GG, *1/*1, 71) (see blue lines); and the remaining 3 have the covariates *χ* = (GA, *1/*2, 46) (see red lines). The typical treatment response across individuals is indicated by a dashed line. The administration times are indicated by blue arrows. Panel 2: Illustrates the effect of the inter-occasion model on the clinical trial simulation. The dashed line shows the treatment response of an individual with a constant vitamin K input rate, i.e. no IOV. The solid lines show the treatment response of the same individual with vitamin K input rates that randomly vary by 10 % (blue), 20 % (red) and 30 % (grey) between days. Panel 3: Illustrates the effect of the execution model on the clinical trial simulation. The dashed line indicates the treatment response of an individual associated with the nominal dosing regimen (hollow arrows). The blue line indicates the treatment response of the same individual associated with the delayed dose administrations (blue arrows). Panel 4: Illustrates the effect of the measurement model on the clinical trial simulation. The dashed line shows the simulated treatment response of an individual and the scatter points show the associated monitoring measurement.

##### 3. The inter-occasion model

This component models the variability of the treatment response over time using time-varying modifications of the model parameters

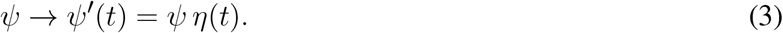

*η* denotes the alterations of the model parameters, and *ψ*′ denotes the new model parameters. The treatment response of an individual with parameters *ψ* is now given by the mechanistic model simulation using the time-varying parameters, 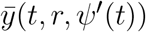 (Karlsson and Sheiner, 1993). The role of the inter-occasion model is to implement changes of the treatment response that are not accounted for by the mechanistic model. Such changes can be externally driven, e.g. by concomitant food intake or comedication, or of completely unknown origin (Keutzer and Simonsson, 2020). The dynamics of *η, p*(*η*|*t*), are a modelling choice.

A source of inter-occasion variability for warfarin is, for example, the time-varying consumption of vitamin K (Xue et al., 2016), changing the amount of vitamin K available in the blood. Since warfarin’s mode of action is to inhibit VKORC – a complex converting one form of vitamin K into another, clotting factor-activating form of vitamin K – an increased availability of vitamin K can reverse the treatment effects of warfarin. In the 2^nd^ panel of Fig 3 we illustrate the effects of varying vitamin K consumption on the warfarin treatment response in the clinical trial simulation. The dashed line shows the treatment response simulation with a constant vitamin K input rate parameter, i.e. no variability in the vitamin K consumption. The solid lines show the treatment response simulations with vitamin K input rates that randomly vary by 10 % (blue), 20 % (red) and 30 % (grey) from day to day.

##### 4. The execution model

This component models unintended deviations from nominal dosing regimens and monitoring schedules during the trial. Nominal dosing regimens are defined by a sequence of doses and administration times, *r* = {(*d*_*j*_, *t*_*j*_)}, where *d*_*j*_ denotes the *j*^*th*^ dose and *t*_*j*_ denotes the associated administration time. Nominal monitoring schedules are similarly defined by a sequence of measurement times. The actual doses and times are modelled in the execution model using random deviations from the nominal schedules

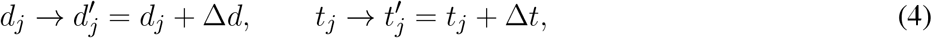

where Δ*d* and Δ*t* denote the deviations. The treatment response corresponding to the actual dosing regimen, 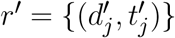, is simulated using 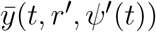. The role of the execution model is to test the robustness of MIPD approaches in clinical practice by emulating the limited control over dose administrations and monitoring. By choosing to report nominal schedules as opposed to actual schedules, the execution model can also be used to mirror common inaccuracies of clinical data. The distributions of dose and time deviations, *p*(Δ*d*) and *p*(Δ*t*), are modelling choices and may differ between dosing and measurement schedules. While not considered in this article, the execution model may be extended to include missed administrations or missed measurements during the trial. Assuming no persistence, this can be implemented using draws of Bernoulli random variables associated with each time point, indicating whether or not a dose was administered or a measurement was taken (Holford et al., 2000, 2010). For infusions, deviations in the duration of the administration may also be modelled.

In the 3^rd^ panel of Fig 3 we illustrate the effects of delayed dose administrations on the treatment response in the warfarin trial simulation. Doses are administered daily. The nominal dosing regimen is illustrated by hollow arrows. The actual dosing regimen with delayed administrations is illustrated by blue arrows. The associated treatment response simulations are indicated by the dashed line (nominal) and by the solid line (actual).

##### 5. The measurement model

This component models the limited accuracy of treatment response measurements

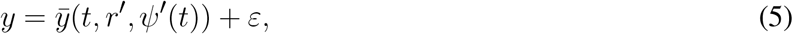

where *y* denotes the measurement and *ε* denotes the measurement error. This defines a distribution of measurements around the mechanistic model output, 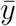, at each time point *t, p*(*y*|*t, r*′, *ψ*′), where measurements may be expected. The wider the measurement distribution, the larger the measurement noise. During the trial simulation, monitoring measurements are sampled from the measurement distribution. This model can be extended to include noise also in the measurement process of covariates, for example to reflect genotyping errors of the VKORC1 or CYP2C9 genes. This is however not considered in this article.

In the 4^th^ panel of Fig 3, we illustrate INR measurements sampled from the measurement distribution during warfarin treatment (blue scatter points). The dashed line shows the treatment response simulation of the mechanistic model without measurement noise.

### 2.2 Implementation of warfarin trial simulation

We develop the warfarin clinical trial model following the framework for MIPD trial simulation introduced in Section 2.1. The mechanistic model is implemented using Wajima et al’s model of the humoral coagulation network (see Fig 2). The model provides a mechanistic description of warfarin’s PKPD using a system of nonlinear differential equations

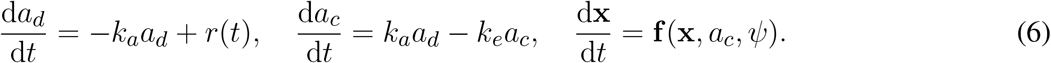

*a*_*d*_ and *a*_*c*_ describe the pharmacokinetics of warfarin and denote the amount of the drug in the dose compartment and the central compartment, respectively. *k*_*a*_ denotes the absorption rate and *k*_*e*_ denotes the elimination rate. *r*(*t*) denotes the dose rate and implements the dosing regimen. The pharmacodynamics of warfarin are captured by **x**, denoting the 51 remaining states of the model. The prothrombin time is simulated as a function of the states, 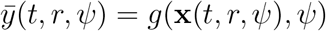, involving the computation of the fibrin AUC after exposure to 300 nM tissue factor. For full details of the mechanistic model, we refer to (Wajima et al., 2009) and Appendix S1. A systems biology markup language (SBML) specification is provided on GitHub (https://github.com/DavAug/mipd-warfarin) for simplified cross-platform implementation of the model.

The population model is implemented using a hybrid of two mixed effects models developed by (Hartmann et al., 2016, 2020) and (Hamberg et al., 2010). Hartmann et al (2016) provide a mixed effects model extension of Wajima et al’s model, capturing the treatment response variability emergent from varying production rates of selected coagulation factors, including prothrombin, protein S, protein C and coagulation factors V, VII, IX, X, XI, XII and XIII. In a subsequent publication, they extend their model to include variability explained by covariates, such as the genotypes of the VKORC1 gene and the CYP2C9 gene (Hartmann et al., 2020). VKORC1 is used to model the variability in warfarin’s EC50, while CYP2C9 is used to model the variability in warfarin’s elimination rate. In our population model, we modify Hartmann et al’s model further using elements from Hamberg et al’s model to incorporate age and heterozygosity in the genotypes as covariates of the IIV (Hamberg et al., 2010). This results in a population distribution whose subpopulations, *p*(*ψ*|*θ, χ*), are defined by the age of a patient, one of three VKORC1 genotypes (GG; GA; AA) and one of six CYP2C9 genotypes (*1*1; *1*2; *1*3; *2*2; *2*3; *3*3). For full details of the population model, we refer to Appendix S1. The population model parameters, *θ*, used to simulate the clinical trial are provided on GitHub (https://github.com/DavAug/mipd-warfarin).

The inter-occasion model is implemented using time-varying vitamin K input rates (see 2^nd^ panel in Fig 3). The input rate alterations are assumed to be normally distributed

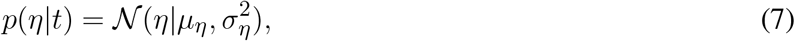

with constant mean and standard deviation: *μ*_*η*_ = 1 and *σ*_*η*_ = 0.1. To allow a change of vitamin K consumption over time, we sample a new *η* for each simulation day, such that the altered input rate may be interpreted as the daily average of the vitamin K consumption.

The execution model is implemented using exponentially distributed delays of the administration and monitoring times

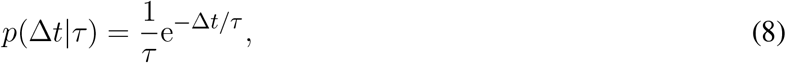

where *τ* = 30 min denotes the average delay. For each nominal administration time and each nominal monitoring time, we independently sample delays from *p*(Δ*t*|*τ*) and compute the actual times according to Eq 4. If dose administrations and monitoring measurements are scheduled at the same nominal times, we only draw one delay random variable for both events.

The measurement model is implemented using lognormally distributed measurements around the mechanistic model output

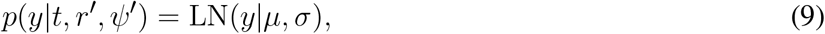

where 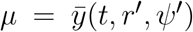 denotes the median, and *σ* = 0.1 denotes the scale of the distribution. This implements a measurement error that scales proportionally with the measured quantity, producing measurement errors of approximately 10 % of the mechanistic model output.

### 2.3 MIPD methods

In this article, we investigate three MIPD models for warfarin treatment individualisation: 1. a neural network regression model (‘Regression model’); 2. a deep reinforcement learning model (‘Deep RL model’); and 3. a pharmacokinetic and pharmacodynamic model (‘PKPD model’). All three models use data specific to the to-be-treated patient to predict individualised dosing regimens. While the details differ substantially between the models, Fig 4 illustrates their general workflow. The bottom of the figure shows the to-be-treated patient characterised by covariates, such as clinical and genetic factors. The top of the figure shows the treating physician and the MIPD model. The physician uses the model and the patient characteristics to predict an individualised dose. For some MIPD approaches, this dose can be iteratively refined using measurements of the patient’s treatment response. Some models can also predict the full future dosing regimen at each iteration rather than just the next dose. Below, we discuss the three methods investigated in this study in more detail.

**Figure 4.**
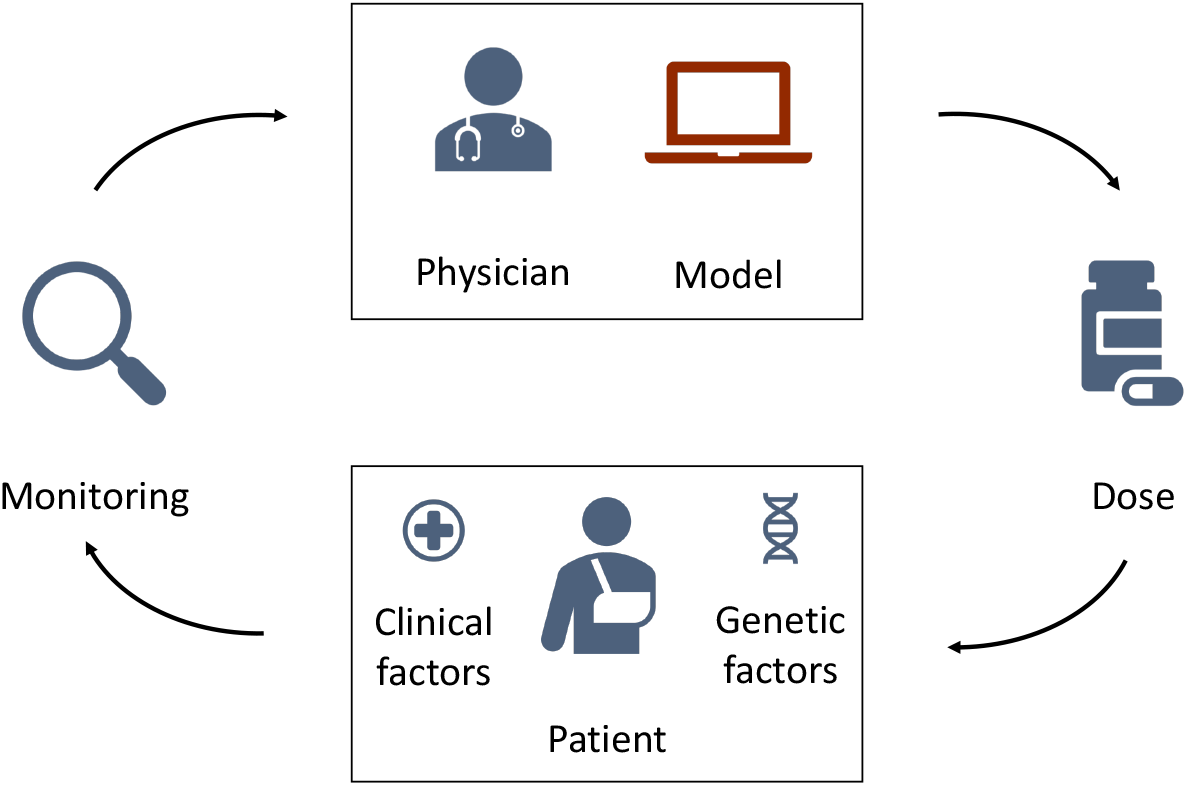
Schematic illustration of model-informed precision dosing. The top of the figure shows the treating physician and the MIPD model. The bottom of the figure shows the to-be-treated patient characterised by clinical and genetic factors. The physician uses the MIPD model and the patient characteristics to predict an individualised dose. If the treatment response is monitored over time, the physician can use the treatment response measurements and the MIPD model to iteratively refine the dose predictions.

#### 1. Regression model

This MIPD approach follows Anderson et al. (2012) and Verhoef et al. (2013) and uses a static model of the daily maintenance dose to individualise treatments. The maintenance dose refers to the constant warfarin dose administered daily to maintain a desired INR level. The maintenance dose is modelled as a function of the desired response, *y*\*, and the patient’s covariates, *χ*,

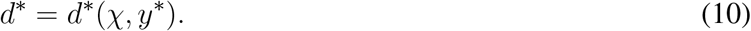

*d*\* denotes the maintenance dose. This makes it possible to predict individualised maintenance doses based on the covariates of an individual. In contrast to the generalised workflow in Fig 4, the model does not iteratively update its predictions using monitoring measurements.

The model can be implemented using a variety of regression approaches, including linear regression, spline regression and tree regression (IWPC, 2009). In this article, we choose a neural network approach. Neural networks are universal function approximators and can therefore learn to approximate the maintenance dose function in Eq 10 from data, even when the relationship between the dose and (*χ, y*\*) is nonlinear.

For those who are more familiar with neural network regression, in summary we compose the network of three sequential fully connected layers implemented in `PyTorch` (Paszke et al., 2019); the two inner layers of this network are of width 1024 and have ReLU activations. The output layer uses a sigmoid activation. The network is trained on simulated trial data (see Section 3.2) to minimise the mean squared error objective function using Adam (Kingma and Ba, 2014). For full details on the implementation and training of the model, we refer to Appendix S2.

#### 2. Deep RL model

This MIPD approach follows a deep reinforcement learning approach similar to (Zadeh et al., 2023) and uses a model of the next-to-administer dose to individualise treatments. The dose is modelled as a function of the covariates and the current monitoring data

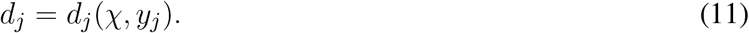

*d*_*j*_ denotes the dose at time *t*_*j*_ and *y*_*j*_ denotes the INR measurement at time *t*_*j*_. This makes it possible to iteratively predict individualised dosages based on monitoring data and the covariates of an individual, as illustrated in Fig 4. The target treatment response, *y*\*, is specified before the training of the model (Zadeh et al., 2023).

Conceptually, RL learns dosing strategies from trial and error: the model sequentially administers dosages and evaluates the ‘goodness’ of the dose decisions based on the feedback from the treatment response. The learned dosing strategy can be shown to optimally target the desired treatment response under certain technical assumptions and takes the form of a function for the next-to-administer dose (see Eq 11). We discuss the limitations of these assumptions in Section 3.5. While trial and error in a clinical setting raises ethical questions, RL models can also be trained on treatment response emulators (Ribba et al., 2022). Popular treatment response emulators are PKPD models (Zadeh et al., 2023). To reduce the number of trial and error iterations needed for convergence of the dosing strategy, RL can be performed in conjunction with function approximators (Baird, 1995). In this article, we choose a deep neural network as the function approximator – an approach commonly referred to as deep RL.

For those who are more familiar with deep RL: we use a DQN model to implement the Deep RL model Mnih et al. (2013). We compose the network of four sequential fully connected layers implemented in `PyTorch` (Paszke et al., 2019): three hidden layers of widths (256, 128, 64) with ReLU activations, and the output layer of width 58. The outputs of the network are the predicted Q-values Mnih et al., 2013). The dose with the maximum Q-value is suggested for administration. Following (Zadeh et al., 2023), the network is trained online using a PKPD model as a treatment response emulator. Prior to the training, the PKPD model is fitted to simulated trial data (see Section 3.2). We train the model to minimise the temporal difference error using Double Q-learning (Van Hasselt et al., 2016) and the Adam optimiser (Kingma and Ba, 2014). For full details on the implementation and training of the model, we refer to Appendix S3.

#### 3. PKPD model

This MIPD approach follows a PKPD modelling approach similar to Hamberg et al. (2015) and uses a model of the dosing regimen to individualise treatments. The dosing regimen is modelled as a function of the covariates, the desired treatment response and the monitoring data

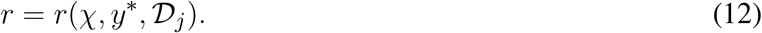

𝒟_*j*_ = {(*y*_1_, *t*_1_),…, (*y*_*j*_, *t*_*j*_), *r*_*j*_} denotes the monitoring data available up to and including time *t*_*j*_, where we use r_j_ to denote the administered dosing regimen for *t* < *t*_*j*_. The function makes it possible to predict individualised dosing regimens based on monitoring data and the covariates of an individual (see Fig 4). The predictions can be iteratively refined as more monitoring data becomes available.

PKPD modelling is a semi-mechanistic modelling approach which makes use of approximate descriptions of the physiological and pharmacological processes to predict the treatment response dynamics. Conceptually, PKPD modelling is similar to the mechanistic model component of the CT model (see e.g. Wajima et al’s model in Fig 2), but generally model the biological processes in lower detail. An example PKPD model is illustrated in Fig S4.7 in Appendix S4. By comparing the predicted and desired treatment responses, this approach is able to determine the optimal dosing regimens for each individual. Analogously to the CT model in Section 2.1.2, inter-individual variability is described by differences in the model parameters across individuals. Patient-specific parameters are derived from the patient’s covariates and monitoring data.

For those who are more familiar with PKPD modelling, in summary we use Hamberg et al’s model (Hamberg et al., 2010) implemented in `chi` (Augustin, 2021) to model the warfarin treatment response dynamics. We fit the model to simulated trial data (see Section 3.2) using hierarchical Bayesian inference and the No-U-Turn sampler (NUTS) (Hoffman et al., 2014) implemented in `pints` (Clerx et al., 2019). Individualised dosing regimens are predicted in two steps: 1. the model is fit to an individual’s monitoring data using the population model and the individual’s covariates as prior knowledge (Maier et al., 2020); and 2. the dosing regimen is optimised to minimise the mean squared error between the model predictions and the desired treatment response. We use Bayesian inference and `pints`’ implementation of the adaptive covariance matrix Markov chain Monte Carlo (ACMC) sampler to fit the model to an individual’s data. For the dosing regimen optimisation, we use `pints`’ implementation of the covariate matrix adaption evolution strategy (CMA-ES) optimiser (Hansen et al., 2003). For full details on the implementation and the dosing regimen prediction, we refer to Appendix S4.

## 3 RESULTS & DISCUSSION

We simulate three MIPD trials – one trial for each MIPD model – and analyse their relative strengths and limitations. To this end, we follow the workflow illustrated in Fig 1 and, first, fit the MIPD models to simulated clinical trial data to emulate a typical starting point for MIPD trials. The data imitate typical data collected during each of the three phases of clinical trials, and are simulated using the CT model. After the model fitting, we simulate the MIPD trials.

### 3.1 Simulating trial phases prior to MIPD

At present, most clinical trials are conducted not having model-informed precision dosing in mind. As a result, the data available for the fitting of MIPD models will often not be tailored to the needs of the method. To reflect this practical limitation in our study, we simulate data from three typical phases of clinical trials for the warfarin use case, not taking the data-requirements for MIPD into account. All MIPD models in this article are fitted using only the data from these trials. The simulated trial data as well as the code to reproduce the trials are hosted on GitHub (https://github.com/DavAug/mipd-warfarin).

#### Trial phase I

Phase I trials are primarily used to establish the safety of drugs and often monitor a drug’s absorption, distribution, metabolism and elimination (ADME) in a relatively small cohort of patients. We emulate such a trial by mirroring a clinical trial reported in (Hamberg et al., 2007). In this trial, the pharmacokinetics of N = 60 individuals is monitored. Each individual is administered with a single 10 mg dose of warfarin. Following the nominal administration time, the warfarin concentration in the blood is measured at 10 h, 35 h and 60 h after the administration. The data collected during the trial are the warfarin concentration measurements, the dosing regimens and the covariates for each of the 60 individuals. The simulated warfarin concentrations are illustrated in the left panel of Fig 5. The demographics of the cohort are reported in Table S1. Pseudo-code outlining the implementation of the trial is presented in Algorithm S1.

**Figure 5.**
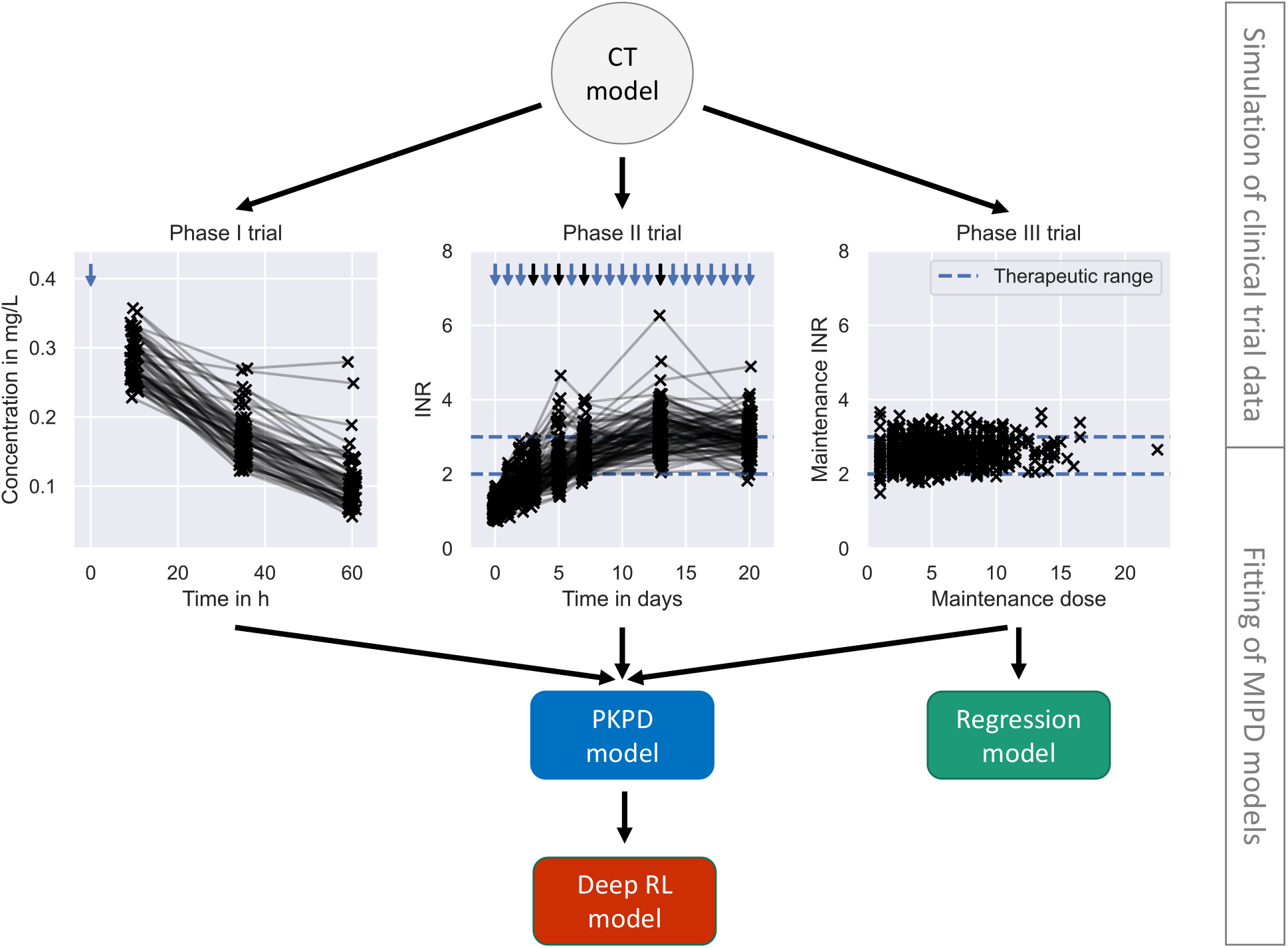
Pre-MIPD trial workflow. The figure shows the two steps performed prior to the MIPD trial simulation: 1. Simulation of clinical trial data (top); and 2. fitting of MIPD models (bottom). The CT model is used to simulate the trial data. The simulated data is illustrated in the middle of the figure: phase I (left); phase II (middle); phase III (right). Measurements of the blood warfarin concentration or the INR are illustrated using scatter points. Measurements connected by solid lines are taken from the same individual at different time points. Nominal administration times are illustrated by blue arrows. Dose administrations with individualised, on-the-fly adjustments of the dose amount are highlighted in black. The therapeutic range is illustrated in the middle and right panel using blue dashed lines. The bottom of the figure shows the fitting workflow of the MIPD models: The PKPD model is fitted to the data from all trials; the Regression model is fitted only to the data from clinical trial phase III; and the Deep RL model is fitted to treatment responses simulated with the PKPD model.

The panel shows the simulated warfarin concentration measurements. Measurements are illustrated using scatter points. Measurements taken from the same individual are connected using a solid line. The nominal administration time of the warfarin dose is indicated using a blue arrow.

#### Trial phase II

Phase II trials are primarily used to establish the efficacy of drugs and monitor a drug’s pharmacodynamics. We simulate a phase II trial using a design similar to trials reported in (Hamberg et al., 2010). We monitor the INR response of 100 simulated individuals for 3 weeks. Each individual is treated with daily warfarin doses. During the first 3 days, all individuals receive the same treatment: *d*_1_ = 10 mg; *d*_2_ = 7.5 mg; and *d*_3_ = 5 mg. The 4th dose is adjusted for each individual based on the INR treatment response by a medical professional, here emulated using a simple linear heuristic: *d*_*j*_ = *d*_*j*−1_ *y*\*/*y*_*j*_, where *y*_*j*_ denotes the INR measurement taken just before the *j*^*th*^ dose administration. This heuristic computes a personalised warfarin dose targeting the desired treatment response, *y*\*, assuming a linear relationship between the INR and the dose amount. The dose amounts are adjusted three more times for each individual on day 5, 7 and 13 of the trial using the same heuristic. The INR of the individuals is closely monitored during the induction phase of the trial (day 0, day 1, day 2, day 3) and less frequently measured as the trial progresses (day 5, day 7, day 13 and day 20). The trial is discontinued for individuals with INRs 10 to emulate safety constrains of real clinical trials. The data collected during the trial are the INR measurements, the nominal dosing regimens and the covariates for each of the 100 individuals.

The trial was not terminated early for any of the simulated individuals. The simulated INR measurements are illustrated in the middle panel of Fig 5. The demographics of the cohort are reported in Table S1. Pseudo-code outlining the implementation of the trial is presented in Algorithm S2. The panel shows the simulated INR measurements. Measurements are illustrated using scatter points. Measurements taken from the same individual are connected using a solid line. The nominal administration times of the warfarin doses are indicated using arrows. Black arrows indicate personalised adjustments of the dose amount. The therapuetic range is indicated using blue dashed lines.

#### Trial phase III

Phase III trials can vary in scope, but tend to involve larger cohorts and have a stronger focus on treatment endpoints. We emulate a phase III trial by mirroring a clinical trial reported in (IWPC, 2009). The focus of the trial is to understand the variability of the maintenance warfarin dose. We simulate the trial analogously to trial phase II, but with a larger cohort, N = 1000, and for a longer duration (8 weeks). Following the initial, identical phase of the trial, the daily doses are adjusted two more times on day 27 and 34. During the final three weeks of the trial, the dose amounts remain unchanged to guarantee that individuals equilibrate to the maintenance treatment response by the end of the trial. The trial is discontinued for individuals with INRs ≥ 10 to emulate safety constraints of real clinical trials. The data collected during the trial are the INR measurements at the end of the trial (day 55), the maintenance warfarin doses and the covariates for each of the 1000 individuals.

The trial was not terminated early for any of the simulated individuals. The simulated maintenance doses and INR measurements are illustrated in the right panel of Fig 5. The demographics of the cohort are reported in Table S1. Pseudo-code outlining the implementation of the trial is presented in Algorithm S3. The panel shows the maintenance dose on the x-axis and the measurement of the INR on the y-axis. Measurements are illustrated using scatter points. The therapuetic range is indicated using blue dashed lines.

### 3.2 Fitting the MIPD models

We fit the MIPD models to the simulated clinical trial data, as illustrated in Fig 5. The fitting is the second step of the MIPD trial simulation workflow (see Fig 1). Only one of the models – the PKPD model – can be fitted to all of the available clinical trial data. The Regression model is fitted using the data from the phase III trial. The Deep RL model is trained indirectly on the trial data through simulations from the fitted PKPD model. For details on the fitting, we refer to Section 2.3, Appendix S2, Appendix S3 and Appendix S4.

### 3.3 Simulating the MIPD trials

We simulate one MIPD trial for each of the three methods in Section 2.3. All trials are conducted using the same cohort. The cohort includes N = 1000 individuals. The demographics of the cohort are modelled after a trial reported in (Hamberg et al., 2010) and are visualised in Fig 6. The figure shows the CYP2C9 genotype distribution, the VKORC1 genotype distribution and the age distribution in the cohort.

**Figure 6.**
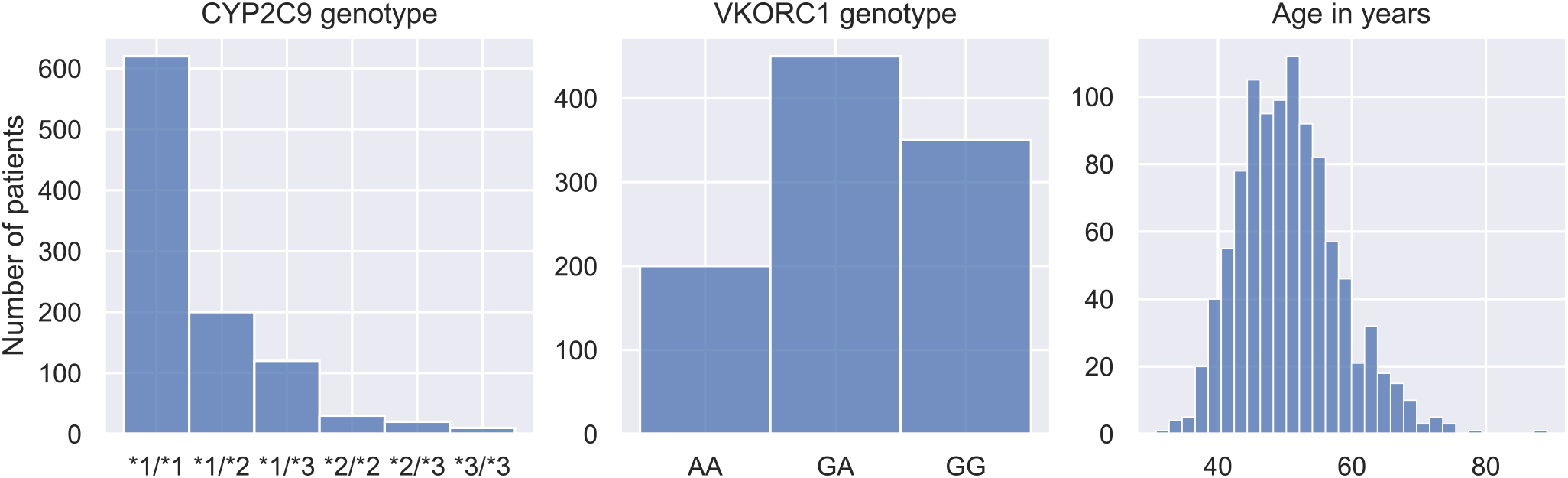
Demographics of MIPD trial cohort. The figure shows the CYP2C9 genotype distribution (left panel), the VKORC1 genotype distribution (middle panel) and the age distribution (right panel) of the MIPD trial cohort. The cohort includes 1000 simulated individuals.

In each trial, individuals are treated with daily warfarin doses for 19 days. The dose amounts are individualised using the respective MIPD model, as described in Section 2.3, and target a treatment response of *y* * = 2.5. The data available for the individualisation are the INR measurements taken daily before each dose administration and the covariates of the individual. The Regression model individualises the treatments by predicting the maintenance dose based on the covariates of the patients. This maintenance dose is administered every day throughout the trial. The Deep RL model and the PKPD model predict the warfarin doses iteratively based on a patient’s covariates and INR measurements. Pseudo-code outlining the implementation of the trial is presented in Algorithm S4.

### 3.4 Results of the MIPD trials

We use three metrics to quantify the success, the safety, and the efficiency of the dosing regimen individualisation: the maintenance INR; the peak INR; and the time in the therapeutic range (TTR). The success of the individualisation is quantified using the maintenance INR measured on the last day of the trial. INR measurements inside the therapeutic range indicate success, while measurements outside the therapeutic range indicate poor dosing regimen individualisation. The safety of the individualisation is quantified using the largest INR measurement recorded during the trial. This peak of the INR response indicates the risk for major bleeding events while transitioning into maintenance treatment. The efficiency of the individualisation is quantified using the TTR, i.e. the number of INR measurements inside the therapeutic range. For successful individualisations, the TTR indicates how quickly the desired treatment response has been achieved.

The results of the trials are visualised in Fig 7. Row 1 shows the results for the Regression model, row 2 shows the results for the Deep RL model, and row 3 shows the results for the PKPD model. The left panel shows the maintenance INR distribution in the cohort. INRs inside the therapeutic range (see blue dashed lines) are highlighted in blue and INRs outside the therapeutic range are highlighted in red. The target INR is illustrated using a black dashed line. The middle panel shows the peak INR distribution in the cohort. The right panel shows the TTR distribution across individuals. The target TTR is illustrated using a dashed line.

**Figure 7.**
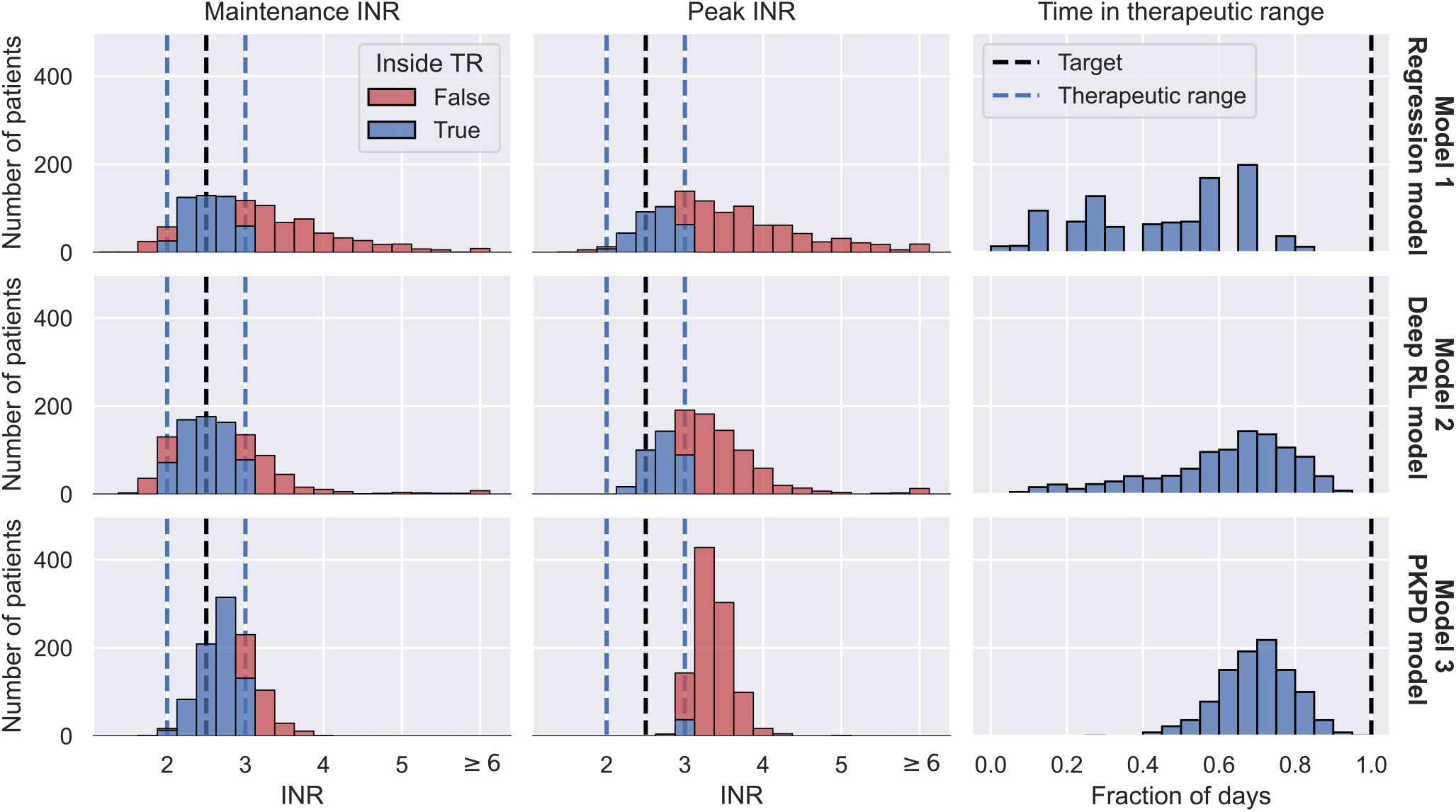
MIPD trial results. The figure shows the outcome of three MIPD trials conducted with an identical cohort of size N = 1000 with different MIPD models. The top row shows the results for the Regression model; the middle row shows the results for the Deep RL model; and the bottom row shows the results for the PKPD model. The outcome of the trials is illustrated using three metrics: the maintenance INR measured on the last day of the trial (left panel); the largest INR value measured during the trial (middle panel); and the time in the therapeutic range (right panel). The panels show the distributions of these metrics across individuals. The therapeutic range is illustrated using blue dashed lines. INR measurements inside the therapeutic range are highlighted in blue, and INR measurements outside the therapeutic range are highlighted in red. Target values of the dosing regimen individualisation are visualised using black dashed lines.

The maintenance INR distributions (left panel) show that in a time span of 19 days all three MIPD methods are able to successfully target the therapeutic window for a large number of individuals. The Regression model successfully individualises the dosing regimen for 47.9% of the individuals, while the Deep RL model and the PKPD model have success rates of 65.8% and 75.1 %, respectively (see blue histograms). For the remaining individuals, the severity of the failed dosing regimen individualisation varies substantially. For the Regression model, 149 individuals display maintenance INRs above 4 with the largest value being 7.51. For the Deep RL model, only 32 individuals exceed maintenance INRs of 4. However, the largest maintenance INR is 29.03 – a value almost 4 times larger than for the Regression model, raising serious safety concerns. For the PKPD model the largest maintenance INR is 3.90. 97.8% of the individuals display maintenance INR measurements less than 3.5, showing that the PKPD model is able to most consistently achieve maintenance INRs close to the therapeutic window.

The peak INR distributions (middle panel) show that, for the majority of the cohort, the methods are not able to individualise the dosing regimens without overshooting the therapeutic window. In the Regression model trial, 68.9% of the individuals display peak INR measurements above the therapeutic range. The Deep RL model controls the treatment response marginally better, overshooting the therapeutic range for 65.1% of the individuals. The PKPD model misses the therapeutic window for almost all individuals (95.9 %) before reaching maintenance treatment. However, the panel also shows that the largest INR value measured in the PKPD model trial is substantially smaller than the largest INR values in the other two trials (Regression model: 7.99; Deep RL model: 29.03; PKPD model: 4.91), indicating that the PKPD model is the safest MIPD approach among the tested models.

The TTR distributions (right panel) show that the time spent inside the therapeutic range varies between individuals and MIPD approaches. For the Regression model, the median TTR is 45 % of the trial duration, with TTRs ranging between 0% and 85 %. The other two methods achieve substantially larger TTRs across individuals. For the Deep RL model, the median TTR is 68 % of the trial duration, with individual values ranging between 5% and 95 %. The PKPD model achieves a median TTR of 74 %, with a minimum TTR of 26 % and a maximum TTR of 100 %. This shows that across individuals the PKPD model takes the least time to successfully reach the therapeutic window.

### 3.5 Degrees of dosing regimen individualisation

The simulated trials in Section 3.4 show that different MIPD approaches have different strengths and limitations. In this section, we study the dosing strategies of the models in more detail to gain a better understanding about their practical and methodological differences. We pay particular attention to attributing generic strengths and limitations to the methodology and specific strengths and limitations to our implementation.

We investigate the dosing strategies by studying the dose decisions suggested by each of the models for three representative individuals from the simulated trials in Section 3.4. The first individual, with ID 716, is characterised by the covariates *χ*_*A*_ = (*1*1, GG, 50). The other two individuals (ID 269; ID 305) are both characterised by the covariates, *χ*_*B*_ = (*1*2, GA, 50). The different dosing strategies and treatment responses are visualised in Fig 8. The figure shows the doses administered during the trials in the top panel and the corresponding treatment response measurements in the bottom panel. Doses or measurements belonging to the same individual are connected using solid lines. The therapeutic range is illustrated using dashed lines. The left panel shows the trial results for the Regression model, the middle panel shows the trial results for the Deep RL model, and the right panel shows the trial results for the PKPD model.

**Figure 8.**
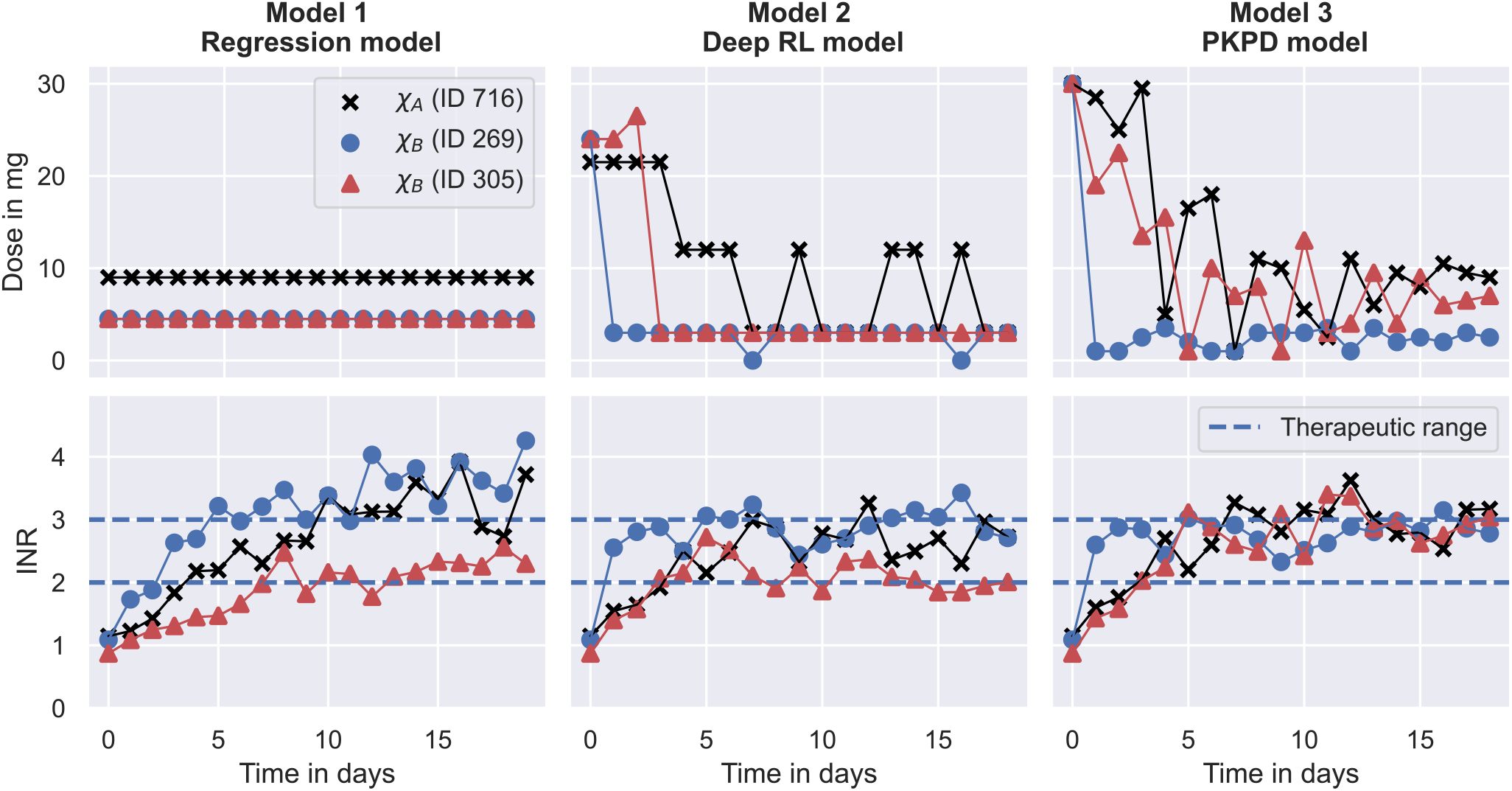
Degrees of dosing regimen individualisation. The figure shows the dosing regimen individualisations achieved by the MIPD models in the simulated MIPD trials for three representative individuals. The left panel shows the results for the Regression model, the middle panel shows the results for the Deep RL model, and the right panel shows the results for the PKPD model. The top row illustrates the administered dose amounts and the bottom panel illustrates the INR monitoring data. Dose amounts or measurements belonging to the same individual are connected using solid lines. The therapeutic range is illustrated using dashed lines. One of the individuals (ID 716) is characterised by the covariates *χ*_*A*_ = (*1*1, GG, 50). The other two individuals (ID 269; ID 305) are both characterised by the covariates *χ*_*B*_ = (*1*2, GA, 50).

#### 3.5.1 The Regression model

The left panel of the figure shows that the Regression model predicts a maintenance dose of 9 mg for the individual with ID 716 (black scatter points), and a maintenance dose of 4.5 mg for the other two individuals (see top left panel in Fig 8). These maintenance doses are administered daily throughout the trial. For the individual with ID 305, the maintenance INR measurement at the end of the trial is inside the therapeutic range, while the maintenance INR measurements for the other two individuals overshoot the therapeutic range (see bottom left panel in Fig 8). Notably, the individual with the successful individualisation (ID 305) has the same covariates as one of the individuals with the failed individualisation (ID 269).

This illustrates both a strength and a limitation of the Regression model: a strength of the model is that it is able to predict individualised dosages using information only about the covariates of individuals, making the model easier to implement in clinical practice than monitoring-based approaches. However, the figure also shows that dosing regimens exclusively derived from covariates will, at best, successfully target the desired treatment response for an average individual characterised by the covariates. Inter-individual differences that are not explained by covariates are not accounted for. In this case, the individual with ID 305 happens to respond similarly to an average individual^1^ with the covariates, x_B_, resulting in a successful dosing regimen individualisation. The individual with ID 269, on the other hand, responds more strongly to warfarin than an average individual^2^, yielding a maintenance INR above the therapeutic range. Not being represented by an average individual also explains the failed individualisation for the individual with ID 716. The inability to account for unexplained IIV is a generic limitation of approaches exclusively based on covariates.

In addition to this limited ability to account for IIV, the figure also shows that the Regression model is incapable of accounting for inter-occasional variation and differences in the execution of the treatment. This limitation is, again, a direct consequence of exclusively using covariates for the dosing regimen individualisation. Without quantifying the individual-specific IOV and EV, predicted dosing regimens can, at best, be successful when the IOV and the EV of the treated individual happens to be close to the average IOV and EV observed in the dataset used for the model fitting.

Another limitation, illustrated in Fig 8, is that the Regression model cannot account for treatment response delays. The model only predicts maintenance dosages, and therefore provides no guidance for individualisation of the induction phase of the treatment. In our implementation, we choose to overcome this limitation of the model by administering the predicted maintenance dosages from the beginning of the trial, not attempting to individualise induction dosing regimens without model guidance. This has the consequence that treatment responses take time to reach their maintenance level, limiting the efficiency attainable by the model. From Fig 8, we can, for example, see that the INR measurements of the individual with ID 305 reach the therapeutic range for the first time on day 8 of the treatment. After that, two more measurements are outside the therapeutic range, giving rise to a TTR of 10 days during the 19 day trial. Together with the top right panel in Fig 7, this indicates that even when the Regression model individualises maintenance dosages successfully, it achieves, at best, an average TTR of around 55 %-65 %. This limit to the efficiency is specific to the treatment response delay of warfarin and our implementation of the Regression model.

A summary of the strengths and limitations specific to the Regression model are presented in the left column of Table 1.

**Table 1.**
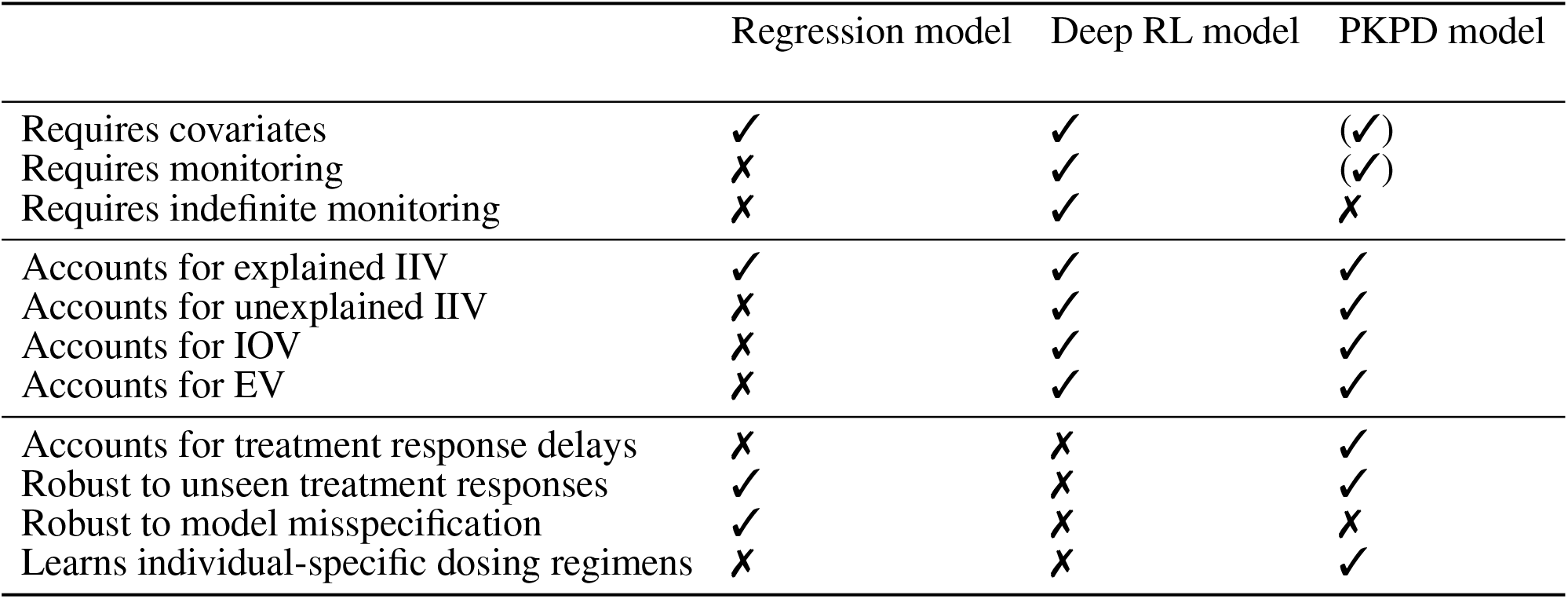
Summary of the strengths and limitations of the MIPD models. The properties of the models are grouped into data-related properties (rows 1-3), variability-related properties (rows 4-8), and strategy-related properties (rows 9-13). The parentheses around the property of the PKPD model in the top right corner indicate that the model, as defined in Section 2.3, can be used without covariates, but, in the simulated MIPD trial in Section 3.4, we did not explore this possibility. Similarly, parentheses around the property below indicate that the PKPD model can be used without the use of monitoring data.

|  | Regression model | Deep RL model | PKPD model |
| --- | --- | --- | --- |
| Requires covariates | ✓ | ✓ | (✓) |
| Requires monitoring | ✗ | ✓ | (✓) |
| Requires indefinite monitoring | ✗ | ✓ | ✗ |
| Accounts for explained IIV | ✓ | ✓ | ✓ |
| Accounts for unexplained IIV | ✗ | ✓ | ✓ |
| Accounts for IOV | ✗ | ✓ | ✓ |
| Accounts for EV | ✗ | ✓ | ✓ |
| Accounts for treatment response delays | ✗ | ✗ | ✓ |
| Robust to unseen treatment responses | ✓ | ✗ | ✓ |
| Robust to model misspecification | ✓ | ✗ | ✗ |
| Learns individual-specific dosing regimens | ✗ | ✗ | ✓ |

#### 3.5.2 The Deep RL model

In comparison, the Deep RL model is able to predict more individualised dosing regimens than the Regression model (see middle panel in Fig 8). The panel shows that for all three individuals, the model begins the treatment with increased warfarin dosages of more than 20 mg. During the maintenance phase, it reduces the dosages. For the individual with covariates *χ*_*A*_ (ID 716), the model alternates between administering dosages of 12 mg and 3 mg. For the other two individuals with covariates *χ*_*B*_, the model administers a constant dose of 3 mg, from which it only occasionally deviates for one of the individuals (ID 269). Overall, the figure shows that the Deep RL model is more successful in individualising the dosing regimens of the individuals, as indicated by their maintenance INR measurements at the end of the trial. The individualisation of the induction phase reduces the time needed to reach the therapeutic range, leading to all three individuals displaying INR measurements inside the therapeutic range within the first four treatment days.

The main reason for the improved performance of the Deep RL model is the use of feedback control from the monitoring data in addition to the covariate information. The model derives predictions from both monitoring data and covariates using the dose function, defined in Eq 11, which predicts the next-to-administer dose based on the most recently measured INR value and the covariates of the to-be-treated individual. We visualise the predictions of the dose function for the three individuals in Fig 9. For illustrative purposes, the figure focuses on INR values between 0.5 and 7. The predicted dose values are illustrated using a black line for individuals with the covariates *χ*_*A*_ and a red-blue line for individuals with the covariates *χ*_*B*_. The target treatment response is illustrated using a black dashed line, and the therapeutic range is indicated using blue dashed lines.

**Figure 9.**
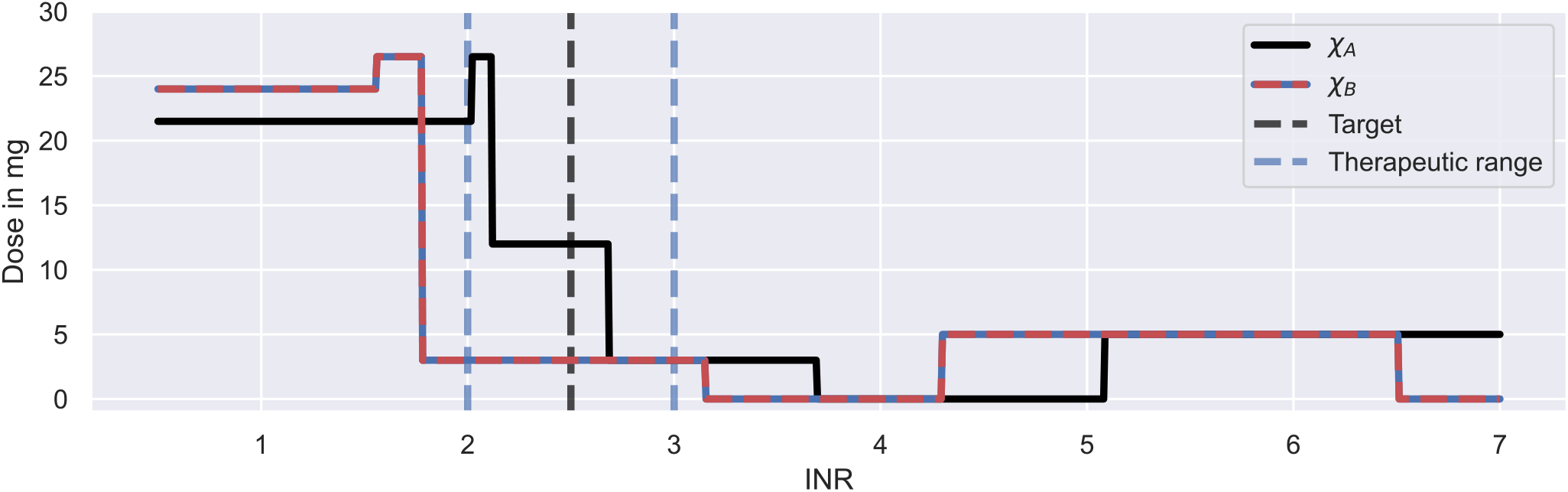
Dosing strategy of the Deep RL model. The figure illustrates the dose function of the Deep RL model, defined in Eq 11, for a range of possible INR monitoring measurements and two sets of covariates: *χ*_*A*_ = (*1*1, GG, 50) (black line); and *χ*_*B*_ = (*1*2, GA, 50) (red-blue line). The function determines the next-to-administer dose based on the most recently measured INR value (bottom axis) and the covariates of an individual. The target treatment response is illustrated using a black dashed line. The therapeutic range is indicated using blue dashed lines.

The figure shows that the Deep RL model has a clear strategy for INR monitoring measurements below 4, and a less clear strategy for INR measurements above 4. We will first focus on the dose decisions for INRs below 4: for INR measurements below the therapeutic range, the model administers large warfarin doses; for INRs inside the therapeutic range, the model administers intermediate warfarin doses; and for INRs above the therapeutic range, the model administers low warfarin doses. The exact change points and dose amounts are specific to the covariates. For example, for individuals with the *χ*_*A*_ covariates, the model starts the treatment with a dose of 21.5 mg and switches to an intermediate dose of 12 mg for INR measurements between 2.1 and 2.7 (see black line in Fig 9). This dosing strategy is consistent with the dose decisions observed in Fig 8.

The dose function illustrates both a strength and a limitation of the Deep RL model (see Fig 9). A strength of the model is that it bases its dose decisions on a simple and interpretable feedback control mechanism: when INR values are too low, it increases the warfarin dose; and when INR values are too high, it decreases the warfarin dose. These dose decisions are tailored to an individual based on the individual’s covariates. This strategy leads to a substantially higher success rate and efficiency of the dosing regimen individualisation relative to the Regression model (see Fig 7), as the feedback control enables the model to maintain treatment responses within a desired range, even when unexplained IIV, IOV and EV are present. This strength is generic to reinforcement learning approaches.

However, while the ability to utilise monitoring data for control is a strength, the model’s inability to learn from the measurements is a limitation of the Deep RL model, making the approach indefinitely reliant on monitoring data. The reliance on monitoring data is specific to the Deep RL model and is a consequence of its learning approach: the model establishes the dose function (Eq 11) prior to the treatment of individuals (see Section 3.2), never individualising the dose function based on the individual-specific monitoring data. Instead, the model treats individuals with dosing strategies exclusively based on covariates, where the monitoring data is only used as a control mechanism to steer treatment responses back to the desired treatment response, if needed (see e.g. the dosing strategy in Fig 8 for ID 269). In principle, reinforcement learning approaches can update the dose function using individual-specific monitoring data (Maier et al., 2021). However, for the Deep RL model, meaningful updates of the dose function are challenging, as the large number of model parameters of its neural network complicates the balance between fine-tuning and overfitting to the limited number of measurements.

The absence of fully individualised dosing strategies explains the observed tendency of the Deep RL model to overshoot the therapeutic range (see middle panel in Fig 7): to successfully target the therapeutic range across individuals with the same set of covariates, the initial warfarin dose predicted by the dose function needs to be high enough, so that even the weakest responders in a subpopulation reach the therapeutic range. This dose leads to INR responses above the therapeutic range for strong responders in the same subpopulation due to the substantial level of warfarin IIV not explained by covariates. The over-treatment of the strong responders is compensated for by reducing the dose for INRs above the therapeutic range so much that even the strongest responders are guaranteed to regress back to the therapeutic range (see large dose steps in Fig 9). This explains the substantial fraction of individuals with peak INRs above the therapeutic range in Fig 7, and points to a general lack of precision of the Deep RL model. This ‘control over precision’ strategy is a generic limitation of reinforcement learning approaches that do not individualise their dose functions based on the individual-specific monitoring data.

The dose function in Fig 9 also demonstrates that the Deep RL model can fail to learn meaningful dosing strategies for the full range of possible INR measurements. When INR measurements become large (≥ 5.05 for *χ*_*A*_; and ≥ 4.3 for *χ*_*B*_), the model suggests increasing the warfarin dose to 5 mg, despite the fact that higher warfarin dosages inevitably lead to even higher INR values. The root for these dose decisions lies in the training of the Deep RL model (see Section 3.2): large INR monitoring measurements remain under-explored during the training, leading to poorly tested dose decisions for large INR measurements. Those dose decisions can remain without consequence, when the model is able to control treatment responses well enough to never reach large INR values (see middle panel in Fig 8). But when an individual responds more strongly to warfarin than expected, for example due to unexplained IIV, IOV or EV, poorly tested dose decisions for large INR values can cause a failure of the model’s feedback control mechanism. This failure explains the severe over-treatments of a few individuals observed in the simulated MIPD trial in Fig 7.

The lack of exploration during the training, despite the use of a standard training procedure (see *ϵ*-greedy policy in Appendix S3), is the consequence of a mismatch between the technical assumptions of the Deep RL model and the reality of warfarin treatment responses: the Deep RL model assumes that there is no delay between dose administrations and the feedback from INR measurements, when, in fact, warfarin treatment responses have delays of up to 10 days (see e.g. Fig 3). This reduces the ability of the standard training procedure to contribute to the exploration of large INR measurements. The mismatch between the model assumptions and the treatment response delay is generic to reinforcement learning models and difficult to overcome, as the convergence and optimality of reinforcement learning centrally relies on the assumption that transition dynamics can be modelled by a Markov decision process with i.i.d. actions and i.i.d. states (Sutton and Barto, 2018). For the Deep RL model, this implies that the model has to assume that INR measurements on one day depend only on the INR measurement and the administered dose on the previous day, i.e. any dose administrations and INR measurements on earlier days cannot be taken into account. Despite those technical constraints, Zadeh et al. (2023) show that their deep reinforcement learning model is able to improve its performance, when the i.i.d. assumption of the states is explicitly violated and dose decisions are conditioned on more than one recent INR measurement. As a result, the technical assumptions of reinforcement learning, needed for convergence and optimality, may limit the ability of reinforcement learning models to account for treatment response delays, in theory, but in practice, it may be possible to overcome those limitations (Gaon and Brafman, 2020).

A summary of the strengths and limitations specific to the Deep RL model are presented in the middle column of Table 1.

#### 3.5.3 The PKPD model

The right panel in Fig 8 shows that the PKPD model achieves the highest degree of dosing regimen individualisation among the tested models. The model begins the treatment for all three individuals with a high dose of 30 mg, followed by a gradual reduction of the dose over the following days. For the individual with ID 269, this dose reduction is more rapid than for the other two individuals. Towards the end of the trial, the dosages converge to constant, individual-specific maintenance dosages. The maintenance INRs are located at the upper threshold of the therapeutic range in close proximity to each other, indicating success of the dosing regimen individualisation. All three individuals display INR measurements inside the therapeutic range within the first three treatment days.

The main reason for the good performance of the PKPD model is its use of both covariate information and monitoring data similar to the Deep RL model. The model uses the covariates to predict initial dosing regimens that target the desired treatment response for average individuals. These initial dosing regimens are further individualised based on the individual-specific monitoring data, achieving an increasing degree of individualisation as more monitoring data becomes available. This increasing degree of individualisation provides a distinct advantage over the other two MIPD models. The PKPD model derives predictions from both monitoring data and covariates using the dosing regimen function, defined in Eq 12, which predicts individualised dosing regimens based on all available INR measurements of the to-be-treated individual, the covariates, and the already-administered dosages. We visualise the values of the dosing regimen function for the three individuals in Fig 10. For illustrative purposes, the figure focuses on four days of the simulated trial: day 1; day 2; day 7; and day 16. The predicted dose values are illustrated using scatter points in two opacity levels: opaque scatter points indicate already administered dosages; and faded scatter points indicate future dose administrations. The treatment day is illustrated using dashed lines.

**Figure 10.**
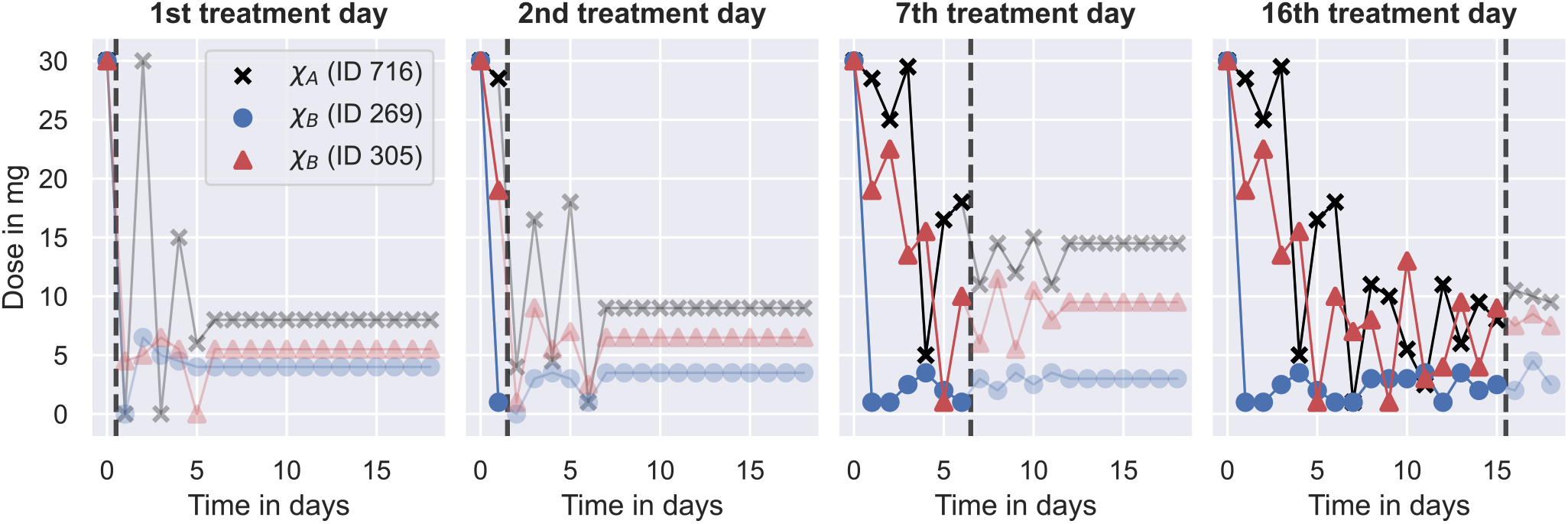
Dosing strategy of the PKPD model. The figure illustrates the dosing regimen function of the PKPD model, defined in Eq 12, for the three individuals from Fig 8 on four days of the trial: on the 1st treatment day (panel 1), on the 2nd treatment day (panel 2); on the 7th treatment day (panel 3) and on the 16th treatment day (panel 4). Dosages are illustrated using scatter points in two opacity levels: opaque scatter points for already administered dosages; faded scatter points for future dose administrations. Dosages belonging to the same individual are connected using solid lines. The day of the treatment is illustrated using black dashed lines.

The figure shows that the dosing regimen predictions are iteratively updated as the treatment progresses. On day 1, the model provides rough estimates of the dosing regimens, scheduling maintenance dosages of 8 mg (ID 716), 4 mg (ID 269), and 5.5 mg (ID 305) after an initial induction phase of the treatment. With the next INR measurement on day 2, both the induction dosing regimen as well as the maintenance dosages are updated (see second panel in Fig 10). On day 16, the model predicts fully individualised dosing regimens which are almost identical to the dosing regimens administered in the simulated trial (see right panel in Fig 8).

The dosing regimen function illustrates both a strength and a limitation of the PKPD model (see Fig 10). A strength of the model is that it predicts full dosing regimens from any number of monitoring measurements, making the dosing schedule transparent, foreseeable, and less reliant on frequent or regular monitoring. The prediction of full dosing regimens is enabled by the model’s explicit description of the pharmacological processes in terms of a semi-mechanistic model. This permits the prediction of treatment responses, and thus optimal dosing regimens, for any future time points. It also helps the model to account for treatment response delays and nonlinearities of the dose-response relationship (see Section 2.3). However, the explicit model of the treatment response bears a risk for model misspecification (Merlé et al., 2004). In particular, neglecting or oversimplifying important treatment response mechanisms can lead to inaccurate treatment response predictions, which, in turn, can impact the quality of the dosing regimen individualisation. For example, the results from the simulated MIPD trial show that the PKPD model tends to administer too large warfarin dosages during the trial, resulting in a systematic bias towards INR measurements larger than the target INR (see bottom panel of Fig 7). This indicates that the PKPD model oversimplifies crucial elements of the treatment response mechanisms, resulting in a tendency to underestimate the treatment response of individuals. The risk for model misspecification is a generic limitation of PKPD modelling, which needs to be mitigated prior to clinical applications, for example by quantifying the structural uncertainty of PKPD models using model selection criteria or probabilistic model averaging (Uster et al., 2021; Augustin et al., 2022).

The dosing regimen function in Fig 10 also illustrates that the updates of the dosing regimens themselves can result in a bias of the treatment strategy. The comparison between the predicted dosing regimens shows that although more individual-specific monitoring measurements are available on day 7 of the treatment, the maintenance dosages predicted on day 2 are closer to the actual maintenance dosages administered towards the end of the trial. This suggests that the degree of the dosing regimen individualisation can temporarily decrease with the number of monitoring measurements – a limitation specific to our implementation of the dosing regimen individualisation. The potential for worse dosing regimens despite more monitoring data is related to the estimation of the individual-specific model parameters from the monitoring data: we estimate the model parameters using Bayesian inference (see Section 2.3). The result of this estimation is a distribution of parameter values consistent with the data, also known as posterior distribution. In our implementation, we estimate the individual-specific model parameters by the modes of the distribution, also known as maximum a posteriori (MAP) estimates. The MAP estimates are a popular choice to reduce posterior distributions to just one set of model parameters (Sheiner et al., 1979). However, by disregarding the other model parameters that are also consistent with the data, it is possible to introduce biases in the treatment response predictions with consequences for the dosing regimen individualisation. In particular for nonlinear treatment responses, the treatment response predicted with the MAP estimates is, generally, not the treatment response that maximises the predictive probability (Maier et al., 2020). This increases the risk for inaccurate treatment response predictions.

A bias of the MAP-based treatment response predictions explains the lack of safety observed during the simulated MIPD trial (see bottom panel in Fig 7). The MAP-based predictions tend to underestimate the warfarin treatment response which leads to individualised dosing regimens with elevated dose amounts. As the treatment progresses, the uncertainty about the model parameters becomes smaller, reducing the error of the MAP estimation, and with it, the bias of the treatment response predictions (see maintenance INR distribution in Fig 7). The bias of the MAP-based predictions is also supported by the goodness-of-fit plot from the model fit to the pre-MIPD trial data in Section 3.2, where already the predictions with the maximum probability parameters of the population distribution show an underestimation of the treatment response for INRs inside the therapeutic range (see Fig S4.9). This limitation of our PKPD model can be mitigated by predicting the treatment response with each parameter set that is consistent with the data, i.e. the parameters in the posterior distribution. This produces a distribution of treatment responses which reflects the uncertainty in the treatment response predictions (Maier et al., 2020). The distribution of treatment responses can be optimised to obtain a dosing regimen with less risk for bias (Maier et al., 2021).

A summary of the strengths and limitations specific to the PKPD model are presented in the right column of Table 1.

## 4 CONCLUSION

Simulated clinical trials provide a resource-efficient way to test and develop fit-for-purpose models for precision dosing. We show that we can emulate clinical trials using a clinical trial model with five independent model components: 1. a mechanistic model; 2. a population model; 3. an inter-occasion model; 4. an execution model; and 5. a measurement model. Each model component captures a different complexity of clinical practice that challenges the successful individualisation of treatments, ranging from PKPD-related challenges, such as nonlinear and delayed treatment responses, to practical challenges, such as unintentional deviations from nominal dosing schedules (see Section 2.1). The modularity of the model components simplifies the development process of the clinical trial model, allowing for independent updates of its components throughout the drug development pipeline. This makes it possible to iteratively improve the trial simulations as more understanding and information about the drug under trial becomes available.

Simulating MIPD trials for warfarin, we find that different MIPD models have different strengths and limitations. These strengths and limitations can be generic to the methodology or specific to the model implementation. Modelling approaches exclusively based on covariates of the treatment response variability are generically limited when unexplained IIV, IOV and EV are present (see Section 3.5.1). However, when the majority of the treatment response variability can be explained by covariates, and those covariates are available in clinical practice, MIPD models exclusively based on covariates provide an excellent solution to the individualisation of dosing regimens. Otherwise, MIPD models based on monitoring data are better at accounting for IIV, IOV and EV, and achieve a higher degree of dosing regimen individualisation in the simulated warfarin trials (see Sections 3.4 and 3.5).

But there are also challenges with monitoring-based MIPD approaches. We find that the Deep RL model adopts a ‘control over precision’ treatment strategy, where doses are adjusted based on the feedback response from the monitoring data with limited foresight (see Section 3.5.2). This lack of precision makes the model reliant on indefinite monitoring and limits its ability to account for treatment response delays. Deep reinforcement learning may, nevertheless, provide a good solution to the individualisation of dosing regimens for applications where monitoring is not a challenge, such as for treatments in the intensive care unit (Moore et al., 2004) or for dosing devices that are physically attached to patients, like insulin pumps (Zhu et al., 2020).

Overall we find that the PKPD model achieves the highest degree of dosing regimen individualisation among the tested models. The model predicts fully individualised dosing regimens whose level of individualisation increases with the amount of available monitoring data. This indicates that across applications PKPD models are the most promising approach for model-informed precision dosing. However, PKPD models are more susceptible to model misspecifications than the other two approaches (see Section 3.5.3), necessitating a careful evaluation of the predictive uncertainty of the model prior to MIPD, for example by means of model selection criteria or probabilistic model averaging (Augustin et al., 2022).

## Footnotes

1 In this case, the close-to-average response can be explained by the individual’s model parameters, *ψ*, being close to the means of the population distribution (Eq 2).

2 Similarly to the above footnote, the stronger-than-average response can be explained by key model parameters being further away from the means of the population distribution.

