## Supplementary Text for "Simulating clinical trials for model-informed precision dosing: Using warfarin treatment as a use case"

### SUPPLEMENTAL DATA

#### Appendix S1

### S1 WARFARIN CLINICAL TRIAL MODEL

#### S1.1 The mechanistic model

We model the PKPD of warfarin using Wajima et al's QSP model defined in Table S1.1. The table shows the 53 differential equations and 52 algebraic equations of the model, describing the dynamics of warfarin, vitamin K and different coagulation factors. The parameters and variables are shown in verbose names for easier comprehesion of the large number of equations. For example, $\mathrm{d}a_d/\mathrm{d}t = -k_a a_d + r$ from Eq 6 is displayed as

$$\frac{d}{dt}\text{drug\_amount} = -\text{absorption\_rate} * \text{drug\_amount} + \text{dose\_rate}.$$

For a detailed discussion of the model we refer to (Wajima et al., 2009). To simplify the implementation of the model, we provide an SBML specification on GitHub (https://github.com/DavAug/mipd-warfarin), which can be used to simulate the model, for example using `chi` (Augustin, 2021).

##### S1.1.1 Implementation of the dosing regimen

We implement the dosing regimen using a time-dependent dose rate defined by a sequence of $K$ dose administrations

$$r(t) = \sum_{k=1}^{K} \frac{d_k}{\Delta_k}\, \Theta(t - t_k)\, \Theta(t_k + \Delta_k - t). \tag{S1.1}$$

$d_k$ denotes the dose administered at time $t_k$. The duration of the administration is denoted by $\Delta_k$. $\Theta(x)$ denotes the Heaviside step function, which is equal to 0 for $x \leq 0$ and is equal to 1 for $x > 0$. As a result, the dose rate, $r(t)$, assumes the constant value $d_k/\Delta_k$ during the $k$th dose administration. For periods outside dose administrations, the dose rate is zero.

For all simulations in this article, the oral dose of warfarin is approximated by a dose duration of $\Delta_k = 0.01\,\mathrm{h}$.

##### S1.1.2 Simulation of the prothrombin time test

The prothrombin time (PT) test quantifies the clotting time of blood. We simulate the PT test following Wajima et al. (2009): First, a blood sample is taken from the virtual patient, characterised by the states of the model (see Table S1.2). The sample is then prepared for the PT test by diluting it $1 : 2$ with a thromboplastin reagent. We implement this dilution by setting the thrombomodulin amount to 0 and by dividing all state values by a factor of 3. Lastly, we set all external input rates and endogenous production rates to 0 to reflect the *in vitro* nature of the test (see parameters marked by (†) in Table S1.3).

The test is initiated by setting the tissue factor concentration to $300$ nmol/L. The nonzero tissue factor concentration triggers the extrinsic activation pathway and leads to an increase in the fibrin concentration over time, which in real blood results in clot formation. In the model, clot formation is assumed to occur when the area under the fibrin concentration curve (fibrin AUC) reaches a threshold of $1500$ nmols/L. The time elapsed between tissue factor exposure and clot formation defines the PT time. The INR of a blood sample is given by the PT normalised by a reference PT. In our experiments, we use a reference PT of 11.8 s, which we adopted from Wajima et al. (2009).

Wajima et al's original model does not directly calculate the fibrin AUC (see Table S1.1), requiring a post-simulation integration of the fibrin concentration time curve to determine the fibrin AUC. To avoid this complication of the PT test simulation, we complement the model with an auxiliary ODE to integrate the fibrin concentration curve over time during the model simulation

$$\frac{d}{dt}\text{fibrin\_area\_under\_the\_curve} = \text{fibrin\_concentration}.$$

This makes it possible to efficiently simulate the fibrin AUC together with the rest of the model using a differential equation solver. The initial value of the fibrin AUC is set to 0.

#### S1.2 The population model

We model the treatment response variability by combining the works of Hartmann et al. (2016, 2020) and Hamberg et al. (2010). The first component of the population model is adopted from (Hartmann et al., 2016), who capture IIV using variability in the production rates of prothrombin, protein S, protein C and coagulation factors V, VII, IX, X, XI, XII and XIII (see parameters marked by (†) in Table S1.3). The rates are assumed to be normally distributed across individuals

$$p(\psi_\dagger|\theta_{\psi_\dagger}) = \mathcal{N}(\psi_\dagger|\mu_{\psi_\dagger}, \sigma^2_{\psi_\dagger}), \tag{S1.2}$$

where we use $\psi_\dagger$ to refer to the production rates. $\mu_{\psi_\dagger}$ and $\sigma_{\psi_\dagger}$ denote the mean and the standard deviation of $\psi_\dagger$ in the population. The standard deviation of the parameters is assumed to be equal to $20\,\%$ of their mean value, $\sigma_{\psi_\dagger} = 0.2\mu_{\psi_\dagger}$.

The second component of the population model is a modification of Hartmann et al's covariate model (Hartmann et al., 2020) based on the work of (Hamberg et al., 2013). Hartmann et al's model introduces VKORC1 as a covariate of warfarin's EC50 variability and CYP2C9 as a covariate of warfarin's clearance. The typical EC50 is reduced by $22\,\%$ relative to the GG VKORC1 genotype for each A allele

$$\mu_{c_{50}}(v_1, v_2) \quad = \quad c_{v_1} + c_{v_2}, \tag{S1.3}$$

where $v_1, v_2 \in \{G, A\}$ denote the VKORC1 alleles and $c_A = 0.66\, c_G$. Hamberg et al. (2013) use a structurally identical model for the explained EC50 variability, but also model unexplained variability using random deviations from the typical value, $c_{50} = \mu_{c_{50}}\, \mathrm{e}^{\eta_{c_{50}}}$. $\eta_{c_{50}}$ is a Gaussian random variable with zero mean and variance $\sigma^2_{c_{50}}$. This defines a lognormal EC50 distribution in the population

$$p(c_{50}|\theta_{c_{50}}, \chi_{c_{50}}) = \mathrm{LN}\left(c_{50}\,|\,\mu_{c_{50}}(v_1, v_2), \sigma_{c_{50}}\right). \tag{S1.4}$$

In our model, we adopt this population model for the EC50.

The typical clearance in Hartmann et al's model is reduced by $30\,\%$ for the *2*2 CYP2C9 genotype and by $80\,\%$ for the *3*3 genotype relative to the *1*1 genotype (Hartmann et al., 2020). In Hamberg et al's model, the clearance contributions of the CYP2C9 alleles are modelled additively Hamberg et al. (2013). In addition, they reduce the clearance with the age of the patient. We adopt Hamberg et al's model structure and use the relationship between the clearance and the elimination rate, clearance $=$ volume $*$ elimination_rate, to define the typical elimination rate as a function of the CYP2C9 alleles and the age

$$\mu_{k_e}(c_1, c_2, t_\star) = (k_{c_1} + k_{c_2})\,(1 - \tanh(r_{age}(t_\star - 71)))\,. \tag{S1.5}$$

$c_1, c_2 \in \{*1, *2, *3\}$ denote the CYP2C9 alleles, $t_\star$ denotes the age and $r_{age}$ denotes the reduction rate of the elimination rate with age. Analogous to the EC50, we use a lognormal distribution to model the unexplained variability of the elimination rate

$$p(k_e|\theta_{k_e}, \chi_{k_e}) = \text{LN}\left(k_e \,|\, \mu_{k_e}(c_1, c_2, t_\star), \sigma_{k_e}\right). \tag{S1.6}$$

We also adopt Hamberg et al's model of the variability of the volume of distribution, $v$, which describes the IIV using a lognormal distribution with no additional subpopulation structure

$$p(v|\theta_v) = \text{LN}\left(v \,|\, \mu_v, \sigma_v\right). \tag{S1.7}$$

The third component further extends the covariate model and implements the reduced ability of VKORC to promote the conversion between vitamin K and vitamin K hydroquinone (VKH2), as well as vitamin K epoxide (VKO) and vitamin K when polymorphisms in VKORC1 are present (Rost et al., 2004). We implement this effect by applying the EC50 covariate model to the conversion rates

$$\begin{aligned} \mu_{r_{\text{vk}\to\text{vkh2}}}(v_1, v_2) &= r_{\text{vk}\to\text{vkh2},v_1} + r_{\text{vk}\to\text{vkh2},v_2} \\ \mu_{r_{\text{vko}\to\text{vk}}}(v_1, v_2) &= r_{\text{vko}\to\text{vk},v_1} + r_{\text{vko}\to\text{vk},v_2}. \end{aligned} \tag{S1.8}$$

The unexplained variability of the conversion rates in the population is captured using lognormal distributions

$$\begin{aligned} p(r_{\text{vk}\to\text{vkh2}}|\theta_{r_{\text{vk}\to\text{vkh2}}}, \chi_{r_{\text{vk}\to\text{vkh2}}}) &= \text{LN}\left(r_{\text{vk}\to\text{vkh2}} \,|\, \mu_{r_{\text{vk}\to\text{vkh2}}}(v_1, v_2), \sigma_{r_{\text{vk}\to\text{vkh2}}}\right) \\ p(r_{\text{vko}\to\text{vk}}|\theta_{r_{\text{vko}\to\text{vk}}}, \chi_{r_{\text{vko}\to\text{vk}}}) &= \text{LN}\left(r_{\text{vko}\to\text{vk}} \,|\, \mu_{r_{\text{vko}\to\text{vk}}}(v_1, v_2), \sigma_{r_{\text{vko}\to\text{vk}}}\right). \end{aligned} \tag{S1.9}$$

The remaining parameters of the model are assumed to display no IIV.

In summary, the population model of the warfarin clinical trial model describes IIV using variability in the EC50, the elimination rate, the vitamin K conversion rates, the volume of distribution as well the production rates of different coagulation factors

$$\begin{aligned} p(\psi|\theta, \chi) = \;& p(c_{50}|\theta_{c_{50}}, \chi_{c_{50}})\, p(k_e|\theta_{k_e}, \chi_{k_e})\, p(r_{\text{vk}\to\text{vkh2}}|\theta_{r_{\text{vk}\to\text{vkh2}}}, \chi_{r_{\text{vk}\to\text{vkh2}}}) \\ & p(r_{\text{vko}\to\text{vk}}|\theta_{r_{\text{vko}\to\text{vk}}}, \chi_{r_{\text{vko}\to\text{vk}}})\, p(v|\theta_v) \prod_{\dagger} p(\psi_\dagger|\theta_{\psi_\dagger}) \prod_k \delta(\mu_{\psi_k} - \psi_k). \end{aligned} \tag{S1.10}$$

All remaining parameters, here indexed by $k$, display no IIV. The parameters of the model, $\theta$, are the parameters of the individual marginal distributions. The covariates of the model are the VKORC1 alleles, the CYP2C9 alleles and the age, $\chi = (v_1, v_2, c_1, c_2, t_\star)$.

The full set of parameter values used for the simulation is provided in a CSV file on GitHub (https://github.com/DavAug/mipd-warfarin). The parameter values are largely adopted from Hartmann et al. (2016, 2020) and Hamberg et al. (2010). However, IIV in the conversion rate variability was not previously captured by the models, leaving the model parameters in Eq S1.9 unspecified.

To calibrate the parameters of the conversion rate distributions, we first set the typical conversion rates for individuals with the GG VKORC1 genotype to the conversion rates provided by Hartmann et al. (2020). The reduction of the typical rates with VKORC1 polymorphisms and the scale parameter of the distribution are adopted from the EC50 distribution. Simulating the baseline INR without warfarin treatment for 1000 random individuals shows that this initial set of parameter values produces INRs around 1, indicating reasonable simulation results (see black line in Fig S1.1). However, for some individuals, the baseline INR displays values $> 2$, which we identify as unrealistically high. To avoid too high baseline INRs, we reduce the effect of VKORC1 polymorphisms to a total reduction of the typical conversion rates of $20\,\%$ relative to the GG genotype. This results in a baseline INR distribution with less extreme values (see blue histogram in Fig S1.1). It should be noted that we did not have access to real baseline INR measurements for this calibration. Calibrating the model to real measurements will likely improve the parameter estimates.

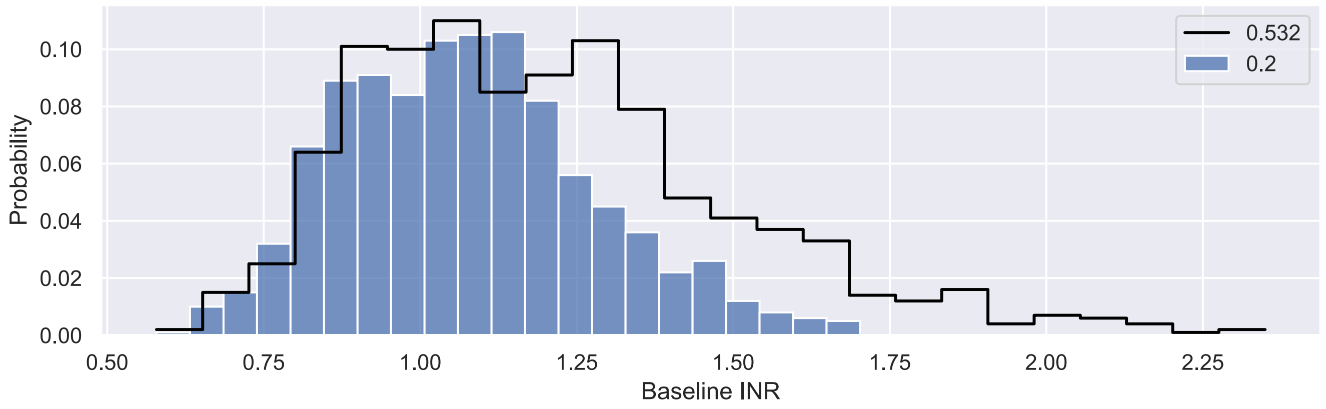


**Figure S1.1. Calibration of baseline INR distribution.** The figure shows the histogram over baseline INR values for 1000 virtual patients simulated with the warfarin clinical trial model before (black) and after (blue) the calibration. Before the calibration the model assumes a $53.2\,\%$ reduction of the vitamin K conversion rate for individuals with the AA VKORC1 genotype relative to individuals with the GG VKORC1 genotype. The calibrated model assumes reductions of $20\,\%$.

In a second calibration step, we adjust the G VKORC1 allele contribution to the EC50 and the scale parameter of the EC50 distribution. This is necessary because: 1. adopting Hamberg et al's value for the volume of distribution shifts the simulated warfarin concentrations relative to Hartmann et al's model which is not accounted for when keeping the original $c_G$ value in Eq S1.3; and 2. the detailed description of warfarin's mode of action in the clinical trial model distinguishes variability contributions from the EC50 and the vitamin K conversion rates to the overall treatment response variability. This reduces the variability contribution from the EC50 variability relative to Hamberg et al's model.

To calibrate the two parameters, we simulate the maintenance warfarin dose distribution and compare it to clinical maintenance doses reported in IWPC (2009). A challenge with this comparison is that individuals in the clinical dataset have varying INR values. For the comparison, we therefore focus on individuals with INRs inside the therapeutic range. To generate a maintenance dose distribution with the clinical trial model, we simulate the treatment response of 1000 individuals under 56 days of warfarin treatment. We start the treatment for all individuals with the same dose (10 mg). We then measure the INR every 7 days and adjust the dose using linear extrapolation ($d' = d\, y^*/y$), when the INR is outside the therapeutic range. $d$ denotes the current dose, $d'$ denotes the adjusted dose, $y$ denotes the INR measurement and $y^* = 2.5$ denotes the target INR. If the measurement is inside the therapeutic range, the dose is not adjusted. We classify an individual as ‘having reached maintenance treatment’ if the individual’s INR measurement is inside the therapeutic range for the last 3 weeks of the simulation.

In S1.2, we show the clinical maintenance dose distribution (black curve) and the simulated maintenance dose distribution (blue histogram) after the model calibration. The two distributions are in good agreement.

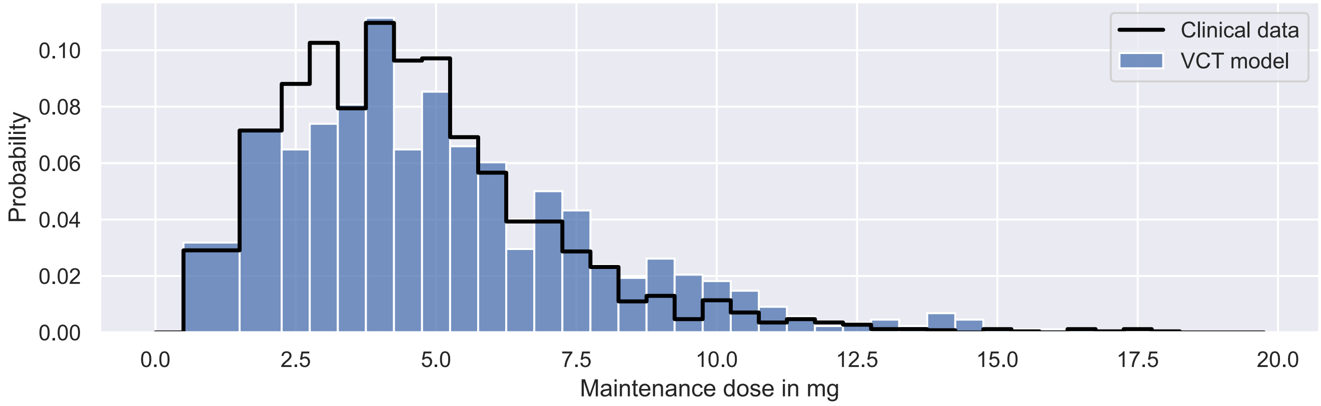


**Figure S1.2. Clinical maintenance dose distribution and simulated maintenance dose distribution.** The figure shows the histogram over warfarin maintenance doses administered in clinical practice (black) and during the clinical trial simulation (blue). The clinical data contains maintenance dosages from 2543 individuals with INRs inside the TR. The data simulated with the clinical trial mdoel contains maintenance dosages from 846 virtual patients with INRs inside the TR.

**Table S1.1.** Definition of the mechanistic model

$$
\begin{aligned}
&\frac{d}{dt}\text{coagulation\_factor\_ii\_amount} \\
&= \text{central\_size} * \text{production\_rate\_ii} * \text{vitamin\_k\_hydroquinone\_concentration} - \text{central\_size} * \text{degradation\_rate\_ii} * \text{coagulation\_factor\_ii\_concentration} \\
&\quad - \frac{\text{central\_size} * \text{maximal\_activation\_rate\_av\_ax\_complex\_ii} * \text{av\_ax\_complex\_concentration} * \text{coagulation\_factor\_ii\_concentration}}{\text{half\_maximal\_effect\_concentration\_av\_ax\_complex\_ii} + \text{av\_ax\_complex\_concentration}} \\
&\quad - \frac{\text{central\_size} * \text{maximal\_activation\_rate\_ax\_ii} * \text{activated\_coagulation\_factor\_x\_concentration} * \text{coagulation\_factor\_ii\_concentration}}{\text{half\_maximal\_effect\_concentration\_ax\_ii} + \text{activated\_coagulation\_factor\_x\_concentration}}
\end{aligned}
$$

$$
\begin{aligned}
&\frac{d}{dt}\text{activated\_coagulation\_factor\_ii\_amount} \\
&= \frac{\text{central\_size} * \text{maximal\_activation\_rate\_av\_ax\_complex\_ii} * \text{av\_ax\_complex\_concentration} * \text{coagulation\_factor\_ii\_concentration}}{\text{half\_maximal\_effect\_concentration\_av\_ax\_complex\_ii} + \text{av\_ax\_complex\_concentration}} \\
&\quad + \frac{\text{central\_size} * \text{maximal\_activation\_rate\_ax\_ii} * \text{activated\_coagulation\_factor\_x\_concentration} * \text{coagulation\_factor\_ii\_concentration}}{\text{half\_maximal\_effect\_concentration\_ax\_ii} + \text{activated\_coagulation\_factor\_x\_concentration}} \\
&\quad - \text{central\_size} * \text{degradation\_rate\_ii\_activated} * \text{activated\_coagulation\_factor\_ii\_concentration} - \text{central\_size} \\
&\quad * \text{complex\_formation\_rate\_aii\_atiii\_heparin\_complex} * \text{activated\_coagulation\_factor\_ii\_concentration} * \text{atiii\_heparin\_complex\_concentration} \\
&\quad - \text{central\_size} * \text{complex\_formation\_rate\_aii\_tmod} * \text{activated\_coagulation\_factor\_ii\_concentration} * \text{thrombomodulin\_concentration}
\end{aligned}
$$

$$
\begin{aligned}
&\frac{d}{dt}\text{coagulation\_factor\_v\_amount} \\
&= \text{central\_size} * \text{production\_rate\_v} - \text{central\_size} * \text{degradation\_rate\_v} * \text{coagulation\_factor\_v\_concentration} \\
&\quad - \frac{\text{central\_size} * \text{maximal\_activation\_rate\_thrombin\_v} * \text{activated\_coagulation\_factor\_ii\_concentration} * \text{coagulation\_factor\_v\_concentration}}{\text{half\_maximal\_effect\_concentration\_thrombin\_v} + \text{activated\_coagulation\_factor\_ii\_concentration}}
\end{aligned}
$$

$$
\begin{aligned}
&\frac{d}{dt}\text{activated\_coagulation\_factor\_v\_amount} \\
&= \frac{\text{central\_size} * \text{maximal\_activation\_rate\_thrombin\_v} * \text{activated\_coagulation\_factor\_ii\_concentration} * \text{coagulation\_factor\_v\_concentration}}{\text{half\_maximal\_effect\_concentration\_thrombin\_v} + \text{activated\_coagulation\_factor\_ii\_concentration}} \\
&\quad - \text{central\_size} * \text{degradation\_rate\_v\_activated} * \text{activated\_coagulation\_factor\_v\_concentration} \\
&\quad - \frac{\text{central\_size} * \text{maximal\_degradation\_rate\_apc\_ps\_complex\_av} * \text{apc\_ps\_complex\_concentration} * \text{activated\_coagulation\_factor\_v\_concentration}}{\text{half\_maximal\_effect\_concentration\_apc\_ps\_complex\_av} + \text{apc\_ps\_complex\_concentration}} \\
&\quad - \text{central\_size} * \text{complex\_formation\_rate\_av\_ax} * \text{activated\_coagulation\_factor\_v\_concentration} * \text{activated\_coagulation\_factor\_x\_concentration}
\end{aligned}
$$

$$
\begin{aligned}
&\frac{d}{dt}\text{coagulation\_factor\_vii\_amount} \\
&= \text{central\_size} * \text{production\_rate\_vii} * \text{vitamin\_k\_hydroquinone\_concentration} - \text{central\_size} * \text{degradation\_rate\_vii} * \text{coagulation\_factor\_vii\_concentration} \\
&\quad - \frac{\text{central\_size} * \text{maximal\_activation\_rate\_thrombin\_vii} * \text{activated\_coagulation\_factor\_ii\_concentration} * \text{coagulation\_factor\_vii\_concentration}}{\text{half\_maximal\_effect\_concentration\_thrombin\_vii} + \text{activated\_coagulation\_factor\_ii\_concentration}} \\
&\quad - \frac{\text{central\_size} * \text{maximal\_activation\_rate\_aix\_vii} * \text{activated\_coagulation\_factor\_ix\_concentration} * \text{coagulation\_factor\_vii\_concentration}}{\text{half\_maximal\_effect\_concentration\_aix\_vii} + \text{activated\_coagulation\_factor\_ix\_concentration}} \\
&\quad - \frac{\text{central\_size} * \text{maximal\_activation\_rate\_ax\_vii} * \text{activated\_coagulation\_factor\_x\_concentration} * \text{coagulation\_factor\_vii\_concentration}}{\text{half\_maximal\_effect\_concentration\_ax\_vii} + \text{activated\_coagulation\_factor\_x\_concentration}} \\
&\quad - \frac{\text{central\_size} * \text{maximal\_activation\_rate\_avii\_tf\_complex\_vii} * \text{avii\_tf\_complex\_concentration} * \text{coagulation\_factor\_vii\_concentration}}{\text{half\_maximal\_effect\_concentration\_avii\_tf\_complex\_vii} + \text{avii\_tf\_complex\_concentration}} \\
&\quad - \text{central\_size} * \text{complex\_formation\_rate\_vii\_tf} * \text{coagulation\_factor\_vii\_concentration} * \text{tissue\_factor\_concentration}
\end{aligned}
$$

$$
\begin{aligned}
&\frac{d}{dt}\text{activated\_coagulation\_factor\_vii\_amount} \\
&= \frac{\text{central\_size} * \text{maximal\_activation\_rate\_thrombin\_vii} * \text{activated\_coagulation\_factor\_ii\_concentration} * \text{coagulation\_factor\_vii\_concentration}}{\text{half\_maximal\_effect\_concentration\_thrombin\_vii} + \text{activated\_coagulation\_factor\_ii\_concentration}} \\
&\quad + \frac{\text{central\_size} * \text{maximal\_activation\_rate\_aix\_vii} * \text{activated\_coagulation\_factor\_ix\_concentration} * \text{coagulation\_factor\_vii\_concentration}}{\text{half\_maximal\_effect\_concentration\_aix\_vii} + \text{activated\_coagulation\_factor\_ix\_concentration}} \\
&\quad + \frac{\text{central\_size} * \text{maximal\_activation\_rate\_ax\_vii} * \text{activated\_coagulation\_factor\_x\_concentration} * \text{coagulation\_factor\_vii\_concentration}}{\text{half\_maximal\_effect\_concentration\_ax\_vii} + \text{activated\_coagulation\_factor\_x\_concentration}} \\
&\quad + \frac{\text{central\_size} * \text{maximal\_activation\_rate\_avii\_tf\_complex\_vii} * \text{avii\_tf\_complex\_concentration} * \text{coagulation\_factor\_vii\_concentration}}{\text{half\_maximal\_effect\_concentration\_avii\_tf\_complex\_vii} + \text{avii\_tf\_complex\_concentration}} \\
&\quad - \text{central\_size} * \text{degradation\_rate\_vii\_activated} * \text{activated\_coagulation\_factor\_vii\_concentration} - \text{central\_size} \\
&\quad * \text{complex\_formation\_rate\_avii\_tf} * \text{activated\_coagulation\_factor\_vii\_concentration} * \text{tissue\_factor\_concentration}
\end{aligned}
$$

$$
\begin{aligned}
&\frac{d}{dt}\text{coagulation\_factor\_viii\_amount} \\
&= \text{central\_size} * \text{production\_rate\_viii} - \text{central\_size} * \text{degradation\_rate\_viii} * \text{coagulation\_factor\_viii\_concentration} \\
&\quad - \frac{\text{central\_size} * \text{maximal\_activation\_rate\_thrombin\_viii} * \text{activated\_coagulation\_factor\_ii\_concentration} * \text{coagulation\_factor\_viii\_concentration}}{\text{half\_maximal\_effect\_concentration\_thrombin\_viii} + \text{activated\_coagulation\_factor\_ii\_concentration}}
\end{aligned}
$$

$$
\begin{aligned}
&\frac{d}{dt}\text{activated\_coagulation\_factor\_viii\_amount} \\
&= \frac{\text{central\_size} * \text{maximal\_activation\_rate\_thrombin\_viii} * \text{activated\_coagulation\_factor\_ii\_concentration} * \text{coagulation\_factor\_viii\_concentration}}{\text{half\_maximal\_effect\_concentration\_thrombin\_viii} + \text{activated\_coagulation\_factor\_ii\_concentration}} \\
&\quad - \text{central\_size} * \text{degradation\_rate\_viii\_activated} * \text{activated\_coagulation\_factor\_viii\_concentration} \\
&\quad - \frac{\text{central\_size} * \text{maximal\_degradation\_rate\_apc\_ps\_complex\_aviii} * \text{apc\_ps\_complex\_concentration} * \text{activated\_coagulation\_factor\_viii\_concentration}}{\text{half\_maximal\_effect\_concentration\_apc\_ps\_complex\_aviii} + \text{apc\_ps\_complex\_concentration}} \\
&\quad - \text{central\_size} * \text{complex\_formation\_rate\_aix\_aviii} * \text{activated\_coagulation\_factor\_ix\_concentration} * \text{activated\_coagulation\_factor\_viii\_concentration}
\end{aligned}
$$

$$
\begin{aligned}
&\frac{d}{dt}\text{coagulation\_factor\_ix\_amount} \\
&= \text{central\_size} * \text{production\_rate\_ix} * \text{vitamin\_k\_hydroquinone\_concentration} - \text{central\_size} * \text{degradation\_rate\_ix} * \text{coagulation\_factor\_ix\_concentration} \\
&\quad - \frac{\text{central\_size} * \text{maximal\_activation\_rate\_axi\_ix} * \text{activated\_coagulation\_factor\_xi\_concentration} * \text{coagulation\_factor\_ix\_concentration}}{\text{half\_maximal\_effect\_concentration\_axi\_ix} + \text{activated\_coagulation\_factor\_xi\_concentration}} \\
&\quad - \frac{\text{central\_size} * \text{maximal\_activation\_rate\_avii\_tf\_complex\_ix} * \text{avii\_tf\_complex\_concentration} * \text{coagulation\_factor\_ix\_concentration}}{\text{half\_maximal\_effect\_concentration\_avii\_tf\_complex\_ix} + \text{avii\_tf\_complex\_concentration}}
\end{aligned}
$$

$$
\begin{aligned}
&\frac{d}{dt}\text{activated\_coagulation\_factor\_ix\_amount} \\
&= \frac{\text{central\_size} * \text{maximal\_activation\_rate\_axi\_ix} * \text{activated\_coagulation\_factor\_xi\_concentration} * \text{coagulation\_factor\_ix\_concentration}}{\text{half\_maximal\_effect\_concentration\_axi\_ix} + \text{activated\_coagulation\_factor\_xi\_concentration}} \\
&\quad + \frac{\text{central\_size} * \text{maximal\_activation\_rate\_avii\_tf\_complex\_ix} * \text{avii\_tf\_complex\_concentration} * \text{coagulation\_factor\_ix\_concentration}}{\text{half\_maximal\_effect\_concentration\_avii\_tf\_complex\_ix} + \text{avii\_tf\_complex\_concentration}} \\
&\quad - \text{central\_size} * \text{degradation\_rate\_ix\_activated} * \text{activated\_coagulation\_factor\_ix\_concentration} - \text{central\_size} * \text{complex\_formation\_rate\_aix\_aviii} \\
&\quad * \text{activated\_coagulation\_factor\_ix\_concentration} * \text{activated\_coagulation\_factor\_viii\_concentration} - \text{central\_size} \\
&\quad * \text{complex\_formation\_rate\_aix\_atiii\_heparin\_complex} * \text{activated\_coagulation\_factor\_ix\_concentration} * \text{atiii\_heparin\_complex\_concentration}
\end{aligned}
$$

$$
\begin{aligned}
&\frac{d}{dt}\text{coagulation\_factor\_x\_amount}\\
&= \text{central\_size} * \text{production\_rate\_x} * \text{vitamin\_k\_hydroquinone\_concentration} - \text{central\_size} * \text{degradation\_rate\_x} * \text{coagulation\_factor\_x\_concentration}\\
&\quad - \frac{\text{central\_size} * \text{maximal\_activation\_rate\_avii\_x} * \text{activated\_coagulation\_factor\_vii\_concentration} * \text{coagulation\_factor\_x\_concentration}}{\text{half\_maximal\_effect\_concentration\_avii\_x} + \text{activated\_coagulation\_factor\_vii\_concentration}}\\
&\quad - \frac{\text{central\_size} * \text{maximal\_activation\_rate\_avii\_tf\_complex\_x} * \text{avii\_tf\_complex\_concentration} * \text{coagulation\_factor\_x\_concentration}}{\text{half\_maximal\_effect\_concentration\_avii\_tf\_complex\_x} + \text{avii\_tf\_complex\_concentration}}\\
&\quad - \frac{\text{central\_size} * \text{maximal\_activation\_rate\_aix\_x} * \text{activated\_coagulation\_factor\_ix\_concentration} * \text{coagulation\_factor\_x\_concentration}}{\text{half\_maximal\_effect\_concentration\_aix\_x} + \text{activated\_coagulation\_factor\_ix\_concentration}}\\
&\quad - \frac{\text{central\_size} * \text{maximal\_activation\_rate\_aix\_aviii\_complex\_x} * \text{aix\_aviii\_complex\_concentration} * \text{coagulation\_factor\_x\_concentration}}{\text{half\_maximal\_effect\_concentration\_aix\_aviii\_complex\_x} + \text{aix\_aviii\_complex\_concentration}}
\end{aligned}
$$

$$
\begin{aligned}
&\frac{d}{dt}\text{activated\_coagulation\_factor\_x\_amount}\\
&= -\text{central\_size} * \text{complex\_formation\_rate\_av\_ax} * \text{activated\_coagulation\_factor\_v\_concentration} * \text{activated\_coagulation\_factor\_x\_concentration}\\
&\quad + \frac{\text{central\_size} * \text{maximal\_activation\_rate\_avii\_x} * \text{activated\_coagulation\_factor\_vii\_concentration} * \text{coagulation\_factor\_x\_concentration}}{\text{half\_maximal\_effect\_concentration\_avii\_x} + \text{activated\_coagulation\_factor\_vii\_concentration}}\\
&\quad + \frac{\text{central\_size} * \text{maximal\_activation\_rate\_avii\_tf\_complex\_x} * \text{avii\_tf\_complex\_concentration} * \text{coagulation\_factor\_x\_concentration}}{\text{half\_maximal\_effect\_concentration\_avii\_tf\_complex\_x} + \text{avii\_tf\_complex\_concentration}}\\
&\quad + \frac{\text{central\_size} * \text{maximal\_activation\_rate\_aix\_x} * \text{activated\_coagulation\_factor\_ix\_concentration} * \text{coagulation\_factor\_x\_concentration}}{\text{half\_maximal\_effect\_concentration\_aix\_x} + \text{activated\_coagulation\_factor\_ix\_concentration}}\\
&\quad + \frac{\text{central\_size} * \text{maximal\_activation\_rate\_aix\_aviii\_complex\_x} * \text{aix\_aviii\_complex\_concentration} * \text{coagulation\_factor\_x\_concentration}}{\text{half\_maximal\_effect\_concentration\_aix\_aviii\_complex\_x} + \text{aix\_aviii\_complex\_concentration}}\\
&\quad - \text{central\_size} * \text{degradation\_rate\_x\_activated} * \text{activated\_coagulation\_factor\_x\_concentration} - \text{central\_size}\\
&\quad * \text{complex\_formation\_rate\_ax\_atiii\_heparin\_complex} * \text{activated\_coagulation\_factor\_x\_concentration} * \text{atiii\_heparin\_complex\_concentration}\\
&\quad - \text{central\_size} * \text{complex\_formation\_rate\_ax\_tfpi} * \text{activated\_coagulation\_factor\_x\_concentration} * \text{tissue\_factor\_pathway\_inhibitor\_concentration}
\end{aligned}
$$

$$
\begin{aligned}
&\frac{d}{dt}\text{coagulation\_factor\_xi\_amount}\\
&= \text{central\_size} * \text{production\_rate\_xi} - \text{central\_size} * \text{degradation\_rate\_xi} * \text{coagulation\_factor\_xi\_concentration}\\
&\quad - \frac{\text{central\_size} * \text{maximal\_activation\_rate\_thrombin\_xi} * \text{activated\_coagulation\_factor\_ii\_concentration} * \text{coagulation\_factor\_xi\_concentration}}{\text{half\_maximal\_effect\_concentration\_thrombin\_xi} + \text{activated\_coagulation\_factor\_ii\_concentration}}\\
&\quad - \frac{\text{central\_size} * \text{maximal\_activation\_rate\_axii\_xi} * \text{activated\_coagulation\_factor\_xii\_concentration} * \text{coagulation\_factor\_xi\_concentration}}{\text{half\_maximal\_effect\_concentration\_axii\_xi} + \text{activated\_coagulation\_factor\_xii\_concentration}}
\end{aligned}
$$

$$\frac{d}{dt}\text{activated\_coagulation\_factor\_xi\_amount} = \frac{\text{central\_size} * \text{maximal\_activation\_rate\_thrombin\_xi} * \text{activated\_coagulation\_factor\_ii\_concentration} * \text{coagulation\_factor\_xi\_concentration}}{\text{half\_maximal\_effect\_concentration\_thrombin\_xi} + \text{activated\_coagulation\_factor\_ii\_concentration}} + \frac{\text{central\_size} * \text{maximal\_activation\_rate\_axii\_xi} * \text{activated\_coagulation\_factor\_xii\_concentration} * \text{coagulation\_factor\_xi\_concentration}}{\text{half\_maximal\_effect\_concentration\_axii\_xi} + \text{activated\_coagulation\_factor\_xii\_concentration}} - \text{central\_size} * \text{degradation\_rate\_xi\_activated} * \text{activated\_coagulation\_factor\_xi\_concentration}$$

$$\frac{d}{dt}\text{coagulation\_factor\_xii\_amount} = \text{central\_size} * \text{production\_rate\_xii} - \text{central\_size} * \text{degradation\_rate\_xii} * \text{coagulation\_factor\_xii\_concentration} - \frac{\text{central\_size} * \text{maximal\_activation\_rate\_ca\_xii} * \text{contact\_system\_activator\_concentration} * \text{coagulation\_factor\_xii\_concentration}}{\text{half\_maximal\_effect\_concentration\_ca\_xii} + \text{contact\_system\_activator\_concentration}} - \frac{\text{central\_size} * \text{maximal\_activation\_rate\_kallikrein\_xii} * \text{kallikrein\_concentration} * \text{coagulation\_factor\_xii\_concentration}}{\text{half\_maximal\_effect\_concentration\_kallikrein\_xii} + \text{kallikrein\_concentration}}$$

$$\frac{d}{dt}\text{activated\_coagulation\_factor\_xii\_amount} = \frac{\text{central\_size} * \text{maximal\_activation\_rate\_ca\_xii} * \text{contact\_system\_activator\_concentration} * \text{coagulation\_factor\_xii\_concentration}}{\text{half\_maximal\_effect\_concentration\_ca\_xii} + \text{contact\_system\_activator\_concentration}} + \frac{\text{central\_size} * \text{maximal\_activation\_rate\_kallikrein\_xii} * \text{kallikrein\_concentration} * \text{coagulation\_factor\_xii\_concentration}}{\text{half\_maximal\_effect\_concentration\_kallikrein\_xii} + \text{kallikrein\_concentration}} - \text{central\_size} * \text{degradation\_rate\_xii\_activated} * \text{activated\_coagulation\_factor\_xii\_concentration}$$

$$\frac{d}{dt}\text{coagulation\_factor\_xiii\_amount} = \text{central\_size} * \text{production\_rate\_xiii} - \text{central\_size} * \text{degradation\_rate\_xiii} * \text{coagulation\_factor\_xiii\_concentration} - \frac{\text{central\_size} * \text{maximal\_activation\_rate\_thrombin\_xiii} * \text{activated\_coagulation\_factor\_ii\_concentration} * \text{coagulation\_factor\_xiii\_concentration}}{\text{half\_maximal\_effect\_concentration\_thrombin\_xiii} + \text{activated\_coagulation\_factor\_ii\_concentration}}$$

$$\frac{d}{dt}\text{activated\_coagulation\_factor\_xiii\_amount} = \frac{\text{central\_size} * \text{maximal\_activation\_rate\_thrombin\_xiii} * \text{activated\_coagulation\_factor\_ii\_concentration} * \text{coagulation\_factor\_xiii\_concentration}}{\text{half\_maximal\_effect\_concentration\_thrombin\_xiii} + \text{activated\_coagulation\_factor\_ii\_concentration}} - \text{central\_size} * \text{degradation\_rate\_xiii\_activated} * \text{activated\_coagulation\_factor\_xiii\_concentration}$$

$$\frac{d}{dt}\text{contact\_system\_activator\_amount} = -\text{central\_size} * \text{degradation\_rate\_ca} * \text{contact\_system\_activator\_concentration}$$

$$\begin{aligned}\frac{d}{dt}\text{fibrin\_amount} =& \frac{\text{central\_size} * \text{maximal\_conversion\_rate\_thrombin\_fibrinogen} * \text{activated\_coagulation\_factor\_ii\_concentration} * \text{fibrinogen\_concentration}}{\text{half\_maximal\_effect\_concentration\_thrombin\_fibrinogen} + \text{activated\_coagulation\_factor\_ii\_concentration}}\\ &- \text{central\_size} * \text{degradation\_rate\_fibrin} * \text{fibrin\_concentration}\\ &- \frac{\text{central\_size} * \text{maximal\_conversion\_rate\_axiii\_fibrin} * \text{activated\_coagulation\_factor\_xiii\_concentration} * \text{fibrin\_concentration}}{\text{half\_maximal\_effect\_concentration\_axiii\_fibrin} + \text{activated\_coagulation\_factor\_xiii\_concentration}}\\ &- \frac{\text{central\_size} * \text{maximal\_degradation\_rate\_plasmin\_fibrin} * \text{plasmin\_concentration} * \text{fibrin\_concentration}}{\text{half\_maximal\_effect\_concentration\_plasmin\_fibrin} + \text{plasmin\_concentration}}\end{aligned}$$

$$\begin{aligned}&\frac{d}{dt}\text{cross\_linked\_fibrin\_amount}\\ &= \frac{\text{central\_size} * \text{maximal\_conversion\_rate\_axiii\_fibrin} * \text{activated\_coagulation\_factor\_xiii\_concentration} * \text{fibrin\_concentration}}{\text{half\_maximal\_effect\_concentration\_axiii\_fibrin} + \text{activated\_coagulation\_factor\_xiii\_concentration}}\\ &\quad - \text{central\_size} * \text{degradation\_rate\_xf} * \text{cross\_linked\_fibrin\_concentration}\\ &\quad - \frac{\text{central\_size} * \text{maximal\_degradation\_rate\_plasmin\_xf} * \text{plasmin\_concentration} * \text{cross\_linked\_fibrin\_concentration}}{\text{half\_maximal\_effect\_concentration\_plasmin\_xf} + \text{plasmin\_concentration}}\\ &\quad - \frac{\text{central\_size} * \text{maximal\_degradation\_rate\_apc\_ps\_complex\_xf} * \text{apc\_ps\_complex\_concentration} * \text{cross\_linked\_fibrin\_concentration}}{\text{half\_maximal\_effect\_concentration\_apc\_ps\_complex\_xf} + \text{apc\_ps\_complex\_concentration}}\end{aligned}$$

$$\begin{aligned}&\frac{d}{dt}\text{fibrin\_degradation\_product\_amount}\\ &= \text{central\_size} * \text{degradation\_rate\_fibrinogen} * \text{fibrinogen\_concentration}\\ &\quad + \frac{\text{central\_size} * \text{maximal\_degradation\_rate\_plasmin\_fibrinogen} * \text{plasmin\_concentration} * \text{fibrinogen\_concentration}}{\text{half\_maximal\_effect\_concentration\_plasmin\_fibrinogen} + \text{plasmin\_concentration}} + \text{central\_size}\\ &\quad * \text{degradation\_rate\_fibrin} * \text{fibrin\_concentration} + \frac{\text{central\_size} * \text{maximal\_degradation\_rate\_plasmin\_fibrin} * \text{plasmin\_concentration} * \text{fibrin\_concentration}}{\text{half\_maximal\_effect\_concentration\_plasmin\_fibrin} + \text{plasmin\_concentration}}\\ &\quad - \text{central\_size} * \text{degradation\_rate\_fdp} * \text{fibrin\_degradation\_product\_concentration}\end{aligned}$$

$$\frac{d}{dt}\text{d\_dimer\_amount} = \text{central\_size} * \text{degradation\_rate\_xf} * \text{cross\_linked\_fibrin\_concentration} + \frac{\text{central\_size} * \text{maximal\_degradation\_rate\_plasmin\_xf} * \text{plasmin\_concentration} * \text{cross\_linked\_fibrin\_concentration}}{\text{half\_maximal\_effect\_concentration\_plasmin\_xf} + \text{plasmin\_concentration}} + \frac{\text{central\_size} * \text{maximal\_degradation\_rate\_apc\_ps\_complex\_xf} * \text{apc\_ps\_complex\_concentration} * \text{cross\_linked\_fibrin\_concentration}}{\text{half\_maximal\_effect\_concentration\_apc\_ps\_complex\_xf} + \text{apc\_ps\_complex\_concentration}} - \text{central\_size} * \text{degradation\_rate\_d\_dimer} * \text{d\_dimer\_concentration}$$

$$\frac{d}{dt}\text{fibrinogen\_amount} = \text{central\_size} * \text{production\_rate\_fibrinogen} - \text{central\_size} * \text{degradation\_rate\_fibrinogen} * \text{fibrinogen\_concentration} - \frac{\text{central\_size} * \text{maximal\_conversion\_rate\_thrombin\_fibrinogen} * \text{activated\_coagulation\_factor\_ii\_concentration} * \text{fibrinogen\_concentration}}{\text{half\_maximal\_effect\_concentration\_thrombin\_fibrinogen} + \text{activated\_coagulation\_factor\_ii\_concentration}} - \frac{\text{central\_size} * \text{maximal\_degradation\_rate\_plasmin\_fibrinogen} * \text{plasmin\_concentration} * \text{fibrinogen\_concentration}}{\text{half\_maximal\_effect\_concentration\_plasmin\_fibrinogen} + \text{plasmin\_concentration}}$$

$$\frac{d}{dt}\text{kallikrein\_amount} = \frac{\text{central\_size} * \text{maximal\_conversion\_rate\_axii\_prekallikrein} * \text{activated\_coagulation\_factor\_xii\_concentration} * \text{prekallikrein\_concentration}}{\text{half\_maximal\_effect\_concentration\_axii\_prekallikrein} + \text{activated\_coagulation\_factor\_xii\_concentration}} - \text{central\_size} * \text{degradation\_rate\_kallikrein} * \text{kallikrein\_concentration}$$

$$\frac{d}{dt}\text{prekallikrein\_amount} = \text{central\_size} * \text{production\_rate\_prekallikrein} - \text{central\_size} * \text{degradation\_rate\_prekallikrein} * \text{prekallikrein\_concentration} - \frac{\text{central\_size} * \text{maximal\_conversion\_rate\_axii\_prekallikrein} * \text{activated\_coagulation\_factor\_xii\_concentration} * \text{prekallikrein\_concentration}}{\text{half\_maximal\_effect\_concentration\_axii\_prekallikrein} + \text{activated\_coagulation\_factor\_xii\_concentration}}$$

$$\begin{aligned}\frac{d}{dt}\text{plasmin\_amount} \\ &= \frac{\text{central\_size} * \text{maximal\_conversion\_rate\_thrombin\_plasminogen} * \text{activated\_coagulation\_factor\_ii\_concentration} * \text{plasminogen\_concentration}}{\text{half\_maximal\_effect\_concentration\_thrombin\_plasminogen} + \text{activated\_coagulation\_factor\_ii\_concentration}} \\ &\quad + \frac{\text{central\_size} * \text{maximal\_conversion\_rate\_fibrin\_plasminogen} * \text{fibrin\_concentration} * \text{plasminogen\_concentration}}{\text{half\_maximal\_effect\_concentration\_fibrin\_plasminogen} + \text{fibrin\_concentration}} \\ &\quad + \frac{\text{central\_size} * \text{maximal\_conversion\_rate\_apc\_ps\_complex\_plasminogen} * \text{apc\_ps\_complex\_concentration} * \text{plasminogen\_concentration}}{\text{half\_maximal\_effect\_concentration\_apc\_ps\_complex\_plasminogen} + \text{apc\_ps\_complex\_concentration}} \\ &\quad - \text{central\_size} * \text{degradation\_rate\_plasmin} * \text{plasmin\_concentration}\end{aligned}$$

$$\begin{aligned}\frac{d}{dt}\text{plasminogen\_amount} \\ &= \text{central\_size} * \text{production\_rate\_plasminogen} - \text{central\_size} * \text{degradation\_rate\_plasminogen} * \text{plasminogen\_concentration} \\ &\quad - \frac{\text{central\_size} * \text{maximal\_conversion\_rate\_thrombin\_plasminogen} * \text{activated\_coagulation\_factor\_ii\_concentration} * \text{plasminogen\_concentration}}{\text{half\_maximal\_effect\_concentration\_thrombin\_plasminogen} + \text{activated\_coagulation\_factor\_ii\_concentration}} \\ &\quad - \frac{\text{central\_size} * \text{maximal\_conversion\_rate\_fibrin\_plasminogen} * \text{fibrin\_concentration} * \text{plasminogen\_concentration}}{\text{half\_maximal\_effect\_concentration\_fibrin\_plasminogen} + \text{fibrin\_concentration}} \\ &\quad - \frac{\text{central\_size} * \text{maximal\_conversion\_rate\_apc\_ps\_complex\_plasminogen} * \text{apc\_ps\_complex\_concentration} * \text{plasminogen\_concentration}}{\text{half\_maximal\_effect\_concentration\_apc\_ps\_complex\_plasminogen} + \text{apc\_ps\_complex\_concentration}}\end{aligned}$$

$$\begin{aligned}\frac{d}{dt}\text{protein\_c\_amount} &= \text{central\_size} * \text{production\_rate\_pc} * \text{vitamin\_k\_hydroquinone\_concentration} \\ &\quad - \text{central\_size} * \text{degradation\_rate\_pc} * \text{protein\_c\_concentration} \\ &\quad - \frac{\text{central\_size} * \text{maximal\_activation\_rate\_aii\_tmod\_complex\_pc} * \text{aii\_tmod\_complex\_concentration} * \text{protein\_c\_concentration}}{\text{half\_maximal\_effect\_concentration\_aii\_tmod\_complex\_pc} + \text{aii\_tmod\_complex\_concentration}}\end{aligned}$$

$$\begin{aligned}\frac{d}{dt}\text{activated\_protein\_c\_amount} &= \frac{\text{central\_size} * \text{maximal\_activation\_rate\_aii\_tmod\_complex\_pc} * \text{aii\_tmod\_complex\_concentration} * \text{protein\_c\_concentration}}{\text{half\_maximal\_effect\_concentration\_aii\_tmod\_complex\_pc} + \text{aii\_tmod\_complex\_concentration}} \\ &\quad - \text{central\_size} * \text{degradation\_rate\_pc\_activated} * \text{activated\_protein\_c\_concentration} - \text{central\_size} \\ &\quad * \text{complex\_formation\_rate\_apc\_ps} * \text{activated\_protein\_c\_concentration} * \text{protein\_s\_concentration}\end{aligned}$$

$$\begin{aligned}\frac{d}{dt}\text{protein\_s\_amount} &= -\text{central\_size} * \text{complex\_formation\_rate\_apc\_ps} * \text{activated\_protein\_c\_concentration} * \text{protein\_s\_concentration} + \text{central\_size} \\ &\quad * \text{production\_rate\_ps} * \text{vitamin\_k\_hydroquinone\_concentration} - \text{central\_size} * \text{degradation\_rate\_ps} * \text{protein\_s\_concentration}\end{aligned}$$

$$\begin{aligned}\frac{d}{dt}\text{thrombomodulin\_amount} = & -\text{central\_size} * \text{complex\_formation\_rate\_aii\_tmod} * \text{activated\_coagulation\_factor\_ii\_concentration} \\ & * \text{thrombomodulin\_concentration} + \text{central\_size} * \text{production\_rate\_tmod} \\ & - \text{central\_size} * \text{degradation\_rate\_tmod} * \text{thrombomodulin\_concentration}\end{aligned}$$

$$\begin{aligned}\frac{d}{dt}\text{tissue\_factor\_amount} = & -\text{central\_size} * \text{complex\_formation\_rate\_vii\_tf} * \text{coagulation\_factor\_vii\_concentration} * \text{tissue\_factor\_concentration} \\ & - \text{central\_size} * \text{complex\_formation\_rate\_avii\_tf} * \text{activated\_coagulation\_factor\_vii\_concentration} \\ & * \text{tissue\_factor\_concentration} - \text{central\_size} * \text{degradation\_rate\_tissue\_factor} * \text{tissue\_factor\_concentration}\end{aligned}$$

$$\begin{aligned}\frac{d}{dt}\text{tissue\_factor\_pathway\_inhibitor\_amount} = & -\text{central\_size} * \text{complex\_formation\_rate\_ax\_tfpi} * \text{activated\_coagulation\_factor\_x\_concentration} \\ & * \text{tissue\_factor\_pathway\_inhibitor\_concentration} + \text{central\_size} * \text{production\_rate\_tfpi} \\ & - \text{central\_size} * \text{degradation\_rate\_tfpi} * \text{tissue\_factor\_pathway\_inhibitor\_concentration}\end{aligned}$$

$$\begin{aligned}\frac{d}{dt}\text{vitamin\_k\_amount} = & \ \text{central\_size} * \text{input\_rate\_vk} - \text{central\_size} * \text{elimination\_rate\_vk} * \text{vitamin\_k\_concentration} - \text{central\_size} * \text{conversion\_rate\_vk\_vkh2} \\ & * \text{vitamin\_k\_concentration} * \left(1.0 - \frac{\text{maximal\_inhibitory\_effect\_warfarin} * \text{warfarin\_concentration}^{\text{gamma}}}{\text{half\_maximal\_effect\_concentration\_warfarin}^{\text{gamma}} + \text{warfarin\_concentration}^{\text{gamma}}}\right) \\ & + \text{central\_size} * \text{conversion\_rate\_vko\_vk} * \text{vitamin\_k\_epoxide\_concentration} \\ & * \left(1.0 - \frac{\text{maximal\_inhibitory\_effect\_warfarin} * \text{warfarin\_concentration}^{\text{gamma}}}{\text{half\_maximal\_effect\_concentration\_warfarin}^{\text{gamma}} + \text{warfarin\_concentration}^{\text{gamma}}}\right) \\ & - \text{central\_size} * \text{transition\_rate\_vk\_central\_to\_peripheral} * \text{vitamin\_k\_concentration} \\ & + \text{peripheral\_vitamin\_k\_size} * \text{transition\_rate\_vk\_peripheral\_to\_central} * \text{vitamin\_k\_peripheral\_concentration}\end{aligned}$$

$$\begin{aligned}\frac{d}{dt}\text{vitamin\_k\_peripheral\_amount} = & \ \text{central\_size} * \text{transition\_rate\_vk\_central\_to\_peripheral} * \text{vitamin\_k\_concentration} \\ & - \text{peripheral\_vitamin\_k\_size} * \text{transition\_rate\_vk\_peripheral\_to\_central} * \text{vitamin\_k\_peripheral\_concentration}\end{aligned}$$

$$\begin{aligned}\frac{d}{dt}\text{vitamin\_k\_epoxide\_amount} = & \ \text{central\_size} * \text{conversion\_rate\_vkh2\_vko} * \text{vitamin\_k\_hydroquinone\_concentration} \\ & - \text{central\_size} * \text{conversion\_rate\_vko\_vk} * \text{vitamin\_k\_epoxide\_concentration} \\ & * \left(1.0 - \frac{\text{maximal\_inhibitory\_effect\_warfarin} * \text{warfarin\_concentration}^{\text{gamma}}}{\text{half\_maximal\_effect\_concentration\_warfarin}^{\text{gamma}} + \text{warfarin\_concentration}^{\text{gamma}}}\right)\end{aligned}$$

$$\frac{d}{dt}\text{vitamin\_k\_hydroquinone\_amount} = \text{central\_size} * \text{conversion\_rate\_vk\_vkh2} * \text{vitamin\_k\_concentration} * \left(1.0 - \frac{\text{maximal\_inhibitory\_effect\_warfarin} * \text{warfarin\_concentration}^{\text{gamma}}}{\text{half\_maximal\_effect\_concentration\_warfarin}^{\text{gamma}} + \text{warfarin\_concentration}^{\text{gamma}}}\right) - \text{central\_size} * \text{conversion\_rate\_vkh2\_vko} * \text{vitamin\_k\_hydroquinone\_concentration}$$

$$\frac{d}{dt}\text{aii\_atiii\_heparin\_complex\_amount} = \text{central\_size} * \text{complex\_formation\_rate\_aii\_atiii\_heparin\_complex} * \text{activated\_coagulation\_factor\_ii\_concentration} * \text{atiii\_heparin\_complex\_concentration}$$

$$\frac{d}{dt}\text{aii\_at\_complex\_amount} = \text{central\_size} * \text{degradation\_rate\_ii\_activated} * \text{activated\_coagulation\_factor\_ii\_concentration} - \text{central\_size} * \text{degradation\_rate\_tat} * \text{aii\_at\_complex\_concentration}$$

$$\frac{d}{dt}\text{aii\_tmod\_complex\_amount} = \text{central\_size} * \text{complex\_formation\_rate\_aii\_tmod} * \text{activated\_coagulation\_factor\_ii\_concentration} * \text{thrombomodulin\_concentration} - \text{central\_size} * \text{degradation\_rate\_aii\_tmod\_complex} * \text{aii\_tmod\_complex\_concentration}$$

$$\frac{d}{dt}\text{av\_ax\_complex\_amount} = \text{central\_size} * \text{complex\_formation\_rate\_av\_ax} * \text{activated\_coagulation\_factor\_v\_concentration} * \text{activated\_coagulation\_factor\_x\_concentration} - \text{central\_size} * \text{degradation\_rate\_av\_ax\_complex} * \text{av\_ax\_complex\_concentration} - \frac{\text{central\_size} * \text{maximal\_degradation\_rate\_apc\_ps\_complex\_av\_ax\_complex} * \text{apc\_ps\_complex\_concentration} * \text{av\_ax\_complex\_concentration}}{\text{half\_maximal\_effect\_concentration\_apc\_ps\_complex\_av\_ax\_complex} + \text{apc\_ps\_complex\_concentration}}$$

$$\frac{d}{dt}\text{vii\_tf\_complex\_amount} = \text{central\_size} * \text{complex\_formation\_rate\_vii\_tf} * \text{coagulation\_factor\_vii\_concentration} * \text{tissue\_factor\_concentration} - \text{central\_size} * \text{degradation\_rate\_vii\_tf\_complex} * \text{vii\_tf\_complex\_concentration} - \frac{\text{central\_size} * \text{maximal\_activation\_rate\_ax\_vii\_tf\_complex} * \text{activated\_coagulation\_factor\_x\_concentration} * \text{vii\_tf\_complex\_concentration}}{\text{half\_maximal\_effect\_concentration\_ax\_vii\_tf\_complex} + \text{activated\_coagulation\_factor\_x\_concentration}} - \frac{\text{central\_size} * \text{maximal\_activation\_rate\_tf\_vii\_tf\_complex} * \text{tissue\_factor\_concentration} * \text{vii\_tf\_complex\_concentration}}{\text{half\_maximal\_effect\_concentration\_tf\_vii\_tf\_complex} + \text{tissue\_factor\_concentration}}$$

$$
\begin{aligned}
&\frac{d}{dt}\text{avii\_tf\_complex\_amount}\\
&\quad= \frac{\text{central\_size} * \text{maximal\_activation\_rate\_ax\_vii\_tf\_complex} * \text{activated\_coagulation\_factor\_x\_concentration} * \text{vii\_tf\_complex\_concentration}}{\text{half\_maximal\_effect\_concentration\_ax\_vii\_tf\_complex} + \text{activated\_coagulation\_factor\_x\_concentration}}\\
&\qquad+ \frac{\text{central\_size} * \text{maximal\_activation\_rate\_tf\_vii\_tf\_complex} * \text{tissue\_factor\_concentration} * \text{vii\_tf\_complex\_concentration}}{\text{half\_maximal\_effect\_concentration\_tf\_vii\_tf\_complex} + \text{tissue\_factor\_concentration}}\\
&\qquad+ \text{central\_size} * \text{complex\_formation\_rate\_avii\_tf} * \text{activated\_coagulation\_factor\_vii\_concentration}\\
&\qquad* \text{tissue\_factor\_concentration} - \text{central\_size} * \text{degradation\_rate\_avii\_tf\_complex} * \text{avii\_tf\_complex\_concentration}\\
&\qquad- \text{central\_size} * \text{complex\_formation\_rate\_avii\_tf\_ax\_tfpi} * \text{avii\_tf\_complex\_concentration} * \text{ax\_tfpi\_complex\_concentration}
\end{aligned}
$$

$$
\begin{aligned}
\frac{d}{dt}\text{avii\_tf\_ax\_tfpi\_complex\_amount} &= \text{central\_size} * \text{complex\_formation\_rate\_avii\_tf\_ax\_tfpi} * \text{avii\_tf\_complex\_concentration} * \text{ax\_tfpi\_complex\_concentration}\\
&\quad- \text{central\_size} * \text{degradation\_rate\_avii\_tf\_ax\_tfpi\_complex} * \text{avii\_tf\_ax\_tfpi\_complex\_concentration}
\end{aligned}
$$

$$
\begin{aligned}
\frac{d}{dt}\text{aix\_aviii\_complex\_amount} &= \text{central\_size} * \text{complex\_formation\_rate\_aix\_aviii} * \text{activated\_coagulation\_factor\_ix\_concentration}\\
&\quad* \text{activated\_coagulation\_factor\_viii\_concentration} - \text{central\_size}\\
&\quad* \text{degradation\_rate\_aix\_aviii\_complex} * \text{aix\_aviii\_complex\_concentration}
\end{aligned}
$$

$$
\begin{aligned}
\frac{d}{dt}\text{aix\_atiii\_heparin\_complex\_amount} &= \text{central\_size} * \text{complex\_formation\_rate\_aix\_atiii\_heparin\_complex}\\
&\quad* \text{activated\_coagulation\_factor\_ix\_concentration} * \text{atiii\_heparin\_complex\_concentration}
\end{aligned}
$$

$$
\begin{aligned}
\frac{d}{dt}\text{ax\_atiii\_heparin\_complex\_complex\_amount} &= \text{central\_size} * \text{complex\_formation\_rate\_ax\_atiii\_heparin\_complex}\\
&\quad* \text{activated\_coagulation\_factor\_x\_concentration} * \text{atiii\_heparin\_complex\_concentration}
\end{aligned}
$$

$$
\begin{aligned}
\frac{d}{dt}\text{ax\_tfpi\_complex\_amount} &= -\text{central\_size} * \text{complex\_formation\_rate\_avii\_tf\_ax\_tfpi} * \text{avii\_tf\_complex\_concentration}\\
&\quad* \text{ax\_tfpi\_complex\_concentration} + \text{central\_size} * \text{complex\_formation\_rate\_ax\_tfpi}\\
&\quad* \text{activated\_coagulation\_factor\_x\_concentration} * \text{tissue\_factor\_pathway\_inhibitor\_concentration}\\
&\quad- \text{central\_size} * \text{degradation\_rate\_ax\_tfpi\_complex} * \text{ax\_tfpi\_complex\_concentration}
\end{aligned}
$$

$$\frac{d}{dt}\text{apc\_ps\_complex\_amount} = \text{central\_size} * \text{complex\_formation\_rate\_apc\_ps} * \text{activated\_protein\_c\_concentration} * \text{protein\_s\_concentration} - \text{central\_size} * \text{degradation\_rate\_apc\_ps\_complex} * \text{apc\_ps\_complex\_concentration}$$

$$\frac{d}{dt}\text{atiii\_heparin\_complex\_amount} = -\text{central\_size} * \text{elimination\_rate\_atiii\_heparin\_complex} * \text{atiii\_heparin\_complex\_concentration} - \text{central\_size} * \text{complex\_formation\_rate\_aii\_atiii\_heparin\_complex} * \text{activated\_coagulation\_factor\_ii\_concentration} * \text{atiii\_heparin\_complex\_concentration} - \text{central\_size} * \text{complex\_formation\_rate\_aix\_atiii\_heparin\_complex} * \text{activated\_coagulation\_factor\_ix\_concentration} * \text{atiii\_heparin\_complex\_concentration} - \text{central\_size} * \text{complex\_formation\_rate\_ax\_atiii\_heparin\_complex} * \text{activated\_coagulation\_factor\_x\_concentration} * \text{atiii\_heparin\_complex\_concentration}$$

$$\frac{d}{dt}\text{drug\_amount} = -\text{absorption\_rate} * \text{drug\_amount} + \text{dose\_rate}$$

$$\frac{d}{dt}\text{warfarin\_amount} = -\text{central\_warfarin\_size} * \text{elimination\_rate\_warfarin} * \text{warfarin\_concentration} + \text{absorption\_rate} * \text{drug\_amount}$$

$$\text{coagulation\_factor\_ii\_concentration} = \frac{\text{coagulation\_factor\_ii\_amount}}{\text{central\_size}}$$

$$\text{activated\_coagulation\_factor\_ii\_concentration} = \frac{\text{activated\_coagulation\_factor\_ii\_amount}}{\text{central\_size}}$$

$$\text{coagulation\_factor\_v\_concentration} = \frac{\text{coagulation\_factor\_v\_amount}}{\text{central\_size}}$$

$$\text{activated\_coagulation\_factor\_v\_concentration} = \frac{\text{activated\_coagulation\_factor\_v\_amount}}{\text{central\_size}}$$

$$\text{coagulation\_factor\_vii\_concentration} = \frac{\text{coagulation\_factor\_vii\_amount}}{\text{central\_size}}$$

$$\text{activated\_coagulation\_factor\_vii\_concentration} = \frac{\text{activated\_coagulation\_factor\_vii\_amount}}{\text{central\_size}}$$

$$\text{coagulation\_factor\_viii\_concentration} = \frac{\text{coagulation\_factor\_viii\_amount}}{\text{central\_size}}$$

$$\text{activated\_coagulation\_factor\_viii\_concentration} = \frac{\text{activated\_coagulation\_factor\_viii\_amount}}{\text{central\_size}}$$

$$\text{coagulation\_factor\_ix\_concentration} = \frac{\text{coagulation\_factor\_ix\_amount}}{\text{central\_size}}$$

$$\text{activated\_coagulation\_factor\_ix\_concentration} = \frac{\text{activated\_coagulation\_factor\_ix\_amount}}{\text{central\_size}}$$

$$\text{coagulation\_factor\_x\_concentration} = \frac{\text{coagulation\_factor\_x\_amount}}{\text{central\_size}}$$

$$\text{activated\_coagulation\_factor\_x\_concentration} = \frac{\text{activated\_coagulation\_factor\_x\_amount}}{\text{central\_size}}$$

$$\text{coagulation\_factor\_xi\_concentration} = \frac{\text{coagulation\_factor\_xi\_amount}}{\text{central\_size}}$$

$$\text{activated\_coagulation\_factor\_xi\_concentration} = \frac{\text{activated\_coagulation\_factor\_xi\_amount}}{\text{central\_size}}$$

$$\text{coagulation\_factor\_xii\_concentration} = \frac{\text{coagulation\_factor\_xii\_amount}}{\text{central\_size}}$$

$$\mathrm{activated\_coagulation\_factor\_xii\_concentration} = \frac{\mathrm{activated\_coagulation\_factor\_xii\_amount}}{\mathrm{central\_size}}$$

$$\mathrm{coagulation\_factor\_xiii\_concentration} = \frac{\mathrm{coagulation\_factor\_xiii\_amount}}{\mathrm{central\_size}}$$

$$\mathrm{activated\_coagulation\_factor\_xiii\_concentration} = \frac{\mathrm{activated\_coagulation\_factor\_xiii\_amount}}{\mathrm{central\_size}}$$

$$\mathrm{contact\_system\_activator\_concentration} = \frac{\mathrm{contact\_system\_activator\_amount}}{\mathrm{central\_size}}$$

$$\mathrm{fibrin\_concentration} = \frac{\mathrm{fibrin\_amount}}{\mathrm{central\_size}}$$

$$\mathrm{cross\_linked\_fibrin\_concentration} = \frac{\mathrm{cross\_linked\_fibrin\_amount}}{\mathrm{central\_size}}$$

$$\mathrm{fibrin\_degradation\_product\_concentration} = \frac{\mathrm{fibrin\_degradation\_product\_amount}}{\mathrm{central\_size}}$$

$$\mathrm{d\_dimer\_concentration} = \frac{\mathrm{d\_dimer\_amount}}{\mathrm{central\_size}}$$

$$\mathrm{fibrinogen\_concentration} = \frac{\mathrm{fibrinogen\_amount}}{\mathrm{central\_size}}$$

$$\mathrm{kallikrein\_concentration} = \frac{\mathrm{kallikrein\_amount}}{\mathrm{central\_size}}$$

$$\text{prekallikrein\_concentration} = \frac{\text{prekallikrein\_amount}}{\text{central\_size}}$$

$$\text{plasmin\_concentration} = \frac{\text{plasmin\_amount}}{\text{central\_size}}$$

$$\text{plasminogen\_concentration} = \frac{\text{plasminogen\_amount}}{\text{central\_size}}$$

$$\text{protein\_c\_concentration} = \frac{\text{protein\_c\_amount}}{\text{central\_size}}$$

$$\text{activated\_protein\_c\_concentration} = \frac{\text{activated\_protein\_c\_amount}}{\text{central\_size}}$$

$$\text{protein\_s\_concentration} = \frac{\text{protein\_s\_amount}}{\text{central\_size}}$$

$$\text{thrombomodulin\_concentration} = \frac{\text{thrombomodulin\_amount}}{\text{central\_size}}$$

$$\text{tissue\_factor\_concentration} = \frac{\text{tissue\_factor\_amount}}{\text{central\_size}}$$

$$\text{tissue\_factor\_pathway\_inhibitor\_concentration} = \frac{\text{tissue\_factor\_pathway\_inhibitor\_amount}}{\text{central\_size}}$$

$$\text{vitamin\_k\_concentration} = \frac{\text{vitamin\_k\_amount}}{\text{central\_size}}$$

$$\text{vitamin\_k\_peripheral\_concentration} = \frac{\text{vitamin\_k\_peripheral\_amount}}{\text{peripheral\_vitamin\_k\_size}}$$

$$\text{vitamin\_k\_epoxide\_concentration} = \frac{\text{vitamin\_k\_epoxide\_amount}}{\text{central\_size}}$$

$$\text{vitamin\_k\_hydroquinone\_concentration} = \frac{\text{vitamin\_k\_hydroquinone\_amount}}{\text{central\_size}}$$

$$\text{aii\_atiii\_heparin\_complex\_concentration} = \frac{\text{aii\_atiii\_heparin\_complex\_amount}}{\text{central\_size}}$$

$$\text{aii\_at\_complex\_concentration} = \frac{\text{aii\_at\_complex\_amount}}{\text{central\_size}}$$

$$\text{aii\_tmod\_complex\_concentration} = \frac{\text{aii\_tmod\_complex\_amount}}{\text{central\_size}}$$

$$\text{av\_ax\_complex\_concentration} = \frac{\text{av\_ax\_complex\_amount}}{\text{central\_size}}$$

$$\text{vii\_tf\_complex\_concentration} = \frac{\text{vii\_tf\_complex\_amount}}{\text{central\_size}}$$

$$\text{avii\_tf\_complex\_concentration} = \frac{\text{avii\_tf\_complex\_amount}}{\text{central\_size}}$$

$$\text{avii\_tf\_ax\_tfpi\_complex\_concentration} = \frac{\text{avii\_tf\_ax\_tfpi\_complex\_amount}}{\text{central\_size}}$$

$$\text{aix\_aviii\_complex\_concentration} = \frac{\text{aix\_aviii\_complex\_amount}}{\text{central\_size}}$$

$$\text{aix\_atiii\_heparin\_complex\_concentration} = \frac{\text{aix\_atiii\_heparin\_complex\_amount}}{\text{central\_size}}$$

$$\text{ax\_atiii\_heparin\_complex\_complex\_concentration} = \frac{\text{ax\_atiii\_heparin\_complex\_complex\_amount}}{\text{central\_size}}$$

$$\text{ax\_tfpi\_complex\_concentration} = \frac{\text{ax\_tfpi\_complex\_amount}}{\text{central\_size}}$$

$$\text{apc\_ps\_complex\_concentration} = \frac{\text{apc\_ps\_complex\_amount}}{\text{central\_size}}$$

$$\text{atiii\_heparin\_complex\_concentration} = \frac{\text{atiii\_heparin\_complex\_amount}}{\text{central\_size}}$$

$$\text{warfarin\_concentration} = \frac{\text{warfarin\_amount}}{\text{central\_warfarin\_size}}$$

**Table S1.2.** States of the mechanistic model

| |
|---|
| atiii_heparin_complex_amount |
| coagulation_factor_ii_amount |
| activated_coagulation_factor_ii_amount |
| aii_at_complex_amount |
| aii_atiii_heparin_complex_amount |
| thrombomodulin_amount |
| aii_tmod_complex_amount |
| coagulation_factor_v_amount |
| activated_coagulation_factor_v_amount |
| activated_coagulation_factor_x_amount |
| av_ax_complex_amount |
| coagulation_factor_vii_amount |
| activated_coagulation_factor_vii_amount |
| tissue_factor_amount |
| vii_tf_complex_amount |
| avii_tf_complex_amount |

ax_tfpi_complex_amount

avii_tf_ax_tfpi_complex_amount

coagulation_factor_viii_amount

activated_coagulation_factor_viii_amount

coagulation_factor_ix_amount

activated_coagulation_factor_ix_amount

aix_aviii_complex_amount

aix_atiii_heparin_complex_amount

coagulation_factor_x_amount

ax_atiii_heparin_complex_complex_amount

tissue_factor_pathway_inhibitor_amount

coagulation_factor_xi_amount

activated_coagulation_factor_xi_amount

coagulation_factor_xii_amount

activated_coagulation_factor_xii_amount

coagulation_factor_xiii_amount

activated_coagulation_factor_xiii_amount

contact_system_activator_amount

fibrinogen_amount

fibrin_degradation_product_amount

fibrin_amount

cross_linked_fibrin_amount

d_dimer_amount

prekallikrein_amount

kallikrein_amount

plasminogen_amount

plasmin_amount

protein_c_amount

activated_protein_c_amount

protein_s_amount

apc_ps_complex_amount

vitamin_k_amount

vitamin_k_hydroquinone_amount

vitamin_k_epoxide_amount

vitamin_k_peripheral_amount

warfarin_amount

drug_amount

**Table S1.3.** Parameters of the mechanistic model (Parameters with the same name as a state refer to the initial condition of the state)

| | |
|---|---|
| degradation_rate_ii | |
| degradation_rate_ii_activated | |
| half_maximal_effect_concentration_av_ax_complex_ii | |
| half_maximal_effect_concentration_ax_ii | |
| maximal_activation_rate_av_ax_complex_ii | |
| maximal_activation_rate_ax_ii | |
| production_rate_ii | (†) |
| degradation_rate_v | |
| degradation_rate_v_activated | |
| half_maximal_effect_concentration_thrombin_v | |
| half_maximal_effect_concentration_apc_ps_complex_av | |
| maximal_activation_rate_thrombin_v | |
| maximal_degradation_rate_apc_ps_complex_av | |
| production_rate_v | (††) |
| degradation_rate_vii | |
| degradation_rate_vii_activated | |

half_maximal_effect_concentration_thrombin_vii

half_maximal_effect_concentration_aix_vii

half_maximal_effect_concentration_ax_vii

half_maximal_effect_concentration_avii_tf_complex_vii

maximal_activation_rate_thrombin_vii

maximal_activation_rate_aix_vii

maximal_activation_rate_ax_vii

maximal_activation_rate_avii_tf_complex_vii

production_rate_vii (†)

degradation_rate_viii

degradation_rate_viii_activated

half_maximal_effect_concentration_thrombin_viii

half_maximal_effect_concentration_apc_ps_complex_aviii

maximal_activation_rate_thrombin_viii

maximal_degradation_rate_apc_ps_complex_aviii

production_rate_viii (†)

degradation_rate_ix

degradation_rate_ix_activated

half_maximal_effect_concentration_avii_tf_complex_ix

half_maximal_effect_concentration_axi_ix

maximal_activation_rate_avii_tf_complex_ix

maximal_activation_rate_axi_ix

production_rate_ix (†)

degradation_rate_x

degradation_rate_x_activated

half_maximal_effect_concentration_avii_x

half_maximal_effect_concentration_avii_tf_complex_x

half_maximal_effect_concentration_aix_x

half_maximal_effect_concentration_aix_aviii_complex_x

maximal_activation_rate_avii_x

maximal_activation_rate_avii_tf_complex_x

maximal_activation_rate_aix_x

maximal_activation_rate_aix_aviii_complex_x

production_rate_x (†)

degradation_rate_xi

degradation_rate_xi_activated

half_maximal_effect_concentration_axii_xi

half_maximal_effect_concentration_thrombin_xi

maximal_activation_rate_axii_xi

maximal_activation_rate_thrombin_xi

production_rate_xi (††)

degradation_rate_xii

degradation_rate_xii_activated

half_maximal_effect_concentration_ca_xii

half_maximal_effect_concentration_kallikrein_xii

maximal_activation_rate_ca_xii

maximal_activation_rate_kallikrein_xii

production_rate_xii (††)

degradation_rate_xiii

degradation_rate_xiii_activated

half_maximal_effect_concentration_thrombin_xiii

maximal_activation_rate_thrombin_xiii

production_rate_xiii (††)

degradation_rate_ca

degradation_rate_fibrinogen

degradation_rate_fibrin

degradation_rate_xf

half_maximal_effect_concentration_axiii_fibrin

half_maximal_effect_concentration_thrombin_fibrinogen

half_maximal_effect_concentration_plasmin_fibrinogen

half_maximal_effect_concentration_plasmin_fibrin

half_maximal_effect_concentration_plasmin_xf

half_maximal_effect_concentration_apc_ps_complex_xf

maximal_conversion_rate_axiii_fibrin

maximal_conversion_rate_thrombin_fibrinogen

maximal_degradation_rate_plasmin_fibrinogen

maximal_degradation_rate_plasmin_fibrin

maximal_degradation_rate_plasmin_xf

maximal_degradation_rate_apc_ps_complex_xf

production_rate_fibrinogen (†)

degradation_rate_fdp

degradation_rate_d_dimer

degradation_rate_kallikrein

degradation_rate_prekallikrein

half_maximal_effect_concentration_axii_prekallikrein

maximal_conversion_rate_axii_prekallikrein

production_rate_prekallikrein (†)

degradation_rate_plasminogen

degradation_rate_plasmin

half_maximal_effect_concentration_apc_ps_complex_plasminogen

half_maximal_effect_concentration_fibrin_plasminogen

half_maximal_effect_concentration_thrombin_plasminogen

maximal_conversion_rate_apc_ps_complex_plasminogen

maximal_conversion_rate_fibrin_plasminogen

maximal_conversion_rate_thrombin_plasminogen

production_rate_plasminogen (†)

degradation_rate_pc

degradation_rate_pc_activated

half_maximal_effect_concentration_aii_tmod_complex_pc

maximal_activation_rate_aii_tmod_complex_pc

production_rate_pc (†)

degradation_rate_ps

production_rate_ps (†)

degradation_rate_tmod

production_rate_tmod (†)

degradation_rate_tissue_factor

conversion_rate_vk_vkh2

conversion_rate_vkh2_vko

conversion_rate_vko_vk

elimination_rate_vk

half_maximal_effect_concentration_warfarin

maximal_inhibitory_effect_warfarin

transition_rate_vk_central_to_peripheral (†)

transition_rate_vk_peripheral_to_central (†)

input_rate_vk (†)

elimination_rate_warfarin

gamma

degradation_rate_tfpi

production_rate_tfpi (†)

degradation_rate_tat

complex_formation_rate_aii_atiii_heparin_complex

complex_formation_rate_aii_tmod

degradation_rate_aii_tmod_complex

complex_formation_rate_av_ax

degradation_rate_av_ax_complex

half_maximal_effect_concentration_apc_ps_complex_av_ax_complex

maximal_degradation_rate_apc_ps_complex_av_ax_complex

complex_formation_rate_vii_tf

degradation_rate_vii_tf_complex

half_maximal_effect_concentration_ax_vii_tf_complex

half_maximal_effect_concentration_tf_vii_tf_complex

maximal_activation_rate_ax_vii_tf_complex

maximal_activation_rate_tf_vii_tf_complex

complex_formation_rate_avii_tf

degradation_rate_avii_tf_complex

complex_formation_rate_aix_atiii_heparin_complex

complex_formation_rate_aix_aviii

degradation_rate_aix_aviii_complex

complex_formation_rate_ax_atiii_heparin_complex

complex_formation_rate_ax_tfpi

degradation_rate_ax_tfpi_complex

complex_formation_rate_apc_ps

degradation_rate_apc_ps_complex

complex_formation_rate_avii_tf_ax_tfpi

degradation_rate_avii_tf_ax_tfpi_complex

elimination_rate_atiii_heparin_complex

atiii_heparin_complex_amount

coagulation_factor_ii_amount

activated_coagulation_factor_ii_amount

aii_at_complex_amount

aii_atiii_heparin_complex_amount

thrombomodulin_amount

aii_tmod_complex_amount

coagulation_factor_v_amount

activated_coagulation_factor_v_amount

activated_coagulation_factor_x_amount

av_ax_complex_amount

coagulation_factor_vii_amount

activated_coagulation_factor_vii_amount

tissue_factor_amount

vii_tf_complex_amount

avii_tf_complex_amount

ax_tfpi_complex_amount

avii_tf_ax_tfpi_complex_amount

coagulation_factor_viii_amount

activated_coagulation_factor_viii_amount

coagulation_factor_ix_amount

activated_coagulation_factor_ix_amount

aix_aviii_complex_amount

aix_atiii_heparin_complex_amount

coagulation_factor_x_amount

ax_atiii_heparin_complex_complex_amount

tissue_factor_pathway_inhibitor_amount

coagulation_factor_xi_amount

activated_coagulation_factor_xi_amount

coagulation_factor_xii_amount

activated_coagulation_factor_xii_amount

coagulation_factor_xiii_amount

activated_coagulation_factor_xiii_amount

contact_system_activator_amount

fibrinogen_amount

fibrin_degradation_product_amount

fibrin_amount

cross_linked_fibrin_amount

d_dimer_amount

prekallikrein_amount

kallikrein_amount

plasminogen_amount

plasmin_amount

protein_c_amount

activated_protein_c_amount

protein_s_amount

apc_ps_complex_amount

vitamin_k_amount

vitamin_k_hydroquinone_amount

vitamin_k_epoxide_amount

vitamin_k_peripheral_amount (†)

warfarin_amount

drug_amount (†)

#### *Appendix S2*

### S2 MODEL 1: NEURAL NETWORK REGRESSION MODEL

#### S2.1 Background

This MIPD method assumes that inter-individual differences of warfarin maintenance doses can be predicted based on covariates of the treatment response variability

$$d^* = d^*(\chi, y^*). \tag{S2.1}$$

$d^*$ denotes the maintenance dose, $\chi$ denotes the covariates and $y^*$ denotes the target treatment response. This assumed functional relationship between $d^*$, $\chi$ and $y^*$ is unknown, but may be learned from data, e.g. using a neural network (Hornik et al., 1989).

#### S2.2 Implementation

In our implementation of the regression model, we approximate the function $d^*$ with a neural network implemented in `PyTorch` (Paszke et al., 2019)

$$d^*(\chi, y^*) \approx f_\theta(\chi, y^*), \tag{S2.2}$$

where $f$ denotes the neural network and $\theta$ denotes its parameters. The architecture of the network is a modelling choice. Our network is composed of three fully connected layers with nonlinear activations

$$f_\theta(\chi, y^*) = f^{(3)}_{\theta_3}\left(f^{(2)}_{\theta_2}\left(f^{(1)}_{\theta_1}(\chi, y^*)\right)\right). \tag{S2.3}$$

The parameters of $f$ are the parameters of the individual layers, $\theta = (\theta_1, \theta_2, \theta_3)$. Layer 1 takes as input the covariates and the target treatment response encoded by five input features, $\mathbf{x} = (a_G, a_{*1}, a_{*2}, t_\star, y^*)$, where $a_G$ denotes the number of G VKORC1 alleles, $a_{*1}$ and $a_{*2}$ denote the number of $*1$ and $*2$ CYP2C9 alleles and $t_\star$ denotes the age. The first 4 features in $\mathbf{x}$ uniquely define the covariates, $\chi$. The output of layer 1 are 1024 hidden features obtained by applying a rectified linear unit (ReLU) activation (Agarap, 2018) to the matrix product of the input features and a weight matrix, $\mathbf{W}$, of shape $(1024, 5)$

$$f^{(1)}(\chi, y^*) = \text{ReLU}(\mathbf{W}\mathbf{x}). \tag{S2.4}$$

The parameters of layer 1, $\theta_1$, are the elements of the weight matrix and the parameters of the ReLU activation function.

Layer 2 is also a fully connected layer with ReLU activation and takes as inputs the hidden features of layer 1

$$f^{(2)}(f^{(1)}) = \text{ReLU}(\mathbf{W} f^{(1)}). \tag{S2.5}$$

The weight matrix of layer 2 is of shape $(1024, 1024)$, resulting in an output of 1024 hidden features. The parameters of layer 2, $\theta_2$, are the elements of the weight matrix and the parameters of the ReLU activation function.

Layer 3 is the final layer of the network, taking the hidden features from layer 2 as an input

$$f^{(3)}(f^{(2)}) = \text{Sigmoid}(\mathbf{W} f^{(2)}). \tag{S2.6}$$

The weight matrix of layer 3 is of shape $(1, 1024)$. In contrast to layer 1 and 2, the final layer has a sigmoid activation, constraining the output values of the network to the interval between $0$ and $1$. The parameters of layer 3, $\theta_3$, are the elements of the weight matrix and the parameters of the sigmoid activation function.

For computational stability, both the input and the output of the network are normalised. The allele counts, $(a_G, a_{*1}, a_{*2})$, are normalised by the maximum allele count, 2. The age, $t_\star$, is centered around $51$ and normalised by $15$, approximating the typical age and the typical standard deviation of the age in a population. The target INR, $y^*$, is centered at $2.5$ and normalised by $2.5$, motivated by the therapeutic range of warfarin. The output of the network is already normalised to $0$ and $1$ by virtue of the sigmoid activation function.

#### S2.3 Dose prediction

For maintenance dose predictions, the model output needs to be translated into dosages of warfarin. We do this by assuming a maximally feasible maintenance dose of $30\,\text{mg}$. This makes it possible to interpret the network output as the predicted maintenance dose in units of the maximal maintenance dose, such that predicted maintenance doses may be obtained by multiplying the model output with $30\,\text{mg}$. Notably, this constrains the model predictions to doses between $0\,\text{mg}$ and $30\,\text{mg}$. For dose predictions during the MIPD trial, the network output is further constrained to doses that can be administered in terms of combinations of commercially available warfarin tablets, namely $0\,\text{mg}$, $1\,\text{mg}$ and any dose $\geq 2\,\text{mg}$ in steps of $0.5\,\text{mg}$. The continuous output of the model is mapped to these dosages by rounding the output to the nearest dose.

#### S2.4 Fitting

We train the model on the simulated clinical phase III data, introduced in Simulating trial phases prior to MIPD. The data includes monitoring data, covariates and warfarin dosing regimens for 1000 patients simulated with the warfarin trial model (see Appendix S1). The data used for training are the covariates, the INR measurements and the maintenance warfarin dosages at the end of the trial. We split the data randomly into a train set $(90\,\%)$ and into a test set $(10\,\%)$. The inputs and the dosages are normalised as outlined above. The model is trained for 1000 epochs to minimise the mean squared error objective function using Adam. We use a batch size of 128 and a learning rate of $10^{-4}$. The learning rate is reduced by a factor of 2 every time the objective function does not improve for 10 consecutive epochs until a minimum learning rate of $10^{-7}$ is reached. The training results are shown in Fig S2.1.

The figure shows the objective function evaluated on the training dataset and the test dataset. The epoch with the smallest test error is highlighted with a dashed line. We use this 'best test error' model for MIPD in the remainder of the article.

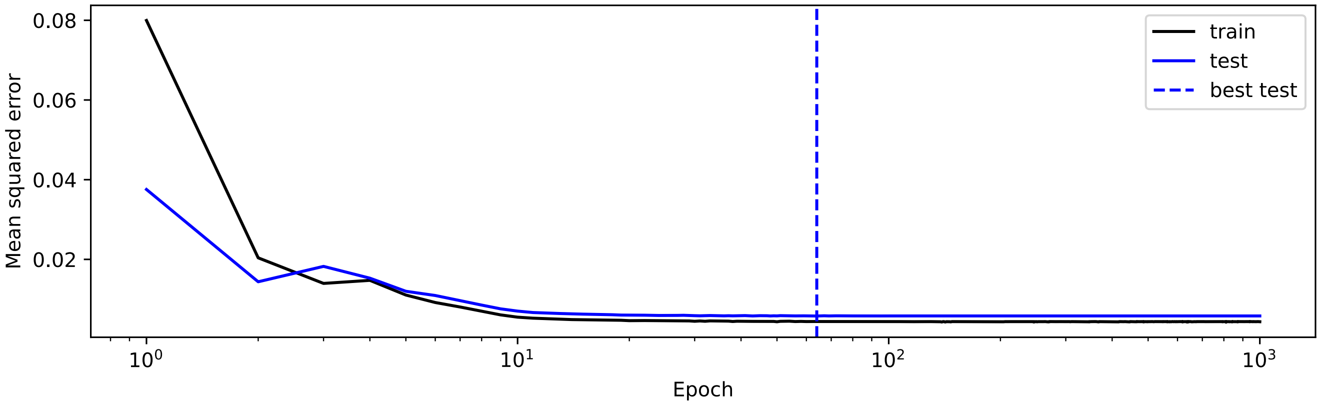


**Figure S2.1. Training of model 1:** The figure shows the mean squared error of the model predictions on the train set (black) and the test set (blue). The epoch with the lowest mean squared error on the test set is indicated by a dashed line.

#### *Appendix S3*

### S3 MODEL 2: DEEP REINFORCEMENT LEARNING MODEL

#### S3.1 Background

This MIPD method is a reinforcement learning approach which assumes that dose decisions can be modelled as a Markov decision process defined by a state space, $\mathcal{S}$, an action space, $\mathcal{A}$, and stochastic transition dynamics, $p(s_{j+1}|s_j, a_j)$ (Zadeh et al., 2023). $s_j \in \mathcal{S}$ denotes the state at time $t_j = j\Delta t$, where $\Delta t$ is the temporal resolution of the model, and $a_j \in \mathcal{A}$ denotes the action at time $t_j$. In our case, we define the state of an individual by the individual's covariates and INR monitoring measurement, $s_j = (\chi, y_j)$. The actions are the different warfarin doses that may be administered, $a_j = d_j$, and the transition dynamics are the treatment response dynamics following a dose administration. We choose a temporal resolution of the model of one day, $\Delta t = 1$ day, since warfarin doses are administered daily. The Markov decision process assumption makes it possible to derive optimal dose decisions based only on the current treatment response of an individual (Bellman, 1966), as discussed in more detail below.

In reinforcement learning optimality is defined using a reward function, $r(s)$. The reward function is a modelling choice and quantifies the desirability of a state. In our case, we define the reward function using the negative mean squared error of the INR measurement, $y$, to the target treatment response, $y^*$,

$$r(s) = -\left(y - y^*\right)^2 . \tag{S3.1}$$

We reward INR measurements close to the target INR, and penalise INR measurements away from the target INR.

Using the reward function, we can retrospectively quantify the 'return' of a sequence of actions, $(a_0, \ldots, a_n)$, using the accumulation of the rewards associated with the states, $(s_1, \ldots, s_{n+1})$, that followed the actions

$$R(a_0, \ldots, a_n, s_1, \ldots, s_{n+1}) = \sum_{j=1}^{n+1} r(s_j). \tag{S3.2}$$

In reinforcement learning, this retrospective evaluation is converted into a prospective evaluation using a policy, $\pi(a|s)$, which formalises the way actions are chosen. The policy defines the probability of taking action $a$ given that the system is currently in state $s$. This makes it possible to estimate the future return or 'value' of choosing an action $a$ when being in state $s$, assuming that all following actions are chosen according to the policy

$$Q^{\pi}(s,a) = \mathbb{E}\left[\sum_{j=1}^{\infty} \gamma^{j-1} r(s_j) \,\middle|\, s, a\right] \tag{S3.3}$$

We use $j$ to reference the time relative to the current state. For the prospective evaluation, it is customary to discount the contributions from states further into the future with a discount factor, $\gamma < 1$, as those states are associated with a larger degree of uncertainty. The expectation is taken with respect to all possible future states and actions, distributed according to the transition dynamics and the policy. Note that the Markov decision process assumption significantly simplifies this expectation, as neither the transition dynamics nor the policy need to take past states and actions into account.

The action-value function, $Q^{\pi}(s,a)$, provides a simple scoring system for actions – the action with the largest function value has the largest expected future return. The value of the function is also referred to as the Q-value. The optimal policy, $\pi^*$, is defined as the policy that always chooses the action with the largest expected future return

$$\pi^*(a|s) = \begin{cases} 1 & \text{for} \quad \underset{a}{\text{argmax}}\, Q^{\pi^*}(s,a) \\ 0 & \text{else.} \end{cases} \tag{S3.4}$$

This optimal policy is independent of time, previous actions and previous states – a consequence of the Markov property of the decision process. In the context of MIPD, this means that reinforcement learning can be used to define doses as a function of the current state of an individual

$$d_j = f(\chi, y_j), \tag{S3.5}$$

where the function $f$ is independent of time, previous treatment response measurements and previous dose administrations. Under the Markov assumption, these doses optimally target the desired treatment response, Eq S3.1.

While the action-value function, $Q^{\pi^*}$, cannot be computed exactly without knowing the transition dynamics, Watkins and Dayan (1992) showed that $Q^{\pi^*}$ can be learned from data iteratively without a detailed understanding of the dynamics of the system

$$Q_{n+1}(s_k, a_k) = (1-\alpha)Q_n(s_k, a_k) + \alpha(r(s'_k) + \gamma \max_{a'_k} Q_n(s'_k, a'_k)). \tag{S3.6}$$

This learning algorithm is known as Q-learning. $n$ denotes the iteration, $\alpha < 1$ denotes the learning rate, $s_k$ denotes an observed state of the system, $a_k$ denotes the action taken and $r(s'_k)$ denotes the reward associated with the state observed after taking action $a_k$. For a sufficiently large dataset, $\mathcal{D} = \{(s_k, a_k, s'_k)\}$, the estimate, $Q_n$, converges to the exact action-value function associated with the optimal policy, $\lim_{n\to\infty} Q_n = Q^{\pi^*}$.

However, a challenge with Eq S3.6 is that each iteration only updates the Q-value of the observed state-action pair, $(s_k, a_k)$, making the learning algorithm slow and incapable of generalising Q-values beyond the state-action pairs contained in the dataset. To address these limitations, Mnih et al. (2013) introduced the deep Q network (DQN) modelling approach, in which the action-value function is approximated using a neural network

$$Q_n(s, a) \approx f_\theta(s, a). \tag{S3.7}$$

$f$ denotes the neural network and $\theta$ denotes its parameters. This converts the learning task to a function approximation task, where each state-action pair in the dataset informs the parameters of the model. As a result, the learning algorithm becomes more efficient and more capable of generalising Q-values to unobserved state-action pairs.

The network is trained by translating the Q-learning algorithm into an objective function. Mnih et al. (2013) do this using the temporal difference error

$$\Delta_k(\theta) = \left| r(s'_k) + \gamma \max_{a'_k} f_\theta(s'_k, a'_k) - f_\theta(s_k, a_k) \right|. \tag{S3.8}$$

The temporal difference error is the difference between the current Q-value estimate in Eq S3.6, $Q_n(s_k, a_k)$, and the update, $r(s'_k) + \gamma \max_{a'_k} Q_n(s'_k, a'_k)$, where we replaced the action-value function by the neural network approximation. The temporal difference error vanishes for the action-value function associated with the optimal policy, $Q^{\pi^*}$. By minimising the temporal difference error, a neural network can be trained to approximate $Q^{\pi^*}$.

#### S3.2 Implementation

In our implementation of the DQN model, we closely follow Mnih et al. (2013) and use a neural network to predict the Q-values for all possible actions simultaneously

$$\mathbf{Q}^{\pi^*}(s) \approx f_\theta(s), \tag{S3.9}$$

where $\mathbf{Q}^{\pi^*}(s)$ is a vector containing the Q-values for all possible actions when being in state $s$, $\mathbf{Q}^{\pi^*}(s) = [Q^{\pi^*}(s, a_1), \ldots, Q^{\pi^*}(s, a_n)]$. In our case, the actions, $(a_1, \ldots, a_n)$, are the daily warfarin dosages that can be administered as a combination of commercially available warfarin tablets, namely 0 mg, 1 mg and any dose $\geq 2$ mg in 0.5 mg increments. We constrain dosages to a maximum of 30 mg, resulting in a total of $n = 58$ possible dosages.

We implement the neural network in `PyTorch` (Paszke et al., 2019) using four fully connected layers with nonlinear activation functions

$$f_\theta(s) = f^{(4)}_{\theta_4}\left( f^{(3)}_{\theta_3}\left( f^{(2)}_{\theta_2}\left( f^{(1)}_{\theta_1}(\chi, y)\right)\right)\right). \tag{S3.10}$$

The parameters of $f$ are the parameters of the individual layers, $\theta = (\theta_1, \theta_2, \theta_3, \theta_4)$. Layer 1 takes as input the state of the system, i.e. an individual's covariates and the treatment response measurement, encoded by five input features, $\mathbf{x} = (a_G, a_{*1}, a_{*2}, t_\star, y)$, where $a_G$ denotes the number of G VKORC1 alleles, $a_{*1}$ and $a_{*2}$ denote the number of $*1$ and $*2$ CYP2C9 alleles and $t_\star$ denotes the age. The first 4 features in $\mathbf{x}$ uniquely define the covariates, $\chi$. The outputs of layer 1 are 256 hidden features obtained by applying a rectified linear unit (ReLU) activation (Agarap, 2018) to the matrix product of the input features and a weight matrix, $\mathbf{W}$, of shape $(256, 5)$

$$f^{(1)}(\chi, y^*) = \mathrm{ReLU}(\mathbf{W}\mathbf{x}). \tag{S3.11}$$

The parameters of layer 1, $\theta_1$, are the elements of the weight matrix and the parameters of the ReLU activation function.

Layer 2 also uses a ReLU activation and takes as inputs the hidden features of layer 1

$$f^{(2)}(f^{(1)}) = \mathrm{ReLU}(\mathbf{W}f^{(1)}). \tag{S3.12}$$

The weight matrix of layer 2 is of shape $(128, 256)$, resulting in an output of 128 hidden features. The parameters of layer 2, $\theta_2$, are the elements of the weight matrix and the parameters of the ReLU activation function.

Layer 3 has a similar architecture as layer 2 but takes as input the outputs from layer 2

$$f^{(3)}(f^{(2)}) = \mathrm{ReLU}(\mathbf{W}f^{(2)}). \tag{S3.13}$$

The weight matrix of layer 3 is of shape $(64, 128)$, resulting in an output of 64 hidden features. The parameters of layer 3, $\theta_3$, are the elements of the weight matrix and the parameters of the ReLU activation function.

Layer 4 is the final layer of the network, taking the hidden features from layer 3 as input

$$f^{(4)}(f^{(3)}) = \mathbf{W}f^{(3)}. \tag{S3.14}$$

The weight matrix of layer 4 is of shape $(58, 64)$, resulting in 58 outputs of the network. These 58 outputs are the predicted Q-values for the 58 possible warfarin dosages. In contrast to the other layers, the final layer uses no activation function.

For computational stability the input to the network is normalised. The allele counts, $(a_G, a_{*1}, a_{*2})$, are normalised by the maximum allele count, 2. The age, $t_\star$, is centered around $51$ and normalised by $15$, approximating the typical age and the typical standard deviation of the age in a population. The INR measurement, $y$, is centered at $2.5$ and normalised by $2.5$, motivated by the therapeutic range of warfarin.

#### S3.3 Dose prediction

For dose predictions, the predicted Q-values need to be translated into dosages of warfarin. We do this following the optimal policy (see Eq S3.4), and administer the dose with the maximal Q-value. During the MIPD trial, we predict individualised dosing regimens iteratively using the current INR measurement and the covariates of an individual.

#### S3.4 Fitting

Following Zadeh et al. (2023), we train the model not directly on trial data, but on treatment responses simulated with a PKPD model that is calibrated to trial data. This is necessary because standard DQN models are trained online (Ribba et al., 2022), making it challenging to use data from finalised clinical trials for training, although offline reinforcement learning is an active field of research (Agarwal et al., 2020). The PKPD model we choose for the simulation is Hamberg et al's model (Hamberg et al., 2010), which we also use for the PKPD model-based MIPD approach (see MIPD methods). For details on the calibration of the PKPD model, we refer to Appendix S4.

We train the model for 1500 epochs with a learning rate of $10^{-4}$, a batch size of 128 and a discount factor of $\gamma = 0.9$. The data used for training are state-action-next state-reward tuples simulated with the PKPD model. To simulate the data, the VKORC1 and CYP2C9 genotypes of individuals are drawn uniformly from the possible variants. The age is drawn from a lognormal distribution with median $51$ and scale $0.15$. The treatment response is then simulated for each individual for 30 days. The dosing regimens are determined by the DQN model using daily monitoring measurements and the covariates. The model administers dosages following an $\epsilon$-greedy policy with $\epsilon = 0.05$, which balances exploration and exploitation during the training. This data-generation results in a total of 30 INR measurements and 30 dose administrations for each individual. For the training, we compute the rewards using Eq S3.1 and format the data into state-action-next state-reward tuples.

We train the model using experience replay (Lin, 1992) with a buffer size of 30,000 data points. The buffer memorises the treatment response data of 1000 simulated individuals. Each epoch, we train the model on the full buffer. At the start of the training, we fill the buffer, simulating 1000 individuals treated under the initial policy. Every 20 epochs, we randomly replace the data of 100 individuals by the data of 100 new individuals treated with the most recent policy. This enables an online learning of the DQN model, while still benefiting from experience (Lin, 1992). To stabilise the training, we use Double Q-learning (Van Hasselt et al., 2016), where two DQN models are trained in tandem. One model is used to predict the Q value of the chosen action and to find the next action with the maximal Q value during the temporal difference error evaluation (see Eq S3.8). The other network is used to evaluate the Q value for the next action. The second network is not updated trained during this iteration. After each epoch, the role of the two networks is reversed. This stabilises the training of the DQN model (Van Hasselt et al., 2016).

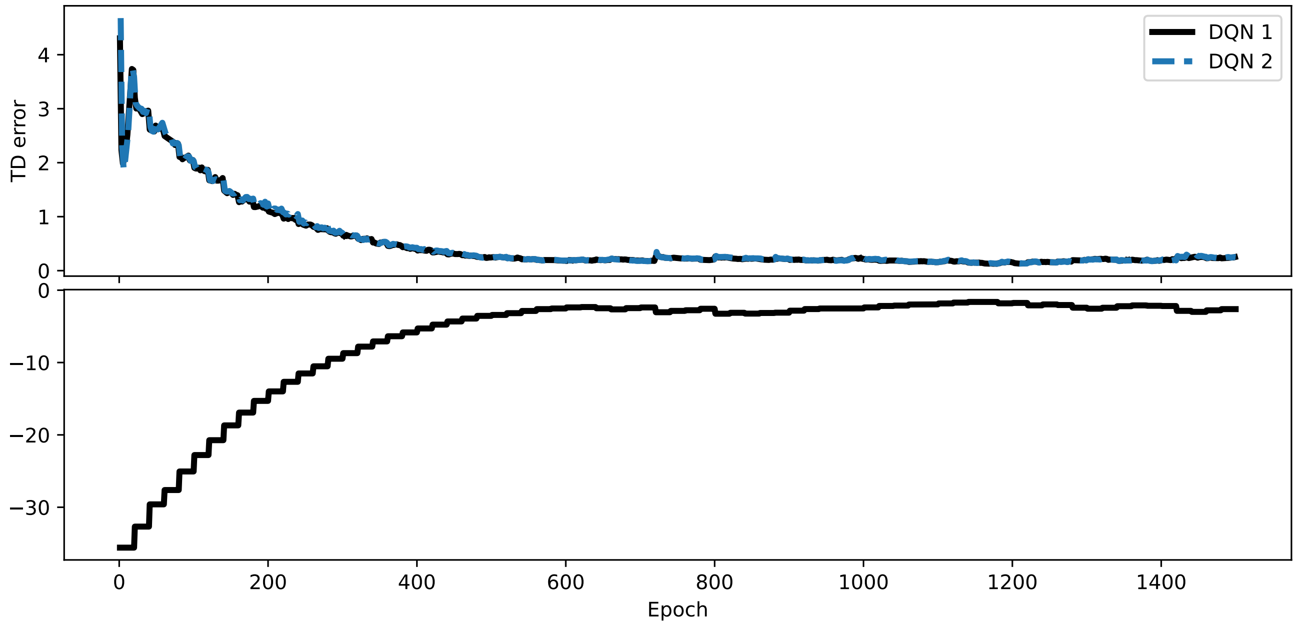


**Figure S3.1. Training of model 2:** The top panel shows the temporal difference error of the two DQN models used during Double Q-learning. The bottom panel shows the average experience buffer reward.

The results of the training are shown in Fig S3.1. The top panel of the figure shows the average temporal difference error of the two models for each epoch. The bottom panel shows the average reward in the experience buffer. The model's policy converges within the first 750 epochs of the training, reaching average rewards close to zero.

#### *Appendix S4*

### S4 MODEL 3: PHARMACOKINETIC AND PHARMACODYNAMIC MODELLING

#### S4.1 Background

This MIPD method uses pharmacokinetic and pharmacodynamic (PKPD) modelling to predict individualised dosing regimens. PKPD models are (semi-)mechanistic or empirical descriptions of treatment response dynamics, formalising biomarkers or clinical readouts as functions of time and the dosing regimen

$$\bar{y}(t, r, \psi). \tag{S4.1}$$

$\bar{y}$ denotes the quantity of interest, $t$ denotes the time, $r$ denotes the dosing regimen and $\psi$ denotes additional model parameters. This formalised relationship can be used to predict treatment responses. In Fig S4.1, we illustrate an example PKPD model of warfarin treatment developed by Hamberg et al. (2010). The model describes the time course of both the blood warfarin concentration and the INR in response to warfarin administration (Hamberg et al., 2010). Parameters of the model, $\psi$, include the half maximal inhibitory effect concentration (EC50) and the elimination rate of warfarin. We will use this model for MIPD of warfarin (see Section S4.2 for the details of the model).

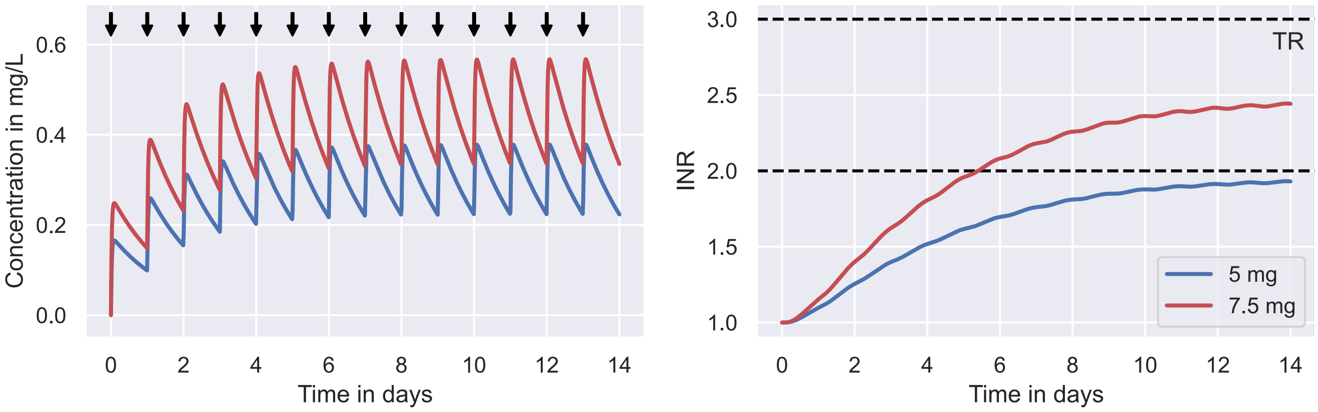


**Figure S4.1. Treatment response prediction.** The figure shows the predicted treatment response by Hamberg et al's model for two daily administered warfarin doses: 5 mg (blue curves); and 7.5 mg (red curves). The predicted warfarin concentrations over a period of 14 days are illustrated in the left panel, and the associated INR values are shown in the right panel. The dose administrations are indicated by black arrows in the left panel. The therapeutic range (TR) is indicated by dashed black lines in the right panel.

The figure shows the model predictions for a given set of model parameters and different daily doses of warfarin: 5 mg (blue); and 7.5 mg (red). The predicted warfarin concentration (see left panel) and the INR (see right panel) increase over time for both dosing regimens. The time points of the dose events are indicated by black arrows.

For a given set of model parameters, PKPD models can be used to find dosing regimens that optimise the predicted treatment response with respect to a desired treatment response. In the simplest case, this desired response is expressed in terms of a target value for the quantity of interest

$$\bar{y}(t, r^*, \psi) = y^*. \tag{S4.2}$$

$y^*$ denotes the target value and $r^*$ denotes the *optimal* dosing regimen. However, weighted ranges or distributions of target values can also be used to define optimal dosing regimens (Maier et al., 2021).

In practice, constraints on the dose amount, the type and the frequency of dose administrations, as well as delays in the treatment response make it difficult (or impossible) to realise the desired treatment response exactly. It is therefore customary to relax the optimisation target to treatment responses that only approximately equal the desired treatment response. A simple approach to quantify the proximity of a predicted treatment response and the desired treatment response is the mean squared error (MSE)

$$\text{MSE} = \frac{1}{n}\sum_{j=1}^{n}\Big(y^* - \bar{y}(t_j, r, \psi)\Big)^2, \tag{S4.3}$$

where $n$ denotes the number of time points, $t_j$, at which the model predictions are compared to the target. Other distance metrics can also be used to define optimal dosing regimens. For weighted ranges or distributions of target values, for example, optimal dosing regimens can be found by optimising the score of predicted treatment responses with respect to the distribution of target values (Maier et al., 2021).

To illustrate the concept of optimal dosing regimens, we show in Fig S4.2 the dosing regimen which minimises the MSE of the predicted treatment response with respect to a target treatment response of $y^* = 2.5$ for Hamberg et al's model. The details for the estimation of $r^*$ are reported in Section S4.3. The dosing regimen is constrained to dose amounts between $0$ mg and $30$ mg that can be administered as combinations of commercially available warfarin tablets, allowing dosages of $0$ mg, $1$ mg and any dose $\geq 2$ mg in $0.5$ mg steps.

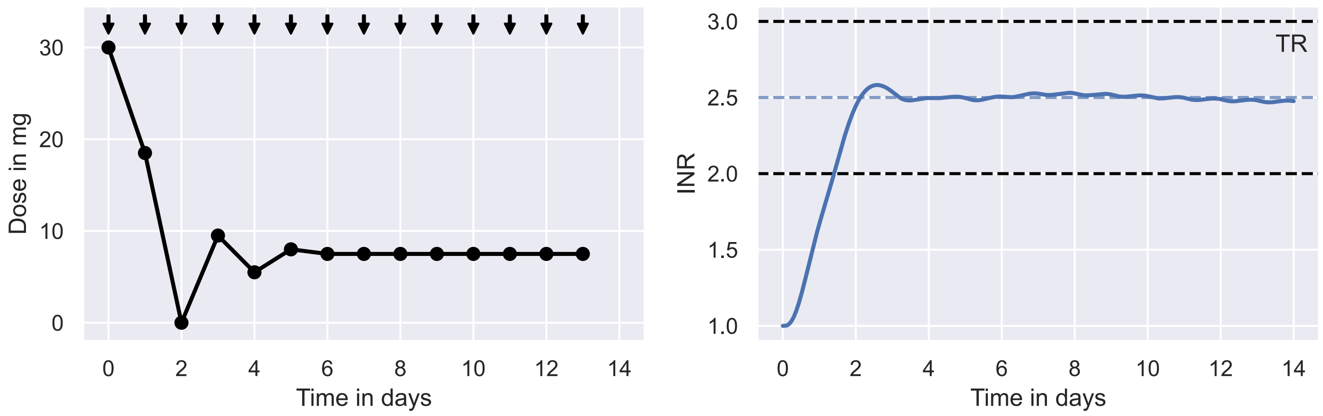


**Figure S4.2. Dosing regimen optimisation.** The figure shows the dosing regimen that minimises the squared distance between predicted INR values and a target value of 2.5 (left panel) and the associated INR response (right panel). The dose administrations are indicated by black arrows in the left panel. In the right panel the therapeutic range (TR) (black) and the target value (blue) are indicated by dashed lines.

The figure shows that a nontrivial series of induction doses ranging from $0$ mg to $30$ mg (see days 0 to 6 in the left panel) is able to reach the lower threshold of the therapeutic window within approximately 1 day, resulting in a predicted time in the therapeutic range of roughly $93\,\%$ within the first 14 treatment days. The maintenance dose from day 6 onwards is $7.5$ mg.

##### S4.1.1 Inter-individual variability

Analogously to the virtual clinical trial model introduced in General framework for MIPD trial simulation, PKPD models can be extended to describe variability in the treatment response using a population model

$$p(\psi|\theta, \chi), \tag{S4.4}$$

where each $\psi$ in the distribution represents an individual. $\theta$ denotes the parameters of the population distribution and $\chi$ denotes the covariates of the IIV. The treatment response for an individual characterised by the parameters, $\psi$, is predicted with the PKPD model (see Eq S4.1). The PKPD model together with the population model is referred to as nonlinear mixed effects (NLME) model.

To demonstrate the effect of varying parameter values on the treatment response predictions, we illustrate Hamberg et al's NLME model in Fig S4.3 (Hamberg et al., 2010). For simplicity, we only consider variability in the EC50 across individuals. The remaining parameter values are fixed. The model incorporates single nucleotide polymorphisms of the VKORC1 genotype as covariates of the EC50 variability (see Section S4.2 for the details of the model).

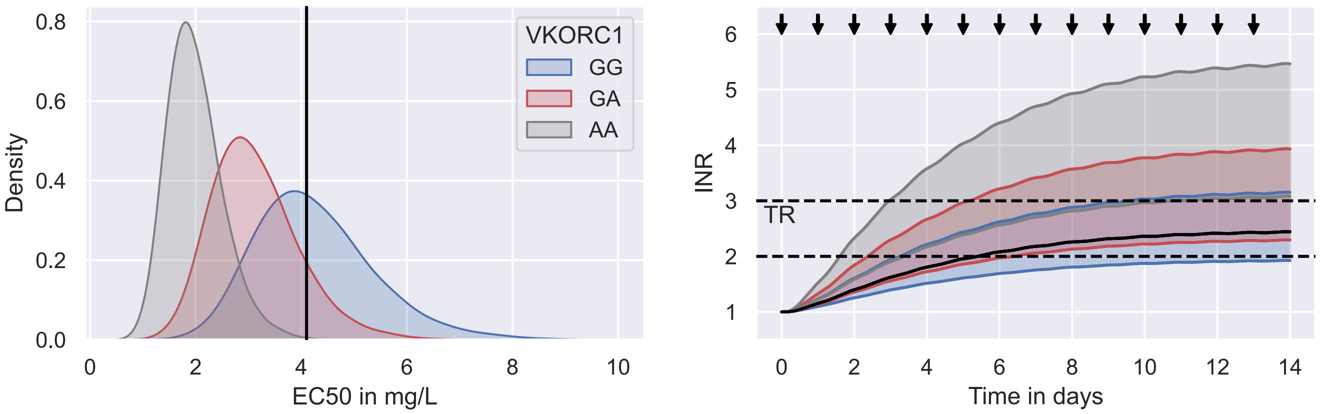


**Figure S4.3. Inter-individual variability of warfarin EC50 and treatment response.** The figure shows Hamberg et al's model of the warfarin EC50 variability across patients (see left panel) and the predicted INR response variability when daily doses of 7.5 mg warfarin are administered to all individuals. The colours indicate individuals with the same VKORC1 genotypes: GG (blue), GA (red) and AA (grey). The coloured areas in the right panel indicate the 5th to 95th percentile interval of the INR response across individuals with the same VKORC1 genotype. The black solid line represents a typical patient with the GG genotype. The dose administrations are indicated by black arrows in the right panel. The therapeutic range (TR) is indicated by dashed black lines.

The figure shows distributions of EC50 values across individuals with the same VKORC1 genotype (see left panel) and their treatment response to daily warfarin doses of 7.5 mg (see right panel) predicted by Hamberg et al's NLME model. The VKORC1 genotype partially explains the IIV of the EC50. Individuals with the GG genotype (blue) have on average the largest EC50, followed by individuals with the GA genotype (red) and the AA genotype (grey). These differences in EC50 values result in a large treatment response variability across individuals (see right panel). The majority of individuals with the GG genotype reach INR values within the therapeutic range, while most individuals with the AA genotype assume INR values larger than three after 14 days of treatment. Notably, Hamberg et al's model predicts substantial IIV that is not explained by the VKORC1 genotype, such that individuals with the GG genotype have EC50 values ranging from 1 mg/L to 9 mg/L. A typical individual with the GG genotype is illustrated by the black solid lines.

Large treatment response variability implies that also optimal dosing regimens will vary between individuals (see Fig S4.4). The top left panel shows the treatment response across individuals treated with optimised dosing regimens. For each VKORC1 genotype, 1000 individuals are simulated. The dose distributions of the first 6 doses are visualised in the right panel. Each subpanel shows the dose distribution across individuals with the different VKORC1 genotypes: GG (blue); GA (red); and AA (grey). The distribution of maintenance doses is illustrated in the bottom left panel.

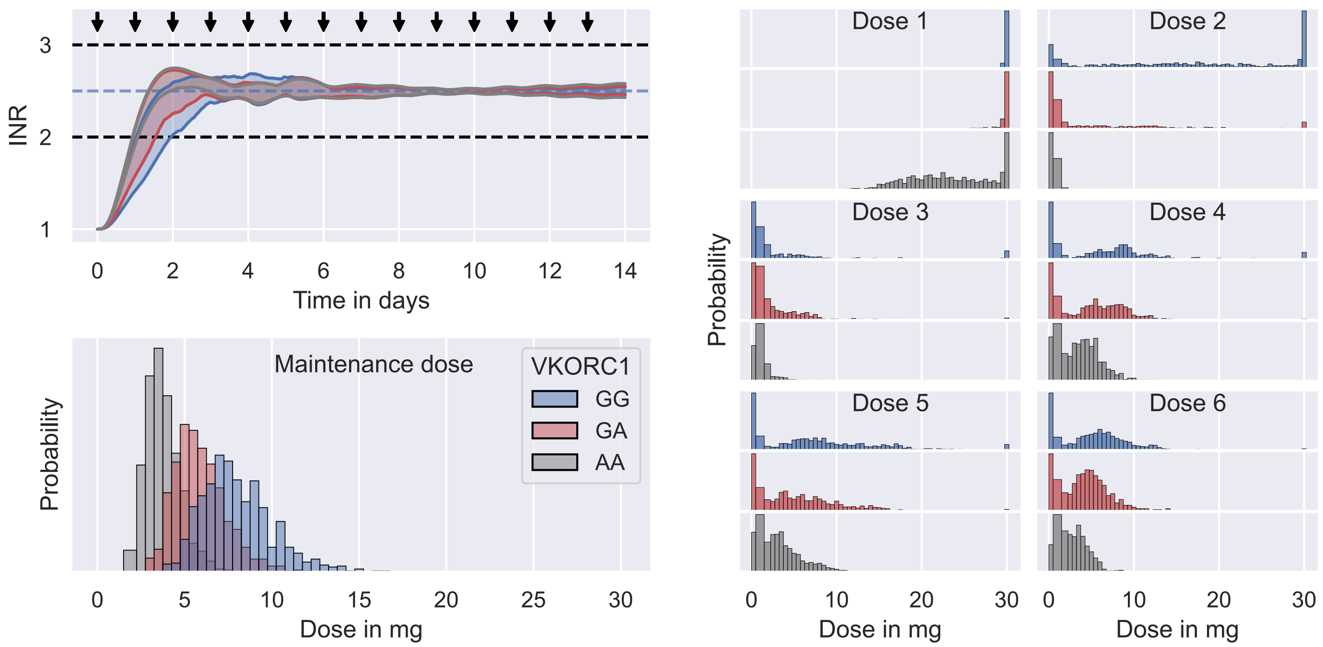


**Figure S4.4. Variability of optimal dosing regimen.** The figure shows predicted treatment responses (top left panel) of 3000 individuals to dosing regimens that minimise the MSE to a target INR of 2.5. The treatment response distributions are illustrated by the 5th to 95th INR percentile intervals across individuals with the same VKORC1 genotype: GG (blue); GA (red); and AA (grey). Each genotype group is represented by 1000 individuals. The distribution of optimal maintenance dosages is illustrated in the bottom left panel. The distributions of optimal induction dosages on days 0 to 5 are illustrated in the 6 panels on the right. The dose administrations are indicated by black arrows in the top left panel. The therapeutic range (black) and the target value (blue) are indicated by dashed lines.

The figure shows that optimising dosing regimens for individuals reduces the predicted treatment response variability (see top left panel), achieving TTRs of $\geq 86\,\%$ within the first 14 treatment days for most individuals. This improved homogeneity in the treatment response is achieved by varying dosing regimens across individuals. Maintenance dosages for individuals with the AA VKORC1 genotype range from 2 mg to 7.5 mg, while dosages for individuals with the GG genotype span a range of $\geq 10$ mg (see bottom left figure). This variability is even more pronounced during the induction phase of the treatment (see right 6 panels).

##### S4.1.2 Covariate-based dosing regimen optimisation

Fig S4.4 shows that dosing regimen optimisation can account for variability in the predicted treatment response when the EC50s of treated individuals are known. In practice, the EC50, and more generally the model parameters, $\psi$, that best predict an individual's treatment response are, however, unknown. One approach to alleviate this problem is to use calibrated NLME models to suggest model parameters for individuals based on covariates of the IIV (Hamberg et al., 2010; Hamberg and Wadelius, 2014). In particular, we can predict a set of typical parameter values for an individual with the covariates $\chi$ using the distribution of model parameters, $p(\psi|\theta, \chi)$. In this article, we define typical parameter values as the median of the parameter distribution. Other choices such as the mean or the values with maximum probability density (MAP) are also possible.

In Fig S4.5, we illustrate this covariate-based dosing regimen optimisation using Hamberg et al's NLME model. The left panel shows the typical EC50 values for individuals with the GG (blue), GA (red) and AA (grey) VKORC1 genotypes predicted by Hamberg et al's NLME model (see solid lines). The dashed lines illustrate the predicted EC50 distributions by the model. The middle panel shows the optimal dosing regimens associated with the typical parameters, obtained by minimising the MSE of the predicted treatment response to the target INR of $2.5$. These optimal dosing regimens are able to partially account for the treatment response variability across individuals (see right panel), such that the majority of the predicted treatment responses is inside the TR.

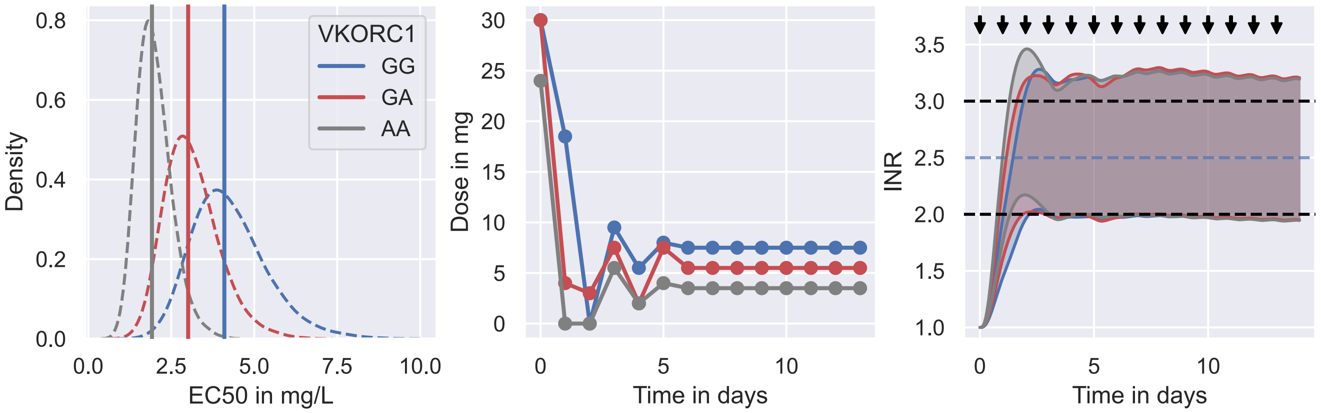


**Figure S4.5. Covariate-based dosing regimen optimisation.** The left panel shows typical EC50 values for individuals with the GG (blue), GA (red) and AA (grey) VKORC1 genotypes predicted by Hamberg et al's NLME model as solid lines. The distributions of EC50 values across individuals are indicated by dashed lines. The middle panel illustrates the dosing regimens that minimise the MSE of the predicted treatment response for the typical EC50 values to a target INR of $2.5$. The right panel shows the treatment responses of 3000 individuals (1000 for each genotype). Each individual is treated with the optimal dosing regimen predicted for the typical EC50 (see middle panel). The treatment response distributions are illustrated by the 5th to 95th INR percentile intervals across individuals with the same VKORC1 genotype. The dose administrations are indicated by black arrows in the top left panel. The therapeutic range (black) and the target value (blue) are indicated by dashed lines.

##### S4.1.3 Bayesian dosing regimen optimisation

Covariate-based dosing regimen optimisation is able to partially account for treatment response variability by predicting typical model parameters based on covariates of the IIV. These predictions rely on the calibration of the NLME model, with no mechanism to adjust predictions if the predicted treatment response deviates from the observed treatment response due to residual uncertainty in the model parameters, model misspecification or IOV of the individual's treatment response.

A complementary approach that addresses these shortcomings is Bayesian dosing regimen optimisation (Maier et al., 2020). Bayesian dosing regimen optimisation uses feedback of the treatment response in form of measurements from therapeutic drug/biomarker monitoring (TDM) to iteratively calibrate the model to the treatment response of an individual, making dosing regimen adjustments and more personalised predictions possible. To this end, Bayes' rule is used to update the distribution of parameter values predicted by the NLME model, $p(\psi|\theta,\chi)$, to a distribution that is also consistent with a patient's TDM data

$$p(\psi|\theta,\chi) \mapsto p(\psi|\theta,\chi,\mathcal{D}) \tag{S4.5}$$

We use $\mathcal{D}$ to denote the TDM data, containing measurements of the treatment response. In the Bayesian literature, the distribution before the update, $p(\psi|\theta,\chi)$, is referred to as the prior distribution and the refined distribution, $p(\psi|\theta,\chi,\mathcal{D})$, is referred to as the posterior distribution. For details on the calibration, see Section S4.3. The posterior distribution predicts parameter values that are consistent with both the prior distribution as well as the observed treatment response of the individual. This makes improved predictions of model parameters for the individual possible. In Fig S4.6, we illustrate this concept of Bayesian dosing regimen optimisation using Hamberg et al's NLME model as an example.

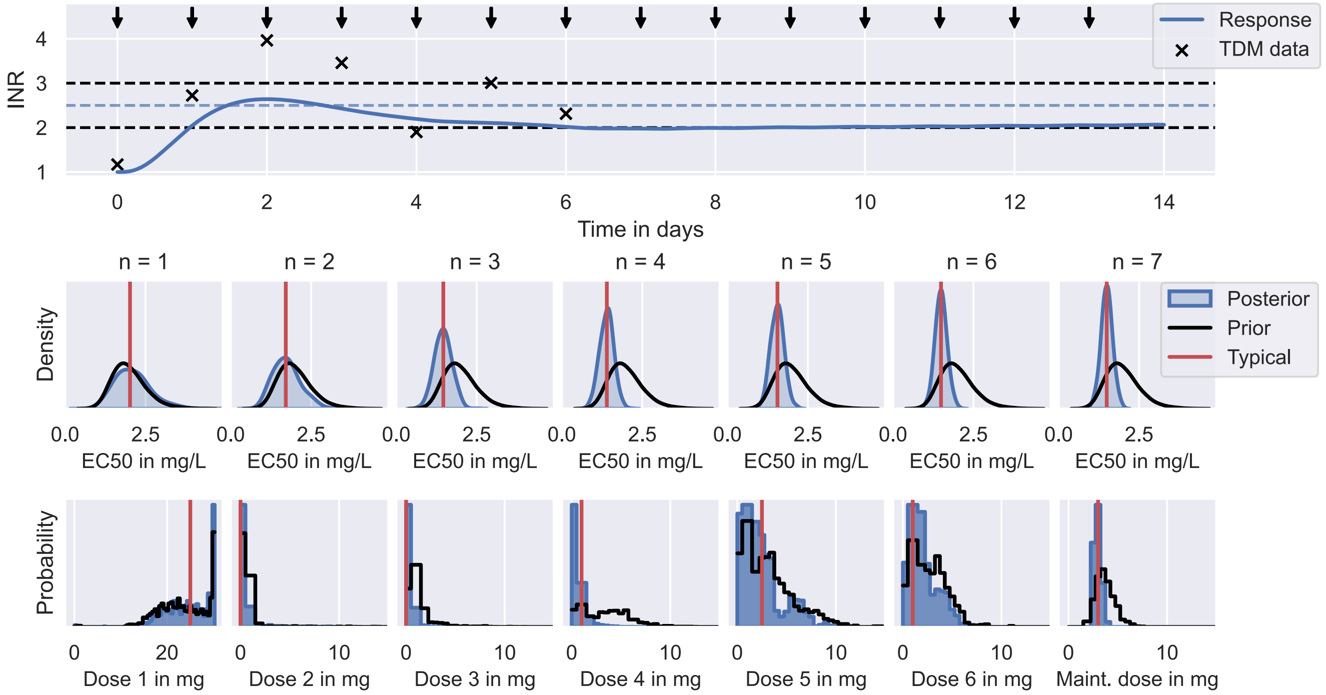


**Figure S4.6. Bayesian dosing regimen optimisation.** The figure illustrates the dosing regimen individualisation for an individual with the AA VKORC1 genotype using Bayesian dosing regimen optimisation. The predicted treatment response (blue line) and the TDM measurements of the response (black crosses) are illustrated in the top panel. The dose administrations are indicated by black arrows. The therapeutic range (black) and the target value (blue) are indicated by dashed lines. The posterior distributions (blue) inferred from the TDM measurements on day 0 ($N = 1$) to day 6 ($N = 7$) are illustrated in the middle panel. The typical parameter value of the posterior distribution is illustrated in red and the prior distribution predicted by the NLME model is depicted in black. The bottom panel shows the optimal dose distributions associated with the posteriors (blue), the typical parameters and the prior (black).

The figure shows the predicted treatment response (blue) of an individual with the AA VKORC1 genotype treated with an iteratively optimised dosing regimen (see top panel). The TDM measurements of the INR used to inform the dosing strategy are collected before each dose administration on days 0 to 6 and are illustrated by black crosses. From day 6 onwards the dose amount remains unchanged. The posterior distributions of EC50 values from days 0 to 6 (blue) are illustrated in the middle panel, together with the typical EC50s of the posteriors (red) and the prior distribution predicted by the NLME model (black). The associated predictions of the optimal dosing regimen are illustrated in the bottom panel. The dosing regimen associated with the typical parameter values (red) is used to predict the treatment response in the top panel.

#### S4.2 Implementation

We use Hamberg et al's PKPD model to predict warfarin treatment response dynamics (Hamberg et al., 2010). The model is schematically illustrated in Fig S4.7.

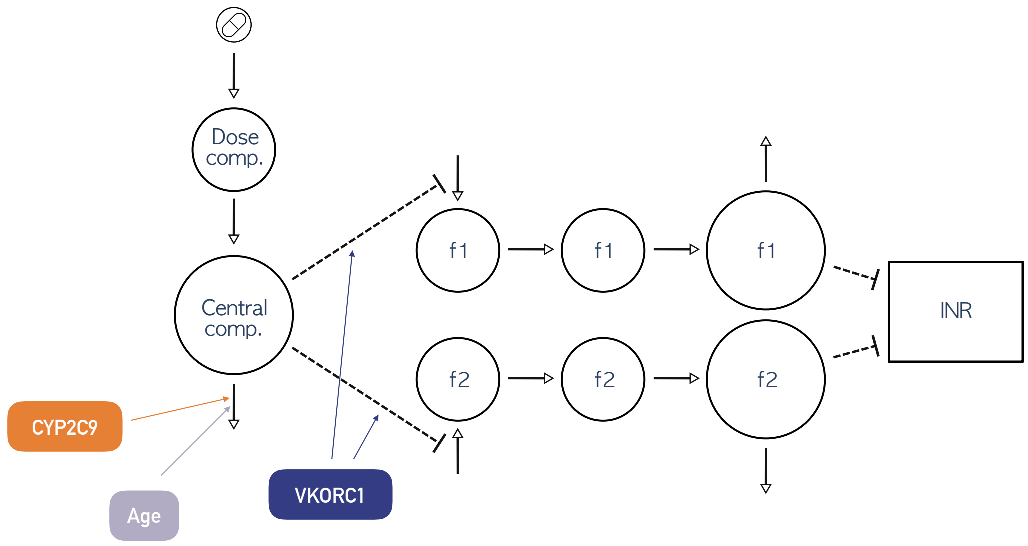


**Figure S4.7. Hamberg et al's model of warfarin treatment response.** The figure shows a diagrammatic representation of Hamberg et al's model of the INR response to warfarin treatment. The diagram illustrates the PK model component (left side) and the PD model component (right side). The drug (see pill in the top left corner) is administered to a dose compartment from which it moves into a central compartment. From the central compartment warfarin is cleared with a rate that is influenced by the CYP2C9 genotype and the age of an individual. The drug concentration in the central compartment inhibits the production of the (effective) coagulation factors $f_1$ and $f_2$. The inhibitory effect of the drug is influenced by the VKORC1 genotype. Each coagulation factor is produced in one compartment, moved to an intermediary compartment and is cleared in a third compartment. The coagulation factor concentrations in the final compartments determine the INR. The lower the coagulation factor concentrations the larger the INR.

The model provides a semi-mechanistic description of the INR response to warfarin treatment

$$\bar{y} = \bar{y}_0 + \bar{y}_{max} \left( 1 - \frac{f_1 + f_2}{2} \right), \tag{S4.6}$$

where $\bar{y}$ denotes the INR (see right-most compartment in Fig S4.7). $\bar{y}_0$ and $\bar{y}_{max}$ are model parameters, denoting the baseline INR and the maximum INR increase from warfarin treatment, respectively. $f_1$ and $f_2$

denote concentrations of effective coagulation factors relative to their baseline concentration prior to the treatment. The reduction of the coagulation factors as a result of warfarin exposure is modelled by a system of ordinary differential equations

$$\begin{aligned} \frac{\mathrm{d}f_i}{\mathrm{d}t} &= k_i f_{i,2} - k_i f_i, \qquad i \in \{1, 2\} \\ \frac{\mathrm{d}f_{i,2}}{\mathrm{d}t} &= k_i f_{i,1} - k_i f_{i,2} \\ \frac{\mathrm{d}f_{i,1}}{\mathrm{d}t} &= k_i \left(1 - \frac{\kappa c^\gamma}{c^\gamma + c_{50}^\gamma}\right) - k_i f_{i,1}, \end{aligned} \tag{S4.7}$$

where $t$ denotes the time. $f_{i,1}$ and $f_{i,2}$ denote the relative concentrations of coagulation factor $i$ in two compartments, implementing a delay of the INR response upon warfarin exposure (see $f_1$ and $f_2$ compartments in the middle of Fig S4.7). The delay is parametrised by the transition rates, $k_1$ and $k_2$, between the compartments. $\kappa$ denotes the maximal inhibitory effect of warfarin on the coagulation factor synthesis, $c$ denotes the blood plasma concentration of warfarin and $c_{50}$ denotes its half maximal inhibitory effect concentration. $\gamma$ parametrises the potency of warfarin.

The pharmacokinetics of warfarin are described by a 1-compartment PK model with a dose compartment

$$\begin{aligned} \frac{\mathrm{d}a_d}{\mathrm{d}t} &= -k_a a_d + r(t) \\ \frac{\mathrm{d}a_c}{\mathrm{d}t} &= k_a a_d - k_e a_c, \end{aligned} \tag{S4.8}$$

where $a_d$ and $a_c$ denote the drug amount in the dose and central compartments, respectively (see compartments on the left of Fig S4.7). $k_a$ denotes the absorption rate and $k_e$ the elimination rate. $r$ denotes the time-dependent dose rate of warfarin, implementing the dosing regimen (see Section S1.1.1). The warfarin concentration is given by the drug amount in the central compartment divided by its effective volume of distribution, $c = a_c/v$. As a result, the parameters of the model are $\psi = (\bar{y}_0, \bar{y}_{max}, k_a, k_e, \kappa, \gamma, c_{50}, k_1, k_2, v)$. The initial values of the state parameters are fixed to $f_i(t = 0) = f_{i,1}(t = 0) = f_{i,2}(t = 0) = 1$ and $a_d(t = 0) = a_c(t = 0) = 0$.

The population model captures variability in the treatment response using varying elimination rates, half maximal inhibitory effect concentrations, and effective volumes of distribution across patients. The values of $k_e$, $c_{50}$ and $v$ are assumed to randomly deviate from a set of typical parameter values

$$k_e = \mu_{k_e}\, \mathrm{e}^{\eta_{k_e}}, \quad c_{50} = \mu_{c_{50}}\, \mathrm{e}^{\eta_{c_{50}}}, \quad v = \mu_v\, \mathrm{e}^{\eta_v}, \tag{S4.9}$$

where $(\mu_{k_e}, \mu_{c_{50}}, \mu_v)$ denote the typical values and $(\eta_{k_e}, \eta_{c_{50}}, \eta_v)$ denote the individual-specific random deviations. For $\eta < 0$, the parameter of an individual is smaller than the typical value and for $\eta > 0$ it exceeds the typical value. The model assumes that the $\eta$ follow Gaussian distributions centered at zero with variance $\sigma^2$, e.g. $\mathcal{N}(\eta_{k_e}|0, \sigma^2_{k_e})$. The remaining parameters are fixed across individuals. The model further assumes that the variability of $k_e$ and $c_{50}$ can be partially explained by 3 covariates: 1. the CYP2C9 genotype (one of *1/*1, *1/*2, *1/*3, *2/*2, *2/*3 and *3/*3); 2. the VKORC1 genotype (one of GG, GA and AA); and 3. the age. In particular, the typical elimination rate and the typical half maximal inhibitory effect concentration are modelled as functions of the covariates $\chi = (c_1, c_2, v_1, v_2, t_\star)$

$$\begin{aligned} \mu_{k_e}(c_1, c_2, t_\star) &= (k_{c_1} + k_{c_2})\,(1 - \tanh(r_{age}(t_\star - 71))) \\ \mu_{c50}(v_1, v_2) &= c_{v_1} + c_{v_2}, \end{aligned} \tag{S4.10}$$

where $c_1, c_2 \in \{*1, *2, *3\}$ denote the genotypes of the CYP2C9 alleles, $v_1, v_2 \in \{\mathrm{G}, \mathrm{A}\}$ denote the genotypes of the VKORC1 alleles and $t_\star$ denotes the age in years. The tanh-function is a modification of the original model that avoids non-positive elimination rates, $\mu_{k_e} \leq 0$, for $t_\star \gg 71$. Eqs S4.9 and S4.10 define the population distribution of the model parameters across individuals conditional on the covariates

$$\begin{aligned} p(\psi|\theta, \chi) = \; & \mathrm{LN}\,(k_e \,|\, \mu_{k_e}(c_1, c_2, t_\star), \sigma_{k_e})\; \mathrm{LN}\,(c_{50} \,|\, \mu_{c50}(v_1, v_2), \sigma_{c50})\; \mathrm{LN}\,(v \,|\, \mu_v, \sigma_v) \\ & \delta(\mu_{\bar{y}_0} - \bar{y}_0)\, \delta(\mu_{\bar{y}_{max}} - \bar{y}_{max})\, \delta(\mu_{k_a} - k_a)\, \delta(\mu_\kappa - \kappa)\, \delta(\mu_\gamma - \gamma) \\ & \delta(\mu_{k_1} - k_1)\, \delta(\mu_{k_2} - k_2), \end{aligned} \tag{S4.11}$$

where $\theta = (k_{*1}, k_{*2}, k_{*3}, r_{age}, \sigma_{k_e}, c_G, c_A, \sigma_{c50}, \mu_v, \sigma_v, \mu_{\bar{y}_0}, \mu_{\bar{y}_{max}}, \mu_{k_a}, \mu_\kappa, \mu_\gamma, \mu_{k_1}, \mu_{k_2})$ denotes the parameters of the population model, $\mathrm{LN}(x|\mu, \sigma) = \mathrm{e}^{(\log\mu - \log x)^2/2\sigma^2}/x\sigma\sqrt{2\pi}$ denotes a lognormal distribution and $\delta(\mu - x)$ denotes a Dirac delta distribution.

In their original study, Hamberg et al. (2010) fixed the parameter values of the absorption rate, the baseline INR and the maximum INR increase to $\mu_{k_a} = 2$ 1/h, $\mu_{\bar{y}_0} = 1$ and $\mu_{\bar{y}_{max}} = 20$, respectively. In our study, we fix the absorption rate and the maximum INR shift to the same values. However, we do not fix the baseline INR. Instead, we replace the pooled baseline INR population model in Eq S4.12 with a lognormal covariate population model

$$\mathrm{LN}\,(\bar{y}_0 \,|\, \mu_{\bar{y}_0}(v_1, v_2), \sigma_{\bar{y}_0}) \quad \text{with} \quad \mu_{\bar{y}_0}(v_1, v_2) = \bar{y}_{0,v_1} + \bar{y}_{0,v_2}. \tag{S4.12}$$

We do this in order to account for IIV in the baseline INR (see Fig S1.1).

#### S4.3 Dose prediction

Individualised dosing regimens are predicted based on the covariates and the monitoring data using Bayesian dosing regimen optimisation (see Fig S4.6). The model is first calibrated to the individual's data. In a second step, the calibrated model is used to predict an optimised dosing regimen.

To calibrate the model, we use Bayesian inference to derive a distribution of model parameters compatible with both, the population model, $p(\psi|\chi, \theta)$, and the individual's data, $\mathcal{D}_i = ((y_{i1}, t_{i1}), \ldots, (y_{in}, t_{in}), r_i, \chi_i)$. $i$ labels the data specific to an individual and $n$ denotes the number of monitoring measurements. This distribution is referred to as posterior distribution in the Bayesian inference literature and is derived from $p(\psi|\chi, \theta)$ and $\mathcal{D}_i$ using Bayes' rule

$$\log p(\psi|\mathcal{D}_i, \theta) = \sum_{j=1}^{n} \log p(y_{ij}|t_{ij}, r_i, \psi) + \log p(\psi|\chi_i, \theta) + \text{constant}. \tag{S4.13}$$

We use the population model, $p(\psi|\chi_i, \theta)$, to capture the prior information about the model parameters. $p(\psi|\chi, \theta)$ is estimated from trial data during the calibration of the model (see Section S4.4).

The first term in Eq S4.13 is the likelihood of the model parameters, which is defined using a measurement distribution, $p(y|t, r, \psi)$. The measurement distribution is an extension of the PKPD model output (see

Eq S4.1), accounting for discrepancies between measurements, $y$, and the model output, $\bar{y}$. A common choice to model these deviations is to use normally distributed residual errors, $y = \bar{y} + \epsilon$, where $\epsilon$ is normally distributed, $\mathcal{N}(\epsilon|0, \sigma^2)$. However, for $\mathcal{O}(\sigma) = \bar{y}$, this error model can lead to unbiological (negative) measurements. We therefore model the errors on the log-scale, $\log y = \log \bar{y} + \epsilon$, which constrains measurements to positive values. This error model defines a lognormal measurement distribution, $p(y|t, r, \psi) = \mathrm{LN}(y|\bar{y}(t, r, \psi), \sigma)$.

We implement the PKPD model, the error model and the posterior distribution in `chi` (Augustin, 2021) and use the adaptive covariance matrix Markov Chain Monte Carlo sampling algorithm implemented in `pints` (Clerx et al., 2019) to sample from the posterior distribution. We run the algorithm for 10.000 iterations and discard the first 5000 iterations as warmup. We estimate the convergence of the marginal posterior distributions using the $\hat{R}$-statistic. All posteriors during the MIPD trial reach $\hat{R} \leq 1.01$. The results of the inference, as well as data and code to reproduce the results are hosted on GitHub (https://github.com/DavAug/mipd-warfarin).

To predict individualised dosing regimens, we minimise the mean squared error of the PKPD model prediction to the target INR value, $y^* = 2.5$, using $n$ evaluation time points

$$\mathrm{MSE}(r_i) = \frac{1}{n} \sum_{j=1}^{n} \Big( \bar{y}(t_j, r_i, \psi_i) - \bar{y}^* \Big)^2 . \tag{S4.14}$$

The time points of the evaluation are a modelling choice. In this article, we choose $n = 5000$ equidistant time points starting at $t_1 = 0\,\mathrm{h}$ and ending at $t_{5000} = 50 \times 24\,\mathrm{h}$ since the start of the treatment, ensuring that the treatment response remains close to the target value even after the 19-day MIPD trial. The parameters, $\psi_i$, used to predict the treatment response are specific to each individual and are estimated from the posterior distribution in Eq S4.13. In this article, we approximate $\psi_i$ by the typical value of the posterior distribution, i.e. $\psi_i = \mathrm{median}[\{\psi_{is}\}]$, where $\psi_{is}$ denotes a sample generated by the MCMC sampling algorithm.

The optimisation parameters are the parameters of the dosing regimen, including dosages, administration times and dose durations (see Eq S1.1). In this article, we constrain dosing regimens to daily bolus dosages, which we implement by setting the administration times to $t_k = (k-1) \times 24\,\mathrm{h}$ and the dose durations to $\Delta_k = 0.01\,\mathrm{h}$. To ensure that the optimisation converges to a single maintenance dose (as opposed to a maintenance dose pattern that may require different doses on different days), doses following the 19th treatment day are set to the 19th dose, i.e. $d_k = d_{19}$ for $k \geq 19$. This leaves 19 parameters for the optimisation – the first 19 warfarin dosages.

We optimise these 19 dosages using a sliding window approach: 1. Doses already administered are fixed to the administered values; 2. the next 7 doses are optimised; 3. the remaining doses are set to the last dose in the optimisation window. At the start of the treatment, no doses have been administered and the first 7 doses are optimised. Doses 8 to 19 are set to the dose amount of dose 7. On the next day, the first dose is fixed, and doses 2 to 8 are optimised. The remaining doses are set to the dose amount of dose 8. This procedure continues until dose 14, where the condition of setting all doses following dose 19 reduces the window size to 6 days, such that on the last day of the trial only the 19th dose is optimised.

The mean squared error objective function is implemented using `chi` (Augustin, 2021) and `pints` (Clerx et al., 2019). We minimise the objective function using the covariance matrix adaptation-evolution strategy (CMA-ES) optimisation algorithm implemented in `pints`.

#### S4.4 Fitting

We calibrate the model to clinical phase I, II and III data simulated in Simulating trial phases prior to MIPD. The data includes monitoring data, covariates and warfarin dosing regimens of a total of 1160 simulated patients. We calibrate the model using hierarchical Bayesian inference.

##### S4.4.1 Calibration to trial phase I data

We first calibrate the model parameters to the data from trial phase I. The data contains warfarin concentration measurements of 60 individuals (see left panel in Fig 5). Denoting the data as $\mathcal{D}_1 = (\mathcal{D}_{1\,1}, \ldots, \mathcal{D}_{1\,60})$, where $\mathcal{D}_{1i}$ denotes the data of individual $i$ analogously to Section S4.3, we can define a hierarchical posterior for the population model parameters using Bayes rule

$$\log p(\theta, \Psi_1 | \mathcal{D}_1) = \sum_i \log p(\mathcal{D}_{1i} | \psi_{1i}) + \sum_i \log p(\psi_{1i} | \theta, \chi_{1i}) + \log p(\theta) + \text{constant}. \tag{S4.15}$$

$\Psi_1 = (\psi_{1\,1}, \ldots, \psi_{1\,60})$ denotes the individual-level parameters of the individuals in the trial. The focus of the calibration is the posterior distribution of the population-level parameters, $\theta$, obtained by marginalising over the individual-level parameters, $p(\theta | \mathcal{D}_1) = \int \mathrm{d}\Psi_1 \, p(\theta, \Psi_1 | \mathcal{D}_1)$. For an in-depth review of Bayesian inference of NLME models, we refer to the methods section in Augustin et al. (2023).

The first term in Eq S4.15 quantifies the likelihood of individual-level parameters using a measurement distribution analogously to Section S4.3. We model the error of the simulated warfarin concentrations with a lognormal error model. The scale parameter of the error model, $\sigma$, is pooled across individuals during the inference.

The data from trial I does not contain INR measurements and can therefore only be used to calibrate the PK component of the model. This leaves 8 parameters for the calibration: $(k_{*1}, k_{*2}, k_{*3}, r_{age}, \sigma_{k_e}, \mu_v, \sigma_v, \mu_\sigma)$, where we use $\mu_\sigma$ to denote the pooled scale parameter of the measurement distribution.

At the start of the calibration, we assume minimal prior knowledge about the model parameters to mimic a typical PKPD modelling scenario. We define weakly informative marginal priors whose responsibility is only to constrain parameter values to biologically meaningful ranges

$$\begin{aligned} \log k_{*1} &\sim \mathcal{N}(-3, 0.1) \\ \sigma_{k_e} &\sim \mathrm{LN}(-1, 0.3) \\ \frac{k_{*1} - k_{*2}}{k_{*2}} &\sim \mathcal{U}(0, 1) \\ \frac{k_{*1} - k_{*3}}{k_{*2}} &\sim \mathcal{U}(0, 1) \\ r_{age} &\sim \mathcal{N}(0, 0.01) \\ \log \mu_v &\sim \mathcal{N}(2.7, 0.1) \\ \sigma_v &\sim \mathrm{LN}(-1, 0.3) \\ \mu_\sigma &\sim \mathrm{LN}(-1, 0.3). \end{aligned} \tag{S4.16}$$

We construct the prior distribution, $p(\theta)$, by multiplying the marginal distributions.

We implement the PKPD model, the likelihood, the population model, the prior distribution and the posterior distribution in `chi` (Augustin, 2021). Samples from the posterior distribution are generated using the No-U-Turn sampler (NUTS) implemented in `pints` (Clerx et al., 2019). We run the algorithm for 1500 iterations and discard the first 500 iterations as warmup. We estimate the convergence of the marginal posterior distributions using the $\hat{R}$-statistic. All marginal posteriors have $\hat{R} \leq 1.01$. We show the inferred posterior distribution in Fig S4.8. The results of the inference, as well as data and code to reproduce the results are hosted on GitHub (https://github.com/DavAug/mipd-warfarin).

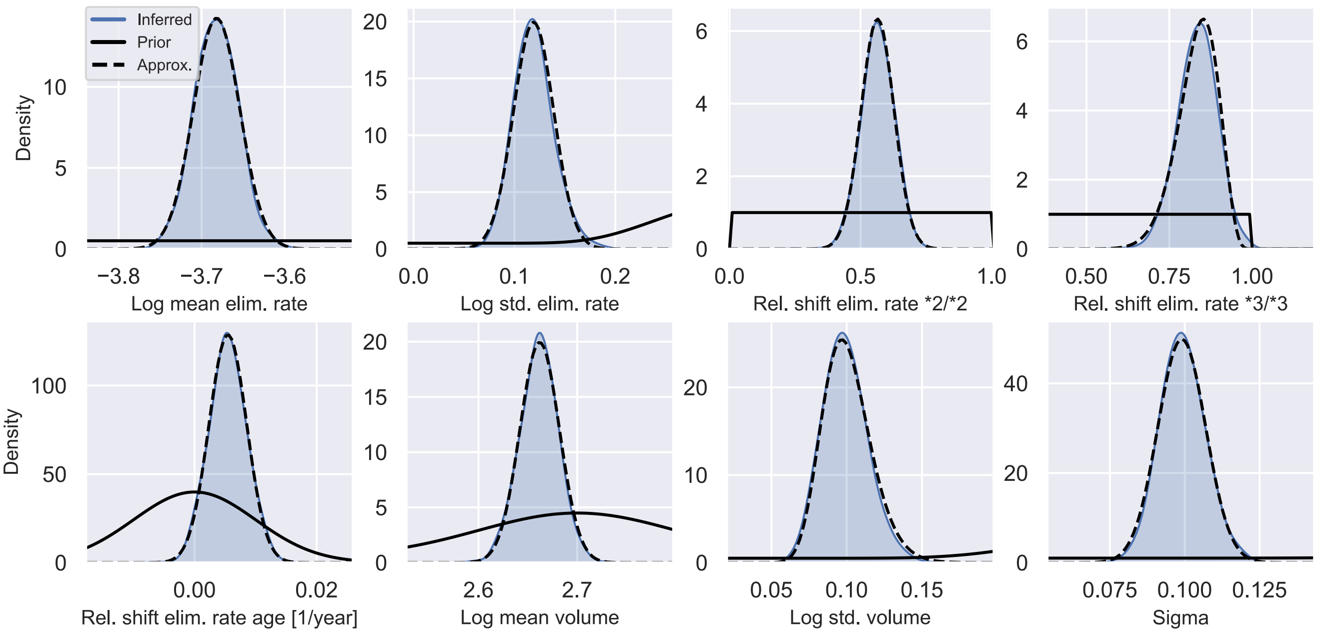


**Figure S4.8. Marginal posteriors after calibration to trial phase I data.** The figure shows the marginal posteriors of the model parameters after calibrating Hamberg et al's model to the phase I trial data. The priors are illustrated in black (solid lines) and the inferred posteriors are illustrated in blue using kernel density estimates. The approximations of the posteriors are illustrated by black dashed lines.

Fig S4.8 shows the results of the calibration. The marginal prior distributions are illustrated in black (solid lines) and the posterior distributions are illustrated in blue. Approximations of the posterior distributions, explained and used in Section S4.4.2, are illustrated in black (dashed lines).

The goodness-of-fit is visualised in the top panel of Fig S4.9. The left panel shows the concentration measurements used for the calibration. Measurements taken from the same individual are connected using a solid line. The nominal administration time of the warfarin dose is indicated using a blue arrow. The middle panel shows the quality of the individual fits. Each scatter point represents the fit of the PKPD model, $\bar{y}(t, r, \psi)$, to a measurement. To illustrate the uncertainty of the model fits, we sample 1000 sets of model parameters for each individual from the posterior distribution (Eq S4.15) and simulate the PKPD model output for each parameter set, e.g. for individual 1, we sample from $p(\psi_{1\,1}|\mathcal{D}_1) = \int \mathrm{d}\theta\, \mathrm{d}\psi_{1\,2} \dots \mathrm{d}\psi_{1\,60}\, p(\theta, \Psi_1|\mathcal{D}_1)$ and solve $\bar{y}(t, r_{1\,1}, \psi_{1\,1})$ for each sample of $\psi_{1\,1}$. The median PKPD model output for each fit is illustrated by a black scatter point. The uncertainty of the fit is illustrated using an error bar, representing the 5th to 95th percentile interval of the output distribution. Model outputs identical to measurements lie on the diagonal line (blue). The right panel shows the quality of the fits on a population level. This time PKPD model outputs are simulated using the typical parameters in

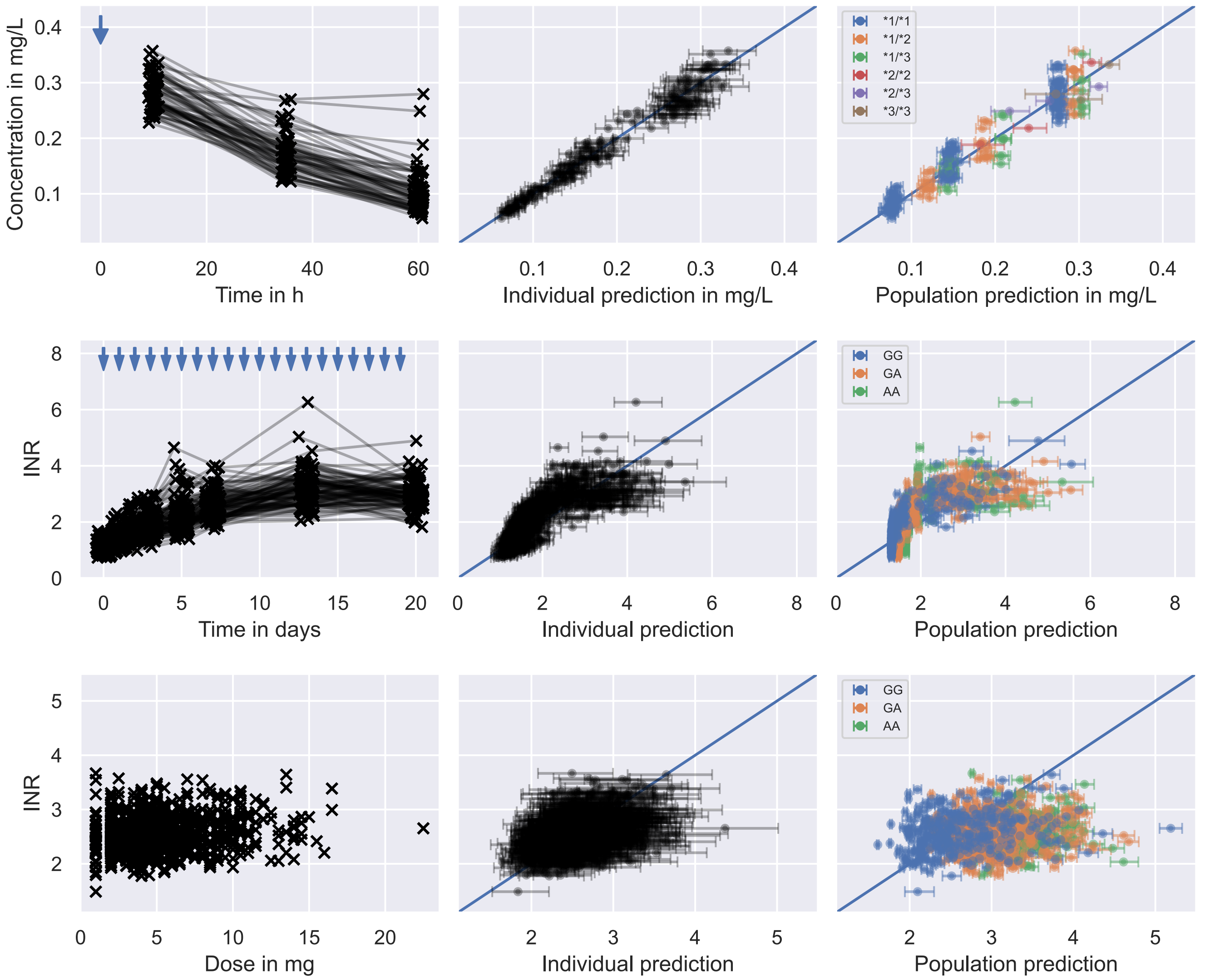


**Figure S4.9. Goodness of fit.** The figure shows the goodness-of-fit to the trial data: phase I (top row); phase II (middle row); phase III (bottom row). In the columns shows the measurements used for the calibration of the model. Nominal administration times are indicated using blue arrows. The middle columns show the individual-level fits to the measurements. Each scatter point represents the fit of the PKPD model, $\bar{y}(t, r, \psi)$, to a measurement. We sample 1000 sets of model parameters for each individual from the posterior distribution and simulate the PKPD model output for each parameter set, e.g. for individual 1 in trial I, we sample from $p(\psi_{1\,1}|\mathcal{D}_1) = \int \mathrm{d}\theta\, \mathrm{d}\psi_{1\,2} \ldots \mathrm{d}\psi_{1\,60}\, p(\theta, \Psi_1|\mathcal{D}_1)$ and solve $\bar{y}(t, r_{1\,1}, \psi_{1\,1})$ for each sample of $\psi_{1\,1}$. The median PKPD model output for each fit is illustrated by the black scatter point. The error bars illustrate the 5th to 95th percentile interval of the output distribution. Model outputs identical to measurements lie on the diagonal line (blue). The right columns show the population-level fits to the measurements. We simulate the PKPD model outputs using 1000 sets of the typical parameters sampled from the posterior, e.g. for individual 1 in trial 1, we sample 1000 sets of population parameters from the posterior distribution, $p(\theta|\mathcal{D}_1)$, and use Eq S4.10 and the the individual's covariates, $\chi_{1\,1}$, to calculate the typical parameters. For each set of typical parameters, we simulate the PKPD model output. The median model output is illustrated by the scatter point and the 5th to 95th percentile interval are illustrated by the error bars.

the population, e.g. for individual 1, we sample 1000 sets of population parameters from the posterior distribution, $p(\theta|\mathcal{D}_1)$, and use Eq S4.10 and the the individual's covariates, $\chi_{1\,1}$, to calculate the typical parameters, $(\mu_{k_e}, \mu_v)$. For each set of typical parameters, we simulate the PKPD model output. The median model output and percentile interval are determined analogously to the middle panel. Individuals with different CYP2C9 genotypes are highlighted in different colours: *1*1 (blue); *1*2 (orange); *1*3 (green); *2*2 (red); *2*3 (purple); *3*3 (brown). The age differences between individuals are not highlighted in the figure, but taken into account in the computation of the typical parameters.

The middle panel shows that the individual-level residuals are symmetrically distributed around the line of identity (blue), giving no indication of model misspecification. The residual error also seems to increase with the model output, which supports the lognormal error model choice. Similarly the residuals with respect to the typical treatment response are symmetrically distributed around the line of identity (see right panel), showing that the typical values of the model parameters are a good representation of the typical treatment response in the population.

##### S4.4.2 Calibration to trial phase II data

After the calibration to the phase I data, we continue the model parameter calibration using the data from trial phase II. The data contains INR measurements of 100 individuals (see middle panel in Fig 5). Denoting the data as $\mathcal{D}_2 = (\mathcal{D}_{2\,1}, \ldots, \mathcal{D}_{2\,100})$, where $\mathcal{D}_{2i}$ denotes the data of individual $i$, we define the hierarchical posterior for the population model parameters analogously to Eq S4.15 using Bayes rule

$$\log p(\theta, \Psi_2 | \mathcal{D}_1, \mathcal{D}_2) = \sum_i \log p(\mathcal{D}_{2i} | \psi_{2i}) + \sum_i \log p(\psi_{2i} | \theta, \chi_{2i}) + \log p(\theta | \mathcal{D}_1) + \text{constant}. \quad \text{(S4.17)}$$

$\Psi_2 = (\psi_{2\,1}, \ldots, \psi_{2\,100})$ denotes the individual-level parameters of the individuals in the trial. The likelihood of the individal-level parameters is defined using the measurement distribution of the INR measurements defined in Section S4.3. The scale parameter of the lognormal measurement distribution, $\sigma$, is pooled across individuals.

The prior distribution of the model parameters is informed by the inference results from the previous section. We set the marginal prior distributions of the PK-related model parameters to the marginal posterior distributions from the phase I calibration in Fig S4.8. For computational efficiency, we approximate the inferred distributions (blue) using parametrised distributions (dashed lines). We calibrate the parameters of the distributions manually, using empirical means and standard deviations of the samples as a staring point. The resulting prior distributions for the PK parameters are

$$\begin{aligned}
\log k_{*1} &\sim \mathcal{N}(-3.682, 0.028) \\
\sigma_{k_e} &\sim \mathcal{N}(0.119, 0.020) \\
\frac{k_{*1} - k_{*2}}{k_{*2}} &\sim \mathcal{N}(0.565, 0.063) \\
\frac{k_{*1} - k_{*3}}{k_{*2}} &\sim \text{Beta}(30, 6) \\
r_{age} &\sim \mathcal{N}(0.00546, 0.0031) \\
\log \mu_v &\sim \mathcal{N}(2.662, 0.020) \\
\sigma_v &\sim \text{LN}(-2.31, 0.16).
\end{aligned}$$

The prior information of the remaining parameters is assumed to be minimal. We define weakly informative marginal distributions only to ensure that parameters assume values in meaningful biological ranges

$$\begin{aligned}
\log \bar{y}_{0,G} &\sim \mathcal{N}(0, 0.5) \\
\sigma_{\bar{y}_0} &\sim \text{LN}(-1, 2) \\
\bar{y}_{0,A}/\bar{y}_{0,G} &\sim \text{LN}(0, 1) \\
\log c_G &\sim \mathcal{N}(1.41, 0.5) \\
\sigma_{c50} &\sim \text{LN}(-1, 0.5) \\
\frac{c_G - c_A}{c_G} &\sim \mathcal{U}(0, 1) \\
\mu_{k_1} &\sim \mathcal{N}(0.2, 0.05) \\
\mu_{k_2} &\sim \mathcal{N}(0.02, 0.005) \\
\mu_{\sigma} &\sim \text{LN}(-1, 0.3).
\end{aligned}$$

We fix the potency of warfarin to $\gamma = 1.15$, following (Hamberg et al., 2010).

We infer the posterior following the same approach as in the previous section (see S4.4.1), using NUTS and 1500 iterations. All marginal posteriors have $\hat{R} \leq 1.01$. We show the inferred posterior distribution in Fig S4.10. The results of the inference, as well as data and code to reproduce the results are hosted on GitHub (https://github.com/DavAug/mipd-warfarin).

Fig S4.10 shows the results of the calibration. The marginal prior distributions are illustrated in black (solid lines) and the posterior distributions are illustrated in blue. Approximations of the posterior distributions, used in Section S4.4.3, are illustrated in black (dashed lines).

The goodness-of-fit is visualised in the middle row of Fig S4.9. The residuals of the individual-level fits in the middle column indicate no sign of model misspecification. The asymmetric distribution of the residuals in the right column suggests however that the typical parameter values do not represent the typical treatment response. In particular, predicted typical INRs in the therapeutic window appear to underestimate the measured INRs, as indicated by the scew of the residual distribution towards values above the line of identity for INRs between 2 and 3. This mismatch between typical model parameters and typical treatment reponses does not necessarily imply a misspecification of the population model, but may simply be the consequence of the nonlinear relationship between the PKPD model output and the model parameters.

##### S4.4.3 Calibration to trial phase III data

We conclude the inference by calibrating the model parameters to the data from trial phase III. The data contains INR measurements of 1000 individuals under maintenance treatment (see right panel in Fig 5). Denoting the data with $\mathcal{D}_3 = (\mathcal{D}_{3\,1}, \ldots, \mathcal{D}_{3\,1000})$, where $\mathcal{D}_{3i}$ denotes the data of individual $i$, we define the hierarchical posterior for the population model parameters using Bayes rule

$$\log p(\theta, \Psi_3 | \mathcal{D}_1, \mathcal{D}_2, \mathcal{D}_3) = \sum_i \log p(\mathcal{D}_{3i} | \psi_{3i}) + \sum_i \log p(\psi_{3i} | \theta, \chi_{3i}) + \log p(\theta | \mathcal{D}_1, \mathcal{D}_2) + \text{constant}. \tag{S4.18}$$

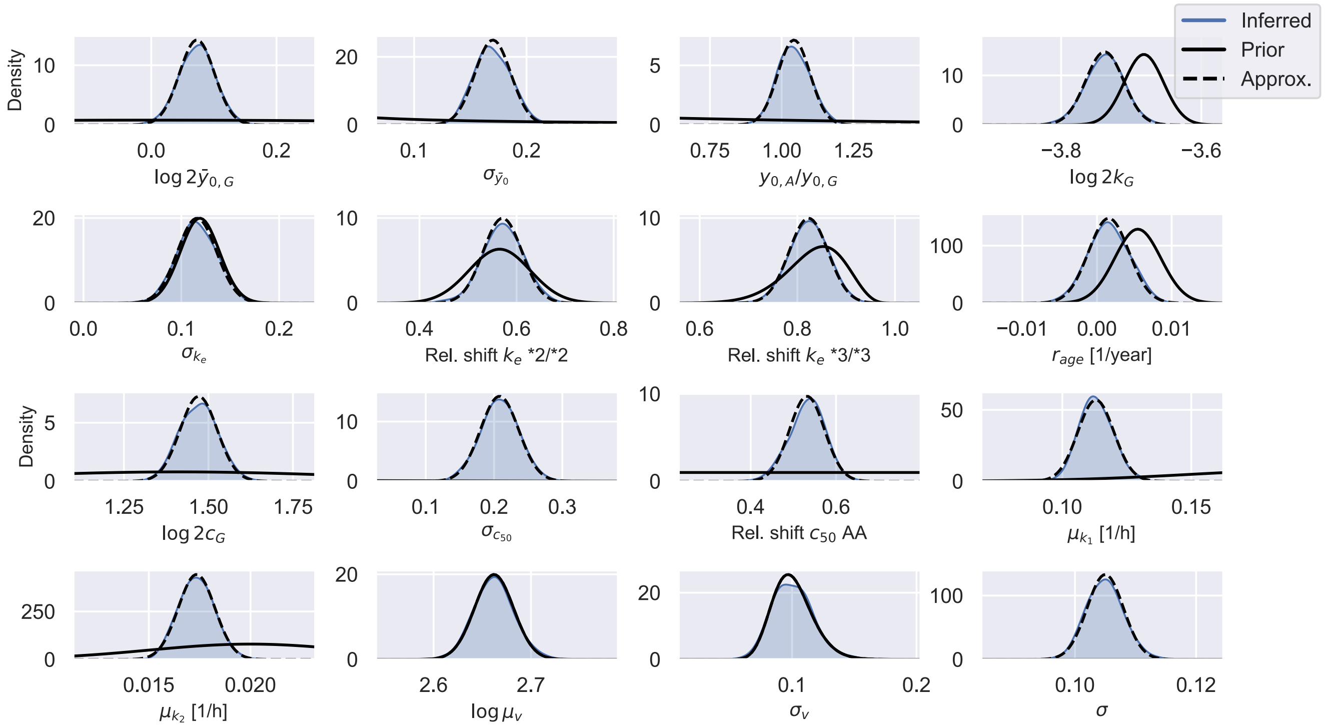


**Figure S4.10. Marginal posteriors after calibration to trial phase II data.** The figure shows the marginal posteriors of the model parameters after calibrating the modified Hamberg et al model to the phase II trial data. The priors are illustrated in black (solid lines) and the inferred posteriors are illustrated in blue using kernel density estimates. The approximations of the posteriors are illustrated by black dashed lines.

$\Psi_3 = (\psi_{3\,1}, \ldots, \psi_{3\,1000})$ denotes the individual-level parameters of the individuals in the trial. For efficiency, we define the INR measurement distribution using the analytically-computed steady state of the PKPD model.

In steady state, all rate of change equations of the PKPD model in Section S4.2 vanish. For a constant dose rate, $\bar{r} =$ constant, we can use this to derive an analytic expression for the steady state of the warfarin concentration from Eq S4.8

$$c_{ss} = \frac{\bar{r}}{v k_e}. \tag{S4.19}$$

Similarly, we can derive a steady state expression for the INR from Eqs S4.6 and S4.7

$$\bar{y}_{ss} = \bar{y}_0 + \bar{y}_{max}\, \kappa \frac{c_{ss}^{\gamma}}{c_{50}^{\gamma} + c_{ss}^{\gamma}}. \tag{S4.20}$$

Thus, we can efficiently simulate the INR under maintenance treatment using these expressions, assuming a dose rate that is constant in time.

However, the daily warfarin doses administered during the trial do not give rise to constant dose rates. The steady state expressions can therefore only approximate the treatment response. The exact warfarin concentrations fluctuate around the steady state value computed with the average daily dose rate, $\bar{r} = d^*/24\,$h. Despite these fluctuations in the warfarin concentration, the approximation error for the INR simulations can be neglected for the inference, as the treatment response delay dampens the effect of the fluctuations on the INR (see Fig S4.1), resulting in an effectively constant INR value under maintenance treatment.

For the inference, we fix the scale parameters of the measurement distribution, the elimination rate distribution, the EC50 distribution and the volume of distribution distirbution to the typical values from the phase II calibration: $\mu_\sigma = 0.105$; $\sigma_{k_e} = 0.116$; $\sigma_{c_{50}} = 0.208$; $\sigma_v = 0.101$ (see Fig S4.10). This is because snapshot measurements make it difficult to distinguish contributions from inter-individual variability and measurement noise to the observed treatment response variability (Augustin et al., 2023). The maintenance treatment data also makes it difficult to distinguish variability in the treatment response from variability in the baseline INR. We therefore also fix the baseline INR parameters to the typical values from Fig S4.10: $\log 2\bar{y}_{0,G} = 0.073$; $\sigma_{\bar{y}_0} = 0.170$; $\bar{y}_{0,A}/\bar{y}_{0,G} = 1.043$. This leaves 7 parameters for the calibration: $(k_{*1}, k_{*2}, k_{*3}, r_{age}, c_G, c_A, \mu_v)$.

The prior distribution of the model parameters is set to the marginal posterior distributions from the phase II calibration in Fig S4.10

$$
\begin{aligned}
\log 2k_{*1} &\sim \mathcal{N}(-3.787, 0.027) \\
\frac{k_{*1} - k_{*2}}{k_{*2}} &\sim \mathcal{N}(0.571, 0.040) \\
\frac{k_{*1} - k_{*3}}{k_{*2}} &\sim \mathcal{N}(0.823, 0.040) \\
r_{age} &\sim \mathcal{N}(0.00157, 0.0027) \\
\log 2c_G &\sim \mathcal{N}(1.471, 0.055) \\
\frac{c_G - c_A}{c_G} &\sim \mathcal{N}(0.532, 0.041) \\
\log \mu_v &\sim \mathcal{N}(2.662, 0.020).
\end{aligned}
$$

We infer the posterior following the same approach as in the previous sections (see S4.4.1), using NUTS and 1500 iterations. All marginal posteriors have $\hat{R} \leq 1.01$. We show the inferred posterior distribution in Fig S4.11. The results of the inference, as well as data and code to reproduce the results are hosted on GitHub (https://github.com/DavAug/mipd-warfarin).

The figure shows the final result of the model calibration. The marginal prior distributions are illustrated in black (solid lines) and the posterior distributions are illustrated in blue. Posteriors not updated since the calibration to the trial phase II data are labelled with (VCT II).

The goodness-of-fit is visualised in the bottom panel of Fig S4.9. Similar to the inference results from trial phase II (see middle row), the residuals of the individual-level fits in the middle column indicate no sign of model misspecification, while the asymmetric distribution of the residuals in the right column suggests that the typical parameter values do not represent the typical treatment response.

##### S4.4.4 Prediction of parameter priors for MIPD

We predict individualised prior distributions used in Eq S4.13 for MIPD (see Section S4.3) using the calibrated model. As illustrated in Fig S4.11, the result of the calibration is a distribution of population-level parameters, $p(\theta|\mathcal{D}_1, \mathcal{D}_2, \mathcal{D}_3)$, compatible with the clinical trial data. The spread of the distribution indicates that there is residual uncertainty about the model parameters.

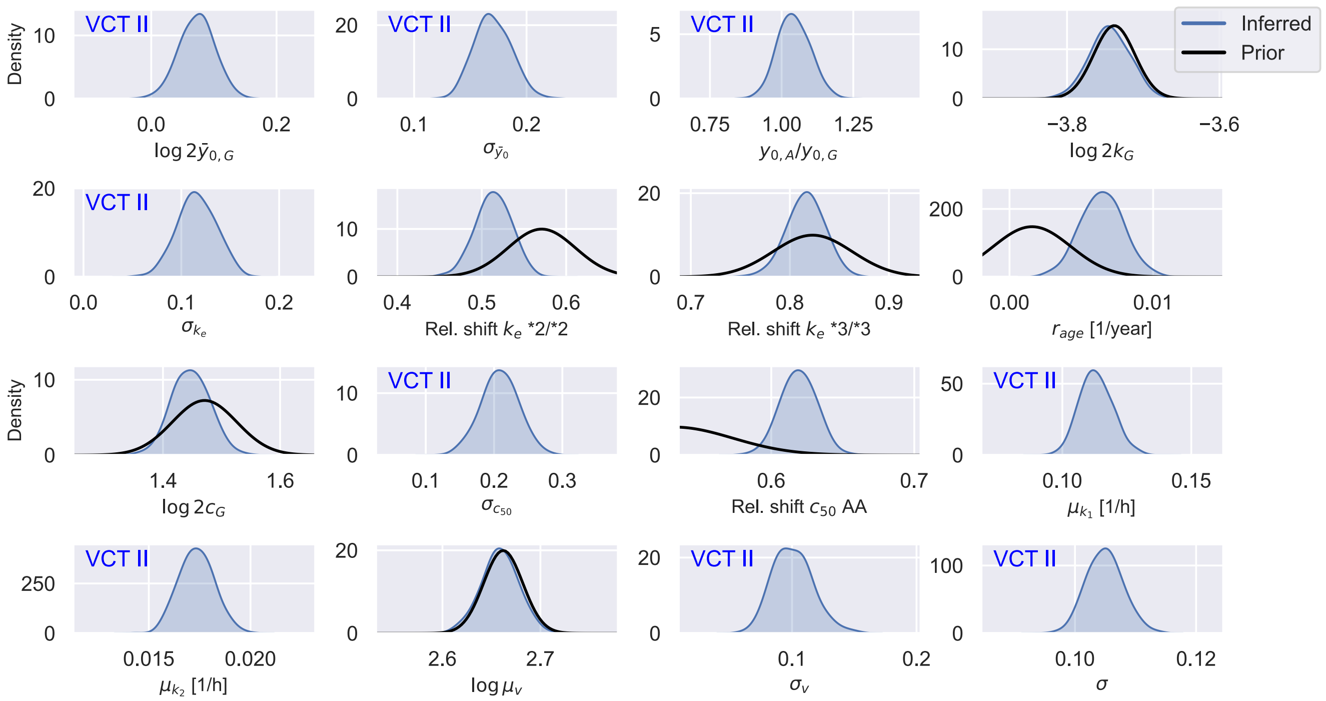


**Figure S4.11. Marginal posteriors after calibration to trial phase III data.** The figure shows the marginal posteriors of the model parameters after calibrating the modified Hamberg et al model to the phase III trial data. The priors are illustrated in black (solid lines) and the inferred posteriors are illustrated in blue using kernel density estimates. The approximations of the posteriors are illustrated by black dashed lines.

To account for this residual uncertainty, we average the population model, $p(\psi|\theta, \chi)$, over the posterior distribution

$$p(\psi|\chi, \mathcal{D}_1, \mathcal{D}_2, \mathcal{D}_3) = \mathbb{E}_{\theta|\mathcal{D}_1,\mathcal{D}_2,\mathcal{D}_3}\Big[p(\psi|\theta, \chi)\Big]. \qquad \text{(S4.21)}$$

In the Bayesian inference literature, $p(\psi|\chi, \mathcal{D}_1, \mathcal{D}_2, \mathcal{D}_3)$, may be referred to as the posterior predictive distribution. An example individualised prior for an individual with the covariates $\chi = (GG, {*1}{*1}, 71)$ is illustrated in Fig S4.12.

The figure shows the individualised prior constructed from 1000 samples of $p(\psi|\chi, \mathcal{D}_1, \mathcal{D}_2, \mathcal{D}_3)$ (blue). Each $\psi$ is generated by first sampling a set of population parameters from $p(\theta|\mathcal{D}_1, \mathcal{D}_2, \mathcal{D}_3)$ and subsequently sampling the $\psi$ from $p(\psi|\theta, \chi)$ using the sampled $\theta$ and the covariates of the individual. For more efficient inference during the MIPD trial, we again approximate the probability densities using parametrised distributions. All marginal distributions are well approximated using Gaussian distributions (black). The prior distributions of all individuals in the MIPD trial, as well as data and code to reproduce the results are hosted on GitHub (https://github.com/DavAug/mipd-warfarin).

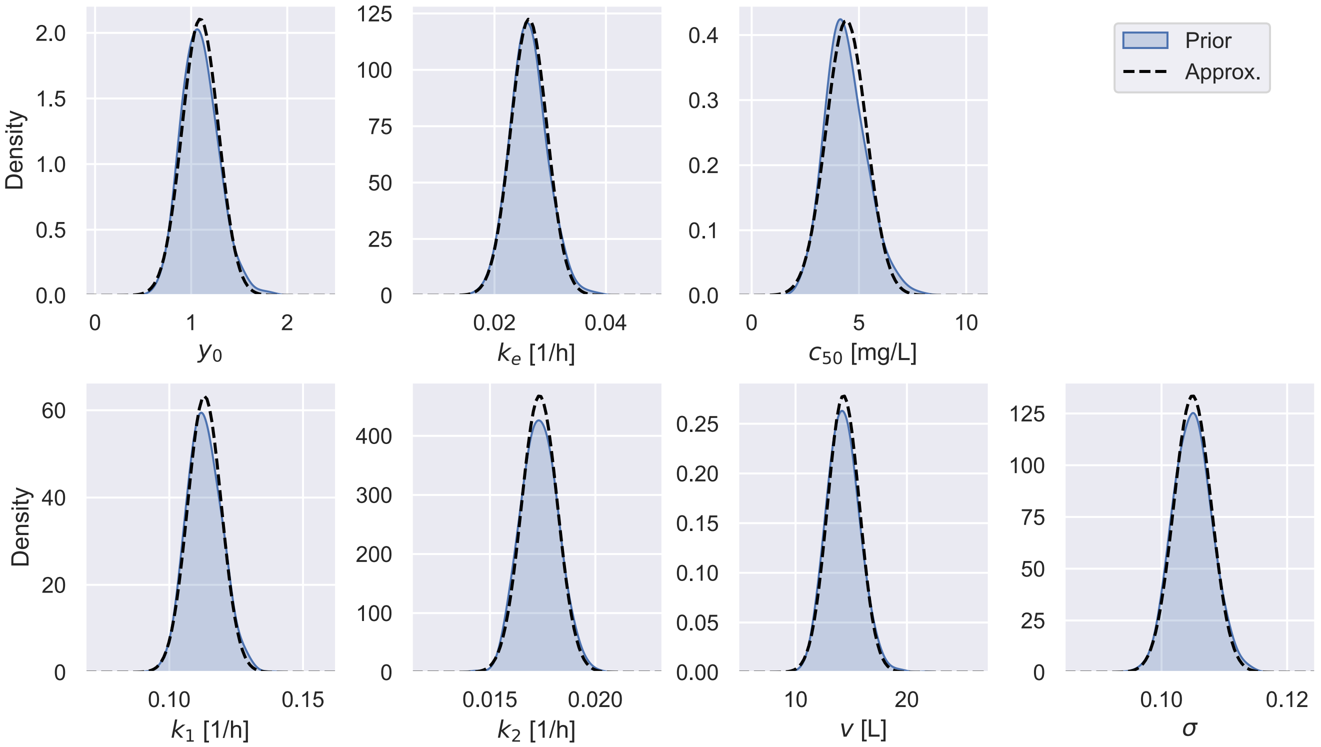


**Figure S4.12. Individualised prior distribution.** The figure shows the prior distribution used during the MIPD trial for individual with the covariates $\chi = (GG, *1*1, 71)$. The inferred prior distributions, $p(\psi|\chi, \mathcal{D}_1, \mathcal{D}_2, \mathcal{D}_3)$, are visualised in blue. Gaussian approximations of the prior distributions are illustrated in black.

#### Algorithm S1

**Simulation of clinical trial phase I: Warfarin concentration after single dose.**

**Input :** 1. Mechanistic model: $\bar{y}(t, r, \psi)$;
2. Population model: $p(\psi|\theta, \chi)$;
3. Execution model: $p(\Delta t|\tau)$;
4. Measurement model: $p(y|\bar{y}, \psi)$;
5. Covariates of virtual patients: $\chi_1, \ldots, \chi_{60}$;
6. Nominal dosing regimen: $r(t) = d\,\Theta(t)\,\Theta(\Delta - t)/\Delta$ with $d = 10\,\text{mg}$, $\Delta = 0.01\,\text{h}$;
7. Nominal measurement times: $t_1 = 10\,\text{h}$, $t_2 = 35\,\text{h}$, $t_3 = 60\,\text{h}$

**Output :** Virtual clinical trial data $\mathcal{D}$.

$\mathcal{D}$ = [ ]
**for** $i \leftarrow 1$ **to** $60$ **do**
    $\psi_i \sim p(\psi|\theta, \chi_i)$ // Sample parameters of virtual patient
    $\Delta t_0 \sim p(\Delta t|\tau)$ // Sample delay of administration
    $r' = d\,\Theta(t - \Delta t_0)\,\Theta(\Delta t_0 + \Delta - t)/\Delta$ // Update dosing regimen
    $\mathcal{D}_i$ = [ ]
    **for** $j \leftarrow 1$ **to** $3$ **do**
        $\Delta t_j \sim p(\Delta t|\tau)$ // Sample delay of measurement
        $\bar{y}_{ij} = \bar{y}(t_j + \Delta t_j, r', \psi_i)$ // Simulate concentration
        $y_{ij} \sim p(y|\bar{y}_{ij}, \psi_i)$ // Sample measurement
        $\mathcal{D}_i$.append$((y_{ij}, t_j))$ // Store measurement with nominal time
    **end for**
    $\mathcal{D}$.append$((\mathcal{D}_i, \chi_i, r))$ // Store data with nominal dosing regimen
**end for**
**return** $\mathcal{D}$

#### *Algorithm S2*

**Simulation of clinical trial phase II: INR time course during 3 weeks of treatment.**

**Input :** 1. Mechanistic model: $\bar{y}(t, r, \psi)$;
2. Population model: $p(\psi|\theta, \chi)$;
3. Inter-occasion model: $p(\eta|t)$;
4. Execution model: $p(\Delta t|\tau)$;
5. Measurement model: $p(y|\bar{y}, \psi)$;
6. Covariates of virtual patients: $\chi_1, \ldots, \chi_{100}$;
7. Nominal regimen: $r_i(t) = \sum_{j=1}^{20} d_{ij}\, \Theta(t - \tilde{t}_j)\, \Theta(\tilde{t}_j + \Delta - t)/\Delta$ with $\tilde{t}_j = 24(j-1)\,\mathrm{h}$;
8. Nominal measurement times: $t_1 = 0\,\mathrm{h}$, $t_2 = 24\,\mathrm{h}$, $t_3 = 2 \times 24\,\mathrm{h}$, $t_4 = 3 \times 24\,\mathrm{h}$, $t_5 = 5 \times 24\,\mathrm{h}$, $t_6 = 7 \times 24\,\mathrm{h}$, $t_7 = 13 \times 24\,\mathrm{h}$, $t_8 = 20 \times 24\,\mathrm{h}$

**Output :** Virtual clinical trial data $\mathcal{D}$.

1 $\mathcal{D}$ = [ ]
2 *measurement_times* $= (t_1, t_2, t_3, t_4, t_5, t_6, t_7, t_8)$
3 *dose_adjustment_times* $= (t_4, t_5, t_6, t_7)$
4 **for** $i \leftarrow 1$ **to** $100$ **do**
5     $\psi_i \sim p(\psi|\theta, \chi_i)$ // Sample parameters of virtual patient
6     $r'_i = r_i$ // Initialise dosing regimen
7     $\mathcal{D}_i$ = [ ]
8     **for** $j \leftarrow 1$ **to** $20$ **do**
9         $\eta_j \sim p(\eta|t_j)$ // Sample inter-occasion variability
10         $\psi_{ij} = \psi_i\, \eta_j$ // Update model parameters
11         $\Delta t_j \sim p(\Delta t|\tau)$ // Sample measurement and administration delay
12         **if** $t_j \in$ *measurement_times* **then**
13             $\bar{y}_{ij} = \bar{y}(\Delta t_j, r'_i, \psi_{ij})$ // Simulate INR starting from $t_j$ (see l. 25)
14             $y_{ij} \sim p(y|\bar{y}_{ij}, \psi_{ij})$ // Sample measurement
15             $\mathcal{D}_i$.append$((y_{ij}, t_j))$ // Store measurement with nominal time
16         **end if**
17         **if** $j \leq 3$ **then**
18             $d'_{ij} = 10 - 2.5(j-1)$ // Get dose 1, 2 or 3
19         **else if** $\tilde{t}_j \in$ *dose_adjustment_times* **then**
20             $d'_{ij}$ = updateDose$(y_{ij})$ // Adjust dose using heuristic
21         **end if**
22         $r_i = r_i$.update$(d_{ij} = d'_{ij})$ // Update nominal dosing regimen
23         $r'_i = d'_{ij}\, \Theta(t - \Delta t_j)\, \Theta(\Delta t_j + \Delta - t)/\Delta$ // Define actual dosing regimen
24         $\psi_0 = \bar{y}(t = 24, r'_i, \psi_{ij})$.getState() // Simulate state of model at next day
25         $\psi_i = \psi_i$.setState$(\psi_0)$ // Update state to shift reference time to $t_j$
26     **end for**
27     $\mathcal{D}$.append$((\mathcal{D}_i, \chi_i, r))$ // Store data with nominal dosing regimen
28 **end for**
29 **return** $\mathcal{D}$

##### *Algorithm S3*

**Simulation of clinical trial phase III: Maintenance treatment.**

**Input :** 1. Mechanistic model: $\bar{y}(t, r, \psi)$;
2. Population model: $p(\psi|\theta, \chi)$;
3. Inter-occasion model: $p(\eta|t)$;
4. Execution model: $p(\Delta t|\tau)$;
5. Measurement model: $p(y|\bar{y}, \psi)$;
6. Covariates of virtual patients: $\chi_1, \ldots, \chi_{1000}$;
7. Nominal regimen: $r_i(t) = \sum_{j=1}^{55} d_{ij}\, \Theta(t - \tilde{t}_j)\, \Theta(\tilde{t}_j + \Delta - t)/\Delta$ with $\tilde{t}_j = 24(j-1)\,\mathrm{h}$;
8. Nominal measurement times: $\{t_j = 24(j-1)\,\mathrm{h} \quad \text{for} \quad j = 1, \ldots, 55\}$

**Output :** Virtual clinical trial data $\mathcal{D}$.

1 $\mathcal{D}$ = [ ]
2 *measurement_times* $= (t_1, t_2, t_3, t_4, t_5, t_6, t_7, t_8, t_9, t_{10}, t_{11}, t_{12})$
3 *dose_adjustment_times* $= (t_4, t_5, t_6, t_7, t_8, t_9)$
4 **for** $i \leftarrow 1$ **to** $1000$ **do**
5     $\psi_i \sim p(\psi|\theta, \chi_i)$ `// Sample parameters of virtual patient`
6     $r'_i = r_i$ `// Initialise dosing regimen`
7     $\mathcal{D}_i$ = [ ]
8     **for** $j \leftarrow 1$ **to** $55$ **do**
9         $\eta_j \sim p(\eta|t_j)$ `// Sample inter-occasion variability`
10         $\psi_{ij} = \psi_i\, \eta_j$ `// Update model parameters`
11         $\Delta t_j \sim p(\Delta t|\tau)$ `// Sample measurement and administration delay`
12         **if** $t_j \in$ *measurement_times* **then**
13             $\bar{y}_{ij} = \bar{y}(\Delta t_j, r'_i, \psi_{ij})$ `// Simulate INR starting from` $t_j$ `(see l. 25)`
14             $y_{ij} \sim p(y|\bar{y}_{ij}, \psi_{ij})$ `// Sample measurement`
15         **end if**
16         **if** $j \leq 3$ **then**
17             $d'_{ij} = 10 - 2.5(j-1)$ `// Get dose 1, 2 or 3`
18         **else if** $\tilde{t}_j \in$ *dose_adjustment_times* **then**
19             $d'_{ij}$ = updateDose$(y_{ij})$ `// Adjust dose using heuristic`
20         **end if**
21         $r_i = r_i$.update$(d_{ij} = d'_{ij})$ `// Update nominal dosing regimen`
22         $r'_i = d'_{ij}\, \Theta(t - \Delta t_j)\, \Theta(\Delta t_j + \Delta - t)/\Delta$ `// Define actual dosing regimen`
23         $\psi_0 = \bar{y}(t = 24, r'_i, \psi_{ij})$.getState() `// Simulate state of model at next day`
24         $\psi_i = \psi_i$.setState$(\psi_0)$ `// Update model state to shift reference time to` $t_j$
25     **end for**
26     $\mathcal{D}_i$.append$((y_{i\,12}, t_{12}))$ `// Store final measurement with nominal time`
27     $\mathcal{D}$.append$((\mathcal{D}_i, \chi_i, r))$ `// Store data with nominal dosing regimen`
28 **end for**
29 **return** $\mathcal{D}$

##### *Algorithm S4*

**Simulation of MIPD trial: INR time course during 19 days of treatment.**

**Input :** 1. MIPD model: $d(\mathcal{D}, \chi, r)$;
2. Mechanistic model: $\bar{y}(t, r, \psi)$;
3. Population model: $p(\psi|\theta, \chi)$;
4. Inter-occasion model: $p(\eta|t)$;
5. Execution model: $p(\Delta t|\tau)$;
6. Measurement model: $p(y|\bar{y}, \psi)$;
7. Covariates of virtual patients: $\chi_1, \ldots, \chi_{1000}$;
8. Nominal regimen: $r_i(t) = \sum_{j=1}^{20} d_{ij}\, \Theta(t - \tilde{t}_j)\, \Theta(\tilde{t}_j + \Delta - t)/\Delta$ with $\tilde{t}_j = 24(j-1)\,\mathrm{h}$

**Output :** Virtual clinical trial data $\mathcal{D}$.

$\mathcal{D}$ = [ ]<br>
**for** $i \leftarrow 1$ **to** $1000$ **do**<br>
    $\psi_i \sim p(\psi|\theta, \chi_i)$ `// Sample parameters of virtual patient`<br>
    $r'_i = r_i$ `// Initialise dosing regimen`<br>
    $\mathcal{D}_i$ = [ ]<br>
    **for** $j \leftarrow 1$ **to** $20$ **do**<br>
        $\eta_j \sim p(\eta|t_j)$ `// Sample inter-occasion variability`<br>
        $\psi_{ij} = \psi_i\, \eta_j$ `// Update model parameters`<br>
        $\Delta t_j \sim p(\Delta t|\tau)$ `// Sample measurement and administration delay`<br>
<br>
        $\bar{y}_{ij} = \bar{y}(\Delta t_j, r'_i, \psi_{ij})$ `// Simulate INR starting from` $t_j$ `(see l. 25)`<br>
        $y_{ij} \sim p(y|\bar{y}_{ij}, \psi_{ij})$ `// Sample measurement`<br>
        $\mathcal{D}_i$.append$((y_{ij}, t_j))$ `// Store measurement with nominal time`<br>
<br>
        $d'_{ij}$ = $d(\mathcal{D}_i, \chi_i, r_i)$ `// Predict next dose using MIPD model`<br>
        $r_i = r_i$.update$(d_{ij} = d'_{ij})$ `// Update nominal dosing regimen`<br>
        $r'_i = d'_{ij}\, \Theta(t - \Delta t_j)\, \Theta(\Delta t_j + \Delta - t)/\Delta$ `// Define actual dosing regimen`<br>
<br>
        $\psi_0 = \bar{y}(t = 24, r'_i, \psi_{ij})$.getState() `// Simulate state of model at next day`<br>
        $\psi_i = \psi_i$.setState$(\psi_0)$ `// Update state to shift reference time to` $t_j$<br>
    **end for**<br>
    $\mathcal{D}$.append$((\mathcal{D}_i, \chi_i, r))$ `// Store data with nominal dosing regimen`<br>
**end for**<br>
**return** $\mathcal{D}$

#### Table S1.

**Demographics of virtual clinical trials.** The age distribution of each cohort is represented by the median and the 5th to 95th percentile interval.

| | Phase I | Phase II | Phase III |
|---|---|---|---|
| Number of patients | 60 | 100 | 1000 |
| VKORC1 GG | N/A | 36 % | 36.5 % |
| VKORC1 GA | N/A | 49 % | 48.5 % |
| VKORC1 AA | N/A | 15 % | 15 % |
| CYPC29 *1*1 | 62 % | 63 % | 66.1 % |
| CYPC29 *1*2 | 20 % | 19 % | 18.4 % |
| CYPC29 *1*3 | 13 % | 13 % | 12.3 % |
| CYPC29 *2*2 | 2 % | 2 % | 1.4 % |
| CYPC29 *2*3 | 2 % | 2 % | 1.2 % |
| CYPC29 *3*3 | 1 % | 1 % | 0.6 % |
| Age | 69 (58 - 79) | 65 (56 - 76) | 50 (40 - 64) |
